# A Multi-omics Data Analysis Workflow Packaged as a FAIR Digital Object

**DOI:** 10.1101/2023.06.07.543986

**Authors:** Anna Niehues, Casper de Visser, Fiona A. Hagenbeek, Purva Kulkarni, René Pool, Naama Karu, Alida Kindt, Gurnoor Singh, Robert R. J. M. Vermeiren, Dorret I. Boomsma, Jenny van Dongen, Peter A. C. ‘t Hoen, Alain J. van Gool

## Abstract

**Background:** Applying good data management and FAIR data principles (Findable, Accessible, Interoperable, and Reusable) in research projects can help disentangle knowledge discovery, study result reproducibility, and data reuse in future studies. Based on the concepts of the original FAIR principles for research data, FAIR principles for research software were recently proposed. FAIR Digital Objects enable discovery and reuse of Research Objects, including computational workflows for both humans and machines. Practical examples can help promote the adoption of FAIR practices for computational workflows in the research community. We developed a multi-omics data analysis workflow implementing FAIR practices to share it as a FAIR Digital Object.

**Findings:** We conducted a case study investigating shared patterns between multi-omics data and childhood externalizing behavior. The analysis workflow was implemented as a modular pipeline in the workflow manager Nextflow, including containers with software dependencies. We adhered to software development practices like version control, documentation, and licensing. Finally, the workflow was described with rich semantic metadata, packaged as a Research Object Crate, and shared via WorkflowHub.

**Conclusions:** Along with the packaged multi-omics data analysis workflow, we share our experiences adopting various FAIR practices and creating a FAIR Digital Object. We hope our experiences can help other researchers who develop omics data analysis workflows to turn FAIR principles into practice.

Key Points

- The FAIR4RS principles provide guidelines to enhance the discovery and reuse of research software.
- FAIR Digital Objects support Findability, Accessibility, Interoperability, and Reusability by both humans and machines.
- We here demonstrate the implementation multi-omics data analysis workflow and share it as a FAIR Digital Object.

## Background

The FAIR principles for research data [1] were proposed to guide researchers to create research data that is findable, accessible, interoperable, and reusable (FAIR). Since these guidelines aim to enable researchers handling and navigating through the rapidly increasing amounts of data, special emphasis was put on concepts to make data not only usable by humans but also machine-actionable. In the past years, various standards [2, 3] and implementations [4, 5, 6, 7] of the FAIR principles have been introduced, and it has been demonstrated that FAIR data is beneficial to research and patients [8, 9, 10]. Reuse of research data and reproducibility of research results [11] are facilitated by good data provenance and this requires not only the data but also the data processing and analysis workflows to be FAIR. Consequently, guidelines and practices for FAIR research software have been proposed [12, 13, 14] (see Table 1) and the special case of computational workflows has been discussed [15, 16]. These guidelines aim to increase reproducibility not only at the experimental level but also at the data analysis level. It has been shown that the availability of data and code alone is not sufficient. They both need to be provided in an open and interoperable format and described by metadata [17].

**Table 1.**
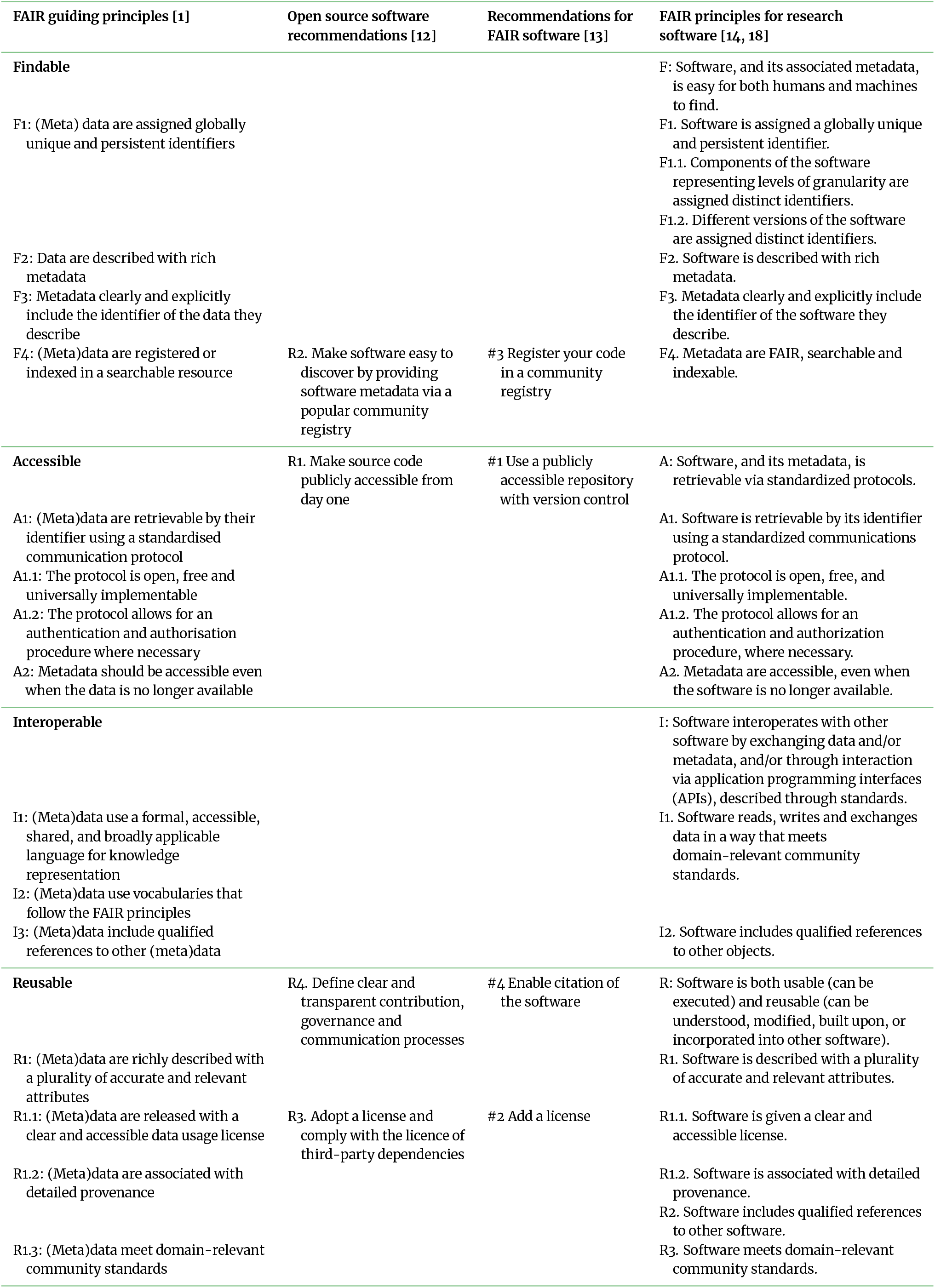
Overview of recommended FAIR practices for research data and software.

Several practices recommended for research software development originate from general software engineering practices [12, 15, 19] which include version control, documentation, and licensing. Version control of source code facilitates collaborative development and monitoring changes [13]. Additionally, making the code publicly available on dedicated software repositories that support version control such as GitHub [20], GitLab [21], or BitBucket [22] contributes to findability [23], accessibility [12], and reusability [13]. The documentation of research software includes multiple levels. First, a comprehensive end user documentation and usage examples enable reusability by other researchers [17, 24, 25, 23]. It should also include the documentation of workflow parameters [17, 16]. Second, source code documentation enables other developers to understand and build upon the software [17]. Documentation of code changes via a version control system helps document the development process [24, 19] and documentation of dependencies are prerequisite for software interoperability [23]. Adding a clear and machine-readable [16] license is essential to allow for software reuse. It is recommended to choose a widely used and preferably open source license that is compatible with licenses of the dependencies [12, 24, 19, 23, 13, 14, 18]. Examples of open-source licenses with few restrictions are the Apache License 2.0 [26] and the MIT License [27].

There are differences between research software that implements a specific method as a standalone tool or a software library and complex analysis workflows [16]. Computational analysis workflows can comprise numerous steps that are integrated into pipelines [16] and are often developed in a specific project [28, 19]. With a multitude of analysis steps being combined into complex workflows, the documentation of the individual analyses and their dependencies can become challenging. To facilitate the automation of analysis tasks and their documentation, workflows can be described using workflow management systems such as Nextflow [29] or Snakemake [30]. Workflow managers that support the creation of reusable modules can help reduce complexity and promote the reuse of workflows or workflow modules [15, 16, 31].

Additionally, notebooks can apply the concept of literate programming and are a useful tool to add human-readable documentation next to code blocks [19]. Interoperability and reusability of workflows can be achieved using portable software containers such as Apptainer/Singularity [32] or Docker [33] that capture the runtime environment of a workflow or a workflow module [15, 34, 16, 25].

Computational workflows can be regarded as digital objects. The concept of FAIR Digital Objects (FDOs) was introduced to make digital objects fully FAIR [35]. FDOs comprise, among others, the digital object, a persistent identifier (PID), and metadata (title, authors, licenses, etc.) describing the object. Based on the FDO concept, the RO-Crate approach was specified to package digital research artefacts or Research Objects (RO) such as computational workflows [36]. The RO-Crate contains a PID that links to an RO, which is described by a structured JSON-LD RO-Crate metadata file. It can additionally contain data on which the workflow can be run. To make an RO-Crate findable, it needs to be registered at a registry such as WorkflowHub [37, 38]. In case the actual data cannot be publicly shared due to privacy reasons, synthetic data can complement analysis workflows to demonstrate the computational procedure [16, 39].

We here demonstrate the development of a FAIR Digital Object comprising a computational workflow that analyzes and integrates multi-omics and phenotype data and is associated with rich human and machine-readable metadata.

## Findings

### Workflow Implementation

To develop a reusable workflow, our input data and intermediate files were largely based on open and widely-used formats or community standards. For the metabolomics data and metadata, we adopted practices of the MetaboLights database [40] of the European Bioinformatics Institute (EBI) of the European Molecular Biology Laboratory (EMBL). Metabolite levels and annotations are reported in metabolite annotation/assignment files (MAF). The experimental metadata for omics measurements is reported using the Investigation/Study/Assay (ISA) metadata framework [41]. We employed Jupyter [42] and the Python ISA API [43] to create ISA-Tab and ISA-JSON files [44]. For machine-readable descriptions of the experiments, ontology terms were used. Ontologies are standardized taxonomies of entities of a specific subject (domain) including definitions of relationships between these entities. Ontology terms refer to these entities [45]. Based on recommended standards from FAIRgenomes [3] and Metabolights [40], we preferably employed the following ontologies: National Cancer Institute Thesaurus (NCIT) [46], Experimental Factor Ontology (EFO) [47], Ontology for Biomedical Investigations (OBI) [48], Metabolomics Standards Initiative Ontology (MSIO) [49], Chemical Methods Ontology (CHMO) [50] and Chemical Entities of Biological Interest (ChEBI) [51]. The DNA methylation levels and associated metadata, behavioral data, and additional information about phenotypes or technical and biological covariates are stored as comma-separated values (CSV) files. This allows our computational workflow to be easily reusable and adaptable for other data sets. The workflow documentation [52] describes all input files used in the workflow and provides human-readable descriptions of every step of the workflow processing and integrating individual input data types. Each of these analysis steps is implemented in Python or R, and added as a module to the workflow. We employ Jupyter and R notebooks for implementing downstream analyses and visualization of results. We chose Nextflow as our workflow management system, since it allows flexible development, can be run on different platforms, supports containers, is well documented, and is already widely adopted by the bioinformatics community [31]. Each module of the workflow is provided with their own Docker container to ensure portability and eliminate the need for local software installations.

Finally, the Nextflow workflow is packaged as an RO-Crate. Besides the workflow and the synthetic data set, it contains a structured metadata file with machine-readable descriptions of input files and analysis steps (ro-crate-metadata.json). We preferably used EDAM—Ontology of bioscientific data analysis and data management [53] as it is recommended for workflow RO-Crates [36]. For terms that were not available in EDAM, alternative ontologies such as NCIT [46], OBI [48], or the Semanticscience Integrated Ontology (SIO) [54] were used. The RO-Crate further contains an image with an overview of the analysis steps. For findability, the packaged workflow (see Figure 2) is registered on WorkflowHub [37] and provided with a Digital Object Identifier (DOI) [https://doi.org/10.48546/workflowhub.workflow.402.5].

### Case Study

Our workflow was developed to analyze and integrate DNA methylation and urine metabolomics profiles with behavioral data originating from the ACTION Biomarker Study (ACTION, Aggression in Children: Unraveling gene-environment interplay to inform Treatment and InterventiON strategies) [55, 56, 57] (see ‘Case Study’ in the ‘Methods’ section on page 5). Within ACTION, urine and buccal-cell samples were collected in a twin cohort from the Netherlands Twin Register (NTR), and in a cohort of children referred to an academic center for child and youth psychiatry in the Netherlands (LUMC-Curium). These children were also characterized for behavioral problems and here we look at externalizing problems. We purposely selected a case of complex human behavioral phenotype that is typically not caused by a single well-defined molecular defect, but originates from changes in multiple factors, and as such would benefit from a multi-omics analysis. Since we consider this data to be potentially personally identifiable information, we share a synthetic data set to demonstrate the workflow. The goal of the analysis is the identification of substructures in the multi-omics data and to determine if they correlate with behavioral data (see ‘Unsupervised Data Analysis’ on page 6). A team comprising members of the Netherlands X-omics Initiative [58] in collaboration with the Netherlands Twin Register (NTR) [59] developed the computational workflow. An overview of the main analysis steps is shown in Figure 1.

**Figure 1.**
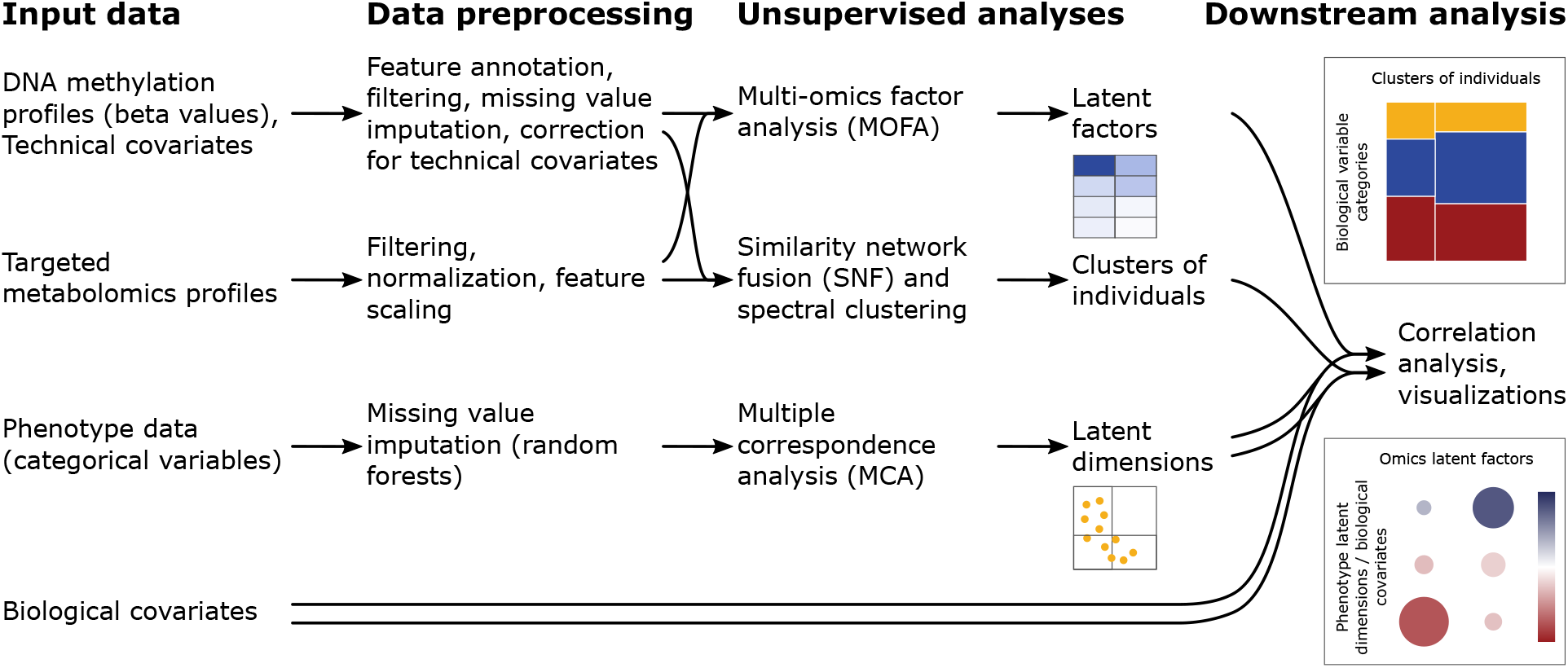
Overview of analysis steps.

**Figure 2.**
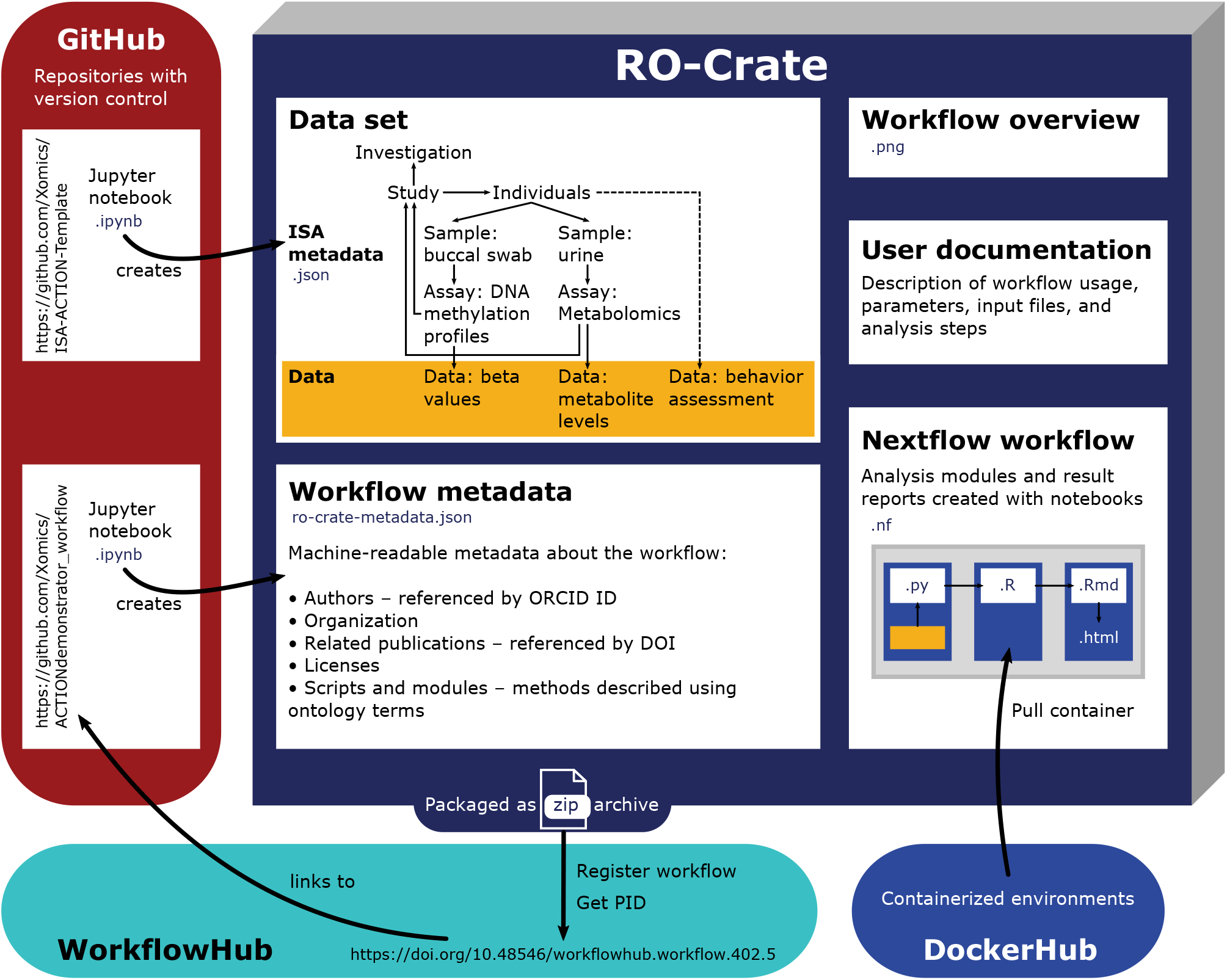
Schematic overview of packaged workflow.

To identify underlying patterns in childhood externalizing behavior, we applied Multiple Correspondence Analysis (MCA) [60, 61] to the parent-rated responses on the externalizing behavior items of the Child Behavior Checklist (CBCL) of the Achenbach System of Empirically Based Assessment (ASEBA) [62] in both cohorts. In NTR participants, the first three MCA dimensions jointly explain 30% of the variation in 26 externalizing behavior items of the ASEBA CBCL [see Additional File 1]. Additional dimensions each explain <5% of the variation. The presence rather than the absence of externalizing behaviors characterized all of the first three dimensions, which reflects the answer options to items (a problem behavior is not present, a little, or a lot). Variables that contributed most to the first dimension, which explained 16% of the variation, represent temperamental behavior (frequent temper tantrums, stubbornness, screaming, and arguing). Variables contributing to the second dimension, which explained 9% of the variation, represent hostile aggressive behaviors (frequent vandalism, bullying, and cruelty). In LUMC-Curium participants, the first two MCA dimensions suffice to explain 30% of the variation in 18 items of the ASEBA CBCL [see Additional File 2]. Similar to NTR, these first dimensions in LUMC-Curium are characterized by the presence of aggressive behaviors.

We applied Multi-Omics Factor Analysis (MOFA) [63] in both cohorts to obtain ten factors to describe the buccal DNA methylation (Illumina EPIC array) and urine metabolomics data. For this analysis, we selected the top 10% most variable probes from DNA methylation data. Cumulatively, the ten factors explained 22.5% and 74.9% of variation in the DNA methylation data and 0.001% and 1.89% in the urine metabolomics data in NTR [see Additional File 3] and LUMC-Curium [see Additional File 4], respectively. We observed no evidence that any of the factors captured sources of variation in both the DNA methylation and urine metabolomics data in NTR and LUMC-Curium. Particularly factors 1 and 2 in NTR and factor 1 in LUMC-Curium were specific to the DNA methylation data. To help elucidate the etiology of the ten MOFA factors, we selected for each factor the top 100 CpGs with the largest weights and performed enrichment analyses within the Epigenome-Wide Association Study (EWAS) atlas [64]. Multiple factors in both cohorts [see Additional File 5 for ACTION-NTR and Additional Files 6 for LUMC-Curium cohort] showed enrichment of CpGs associated with glucocorticoid exposure (i.e., administration of corticosteroid medication [65]), CpGs associated with ageing, and CpGs associated with immune-related traits, such as psoriasis. Apart from these robustly enriched traits, additional significant enrichments were found but were often based on ^≤^ 5 overlapping CpGs between the factor results and the original studies. A limitation of the enrichment analysis is that the majority of previous EWAS studies included in this analysis were conducted on blood samples from adult populations with the Illumina 450K BeadChip. In the factor weights for metabolites, we observe that for both NTR [Additional File 3] and LUMC-Curium [Additional File 4] many of the factors are characterized by only one or few metabolites. We note that in both cohorts the factors explained only a small amount of variation in the metabolomics data. To investigate whether the omics factors were associated with behavioral dimensions (MCA), we ran gee models adjusting for relatedness in NTR, and correlation analyses in Curium [see Additional File 3 for ACTION-NTR and Additional Files 4 for LUMC-Curium cohort]. None of the omics factors were significantly associated with the behavioral dimensions in NTR or LUMC-Curium, nor did we observe significant associations of sex-and-age-specific T-scores for aggressive behavior with the omics factors. In previous multi-omics analyses of high versus low levels of childhood aggression [66] and Attention-Deficit/Hyperactivity Disorder [67] we applied supervised analyses in these cohorts while applying unsupervised analyses here. In these previous supervised analyses, where we also included an additional omics layer—polygenic scores—we found that although multi-omics models had low predictive value, they revealed some connections of omics traits with externalizing problems, that suggested biological plausibility.

We also constructed integrated similarity networks with Similarity Network Fusion (SNF) [68] to identify subgroups of individuals based on omics data. In both NTR and LUMC-Curium, we defined integrated similarity networks based on two and four clusters. The two clusters in NTR were characterized by differences in age whereas the two clusters in LUMC-Curium were characterized by differences in the proportion of boys and girls. To investigate whether the omics clusters were associated with externalizing behavior, we compared the behavioral dimension scores from MCA between children in the different clusters. In both NTR and LUMC-Curium, we observed no significant differences in the behavioral dimensions across the two omics clusters after correction for multiple testing [see Additional File 7 for NTR and Additional File 8 for LUMC-Curium cohort]. Similarly, no differences in behavioral dimensions were observed between the four omics clusters in NTR, however, in LUMC-Curium, behavioral dimension 6 differed significantly between the four omics clusters. In LUMC-Curium, dimension 6 explained 3.9% of the variance in childhood externalizing behavior and the strongest contributors to this dimension comprised higher frequencies of parent-rated tendencies to be suspicious and loud. Such forms of direct aggressive behavior, particularly physical aggressive behavior, are common in early childhood in both boys and girls, and while overall levels of aggression decline with age and are roughly similar for boys and girls [69], boys are more likely to engage in direct and physical forms of aggression by age 11 [70]. Thus, this finding aligns with the observation that the two omics clusters differ in the proportion of males and females and in the age composition.

## Discussion

In this collaborative research project, partners from the Netherlands X-omics Initiative co-developed a workflow to analyze a complex multimodal data set. Developing workflows with partners across multiple institutions can pose a challenge and we experienced that a secure shared computing environment was key to the success of this project. Additionally, practices aiming to increase FAIRness of the shared workflow such as version control with git and a modular workflow structure allowed for transparent and target-oriented workflow development. Therefore, while the use of technologies like git or workflow management systems might require initial training of researchers, we believe this to be worthwhile.

Several FAIR practices for workflows include existing best practices of software development, for example, version control and good documentation. Adoption of these practices, along with the use of workflow managers and software containers, aims to contribute to better interoperability, reusability and reproducibility of analysis workflows and research results. While we experienced the adoption of these technologies to be straightforward, fully FAIR data or software requires also machine-understandable semantic metadata. Specifications like the ISA metadata framework and RO-Crate allow ontology-based annotations of omics experiments and analysis workflows, respectively. Our choice of ontologies was mainly guided by the documented submission requirements or recommendations provided by services such as the MetaboLights archive or WorkflowHub. However, when recommended ontologies do not comprise suitable terms, choosing appropriate ones from ontologies can be challenging. For example, no exact match to the generic term *sample collection* that is part of the ISA schema can be found in any ontology available in EBI’s Ontology Lookup Serivce (OLS) [71]. To describe workflow steps in RO-Crate with *unsupervised learning* we had to employ the eNanoMapper Ontology [72] as no matching term was available in the recommended EDAM ontology. Consequently, we recognize the importance of teams dedicated to ontology curation, active user communities, and training of researchers in using semantic technologies. This is especially important for multi-omics research that spans multiple research domains.

For reproducibility of research results, it is essential that data are shared along with the workflow. However, privacy regulations prohibit sharing of potentially personally identifiable data such as omics measurements or clinical information. To demonstrate the functionality of the workflow, we shared a synthetic data set that emulates the structure of the case study data set. Current developments in the areas of federated data storage and analysis such as Federated European Genome-phenome Archive (EGA) [73] and the Personal Health Train [74] have the potential to allow fully FAIR and reproducible data analysis workflows while maintaining privacy regulation compliance.

We hope our experiences help other researchers who develop multi-omics data analysis workflows choosing and implementing practices that makes their research more FAIR.

## Data and Methods

### Case Study

Our case study comprises data from two cohorts that took part in the ACTION Biomarker Study (ACTION, Aggression in Children: Unraveling gene-environment interplay to inform Treatment and InterventiON strategies) [55, 56, 57]. The ACTION Biomarker Study collected buccal DNA samples for large-scale genome-wide and epigenome-wide association studies [75, 76] and first-morning urine samples to investigate the association of urine biomarkers and metabolites with childhood aggression [57]. These urine and buccal-cell samples were collected in a twin cohort from the Netherlands Twin Register (NTR) [77], where twin pairs were selected on their longitudinal concordance or discordance for childhood aggression, and in a cohort of children referred to an academic center for child and youth psychiatry in the Netherlands (LUMC-Curium). The DNA methylation, genotype, metabolomics, and behavioural data from these cohorts were previously used for multi-omics analyses of aggressive behaviour [66] and Attention Deficit Hyperactivity Disorder (ADHD) [67]. Detailed information on the study populations and study protocol is available at protocols.io [78].

#### Data

Genome-wide DNA methylation data in buccal DNA samples were measured on the Infinium MethylationEPIC BeadChip kit (Illumina, San Diego, CA, USA [79]) by the Human Genotyping Facility (HuGeF) of ErasmusMC (the Netherlands; [80]). The ZymoResearch EZ DNA Methylation kit (Zymo Research Corp, Irvine, CA, USA) was used for bisulfite treatment of 500 ng of genomic DNA obtained from buccal swabs. The Infinium HD Methylation Assay was performed according to the manufacturer’s specification. Good Biomarker Sciences Leiden (Leiden, the Netherlands) measured the specific gravity (by refractomertry), levels of creatinine (by colorimertry), blood traces, markers of leukocytes, proteins, glucose and nitrites (the latter five by dipstick) of each urine sample. The Metabolomics Facility of the University of Leiden (Leiden, the Netherlands) quantified urine metabolites using three platforms: a liquid chromatography mass spectrometry (LC-MS) platform targeting amines (66 biomarkers), an LC-MS platform targeting steroid hormones (13 biomarkers), and a gas chromatography (GC)-MS platform targeting organic acids (21 biomarkers). Behavioral data comprises the 115 items of the Dutch version of the Achenbach System of Empirically Based Assessment (ASEBA) Child Behavior Checklist (CBCL) for school-aged children (6–18 years) [62]. For participants of the NTR cohort we used the mother-rated CBCL as completed at the time of biological sample collection, and for participants of the LUMC-Curium cohort we used the parent-rated (90% mother ratings) CBCL as completed in a 6-month window surrounding the biological sample collection. Again, details on the data generation are available in Ref. [78].

#### Synthetic Data

The purpose of the synthetic data set is to demonstrate how the workflow can be run. It resembles the structure of the files of the cohort data. The values were randomly sampled from the observed values in the NTR cohort without preserving any correlations.

### Data Processing

To ensure the urine sample metabolic integrity and to minimize bias contributed by health conditions, we excluded samples from the metabolomics data from *(1)* subjects who have started menstruating, *(2)* subjects where the time between urine sample collection and storing in the freezer was > 2 hours, *(3)* subjects where severe violations to the sampling protocol occurred (e.g. not putting a lid on the container), *(4)* subjects where the leukocyte count was above trace, *(5)* subjects where the nitrites level was “positive high”, *(6)* subjects where the protein level was > 0.3, *(7)* subjects with glucose levels above trace, *(8)* subjects with blood levels above trace, *(9)* subjects having the flu, *(10)* subjects reporting inflammation, *(11)* subjects reporting vomiting, *(12)* subjects reporting abdominal pain and *(12)* subjects reporting general health problems. Note that the above criteria 4–8 are based on the dipstick marker estimation performed separately from the metabolomics measurements on the same samples [78], while the other criteria are based on questionnaire data at the time of sampling.

The metabolomics features were filtered based on missing values. Missing values were reported for cases where the metabolite concentration is below the limit of quantification. Samples and metabolites with 15% or more of missing values were discarded. Sample-wise normalization to correct for urine concentration was conducted by adjusting metabolite intensities to the sample creatinine levels [78]. This was followed metabolite-wise by Pareto scaling [81] to statistically account for large differences in reported values.

Quality Control (QC) and normalization of the DNA methylation array data have been previously described [75] and were carried out with a pipeline developed by the Biobank-based Integrative Omics Study (BIOS) consortium [82]. From the 787,711 autosomal methylation probes that survived QC, the top 10% most variable probes were included in the analyses. Cellular proportions of buccal samples were predicted with Hierarchical Epigenetic Dissection of IntraSample-Heterogeneity (HepiDISH) with the RPC method (reduced partial correlation), as described Zheng et al. [83] and implemented in the R/Bioconductor package EpiDISH. Median imputation was carried out on the epigenetics data. Residual methylation levels were obtained by regressing the effects of percentages of epithelial and natural killer cells, EPIC array row, and bisulfite sample plate, from the methylation beta-values.

Missing values in the externalizing behavior items were imputed with the non-parametric random forests method from the R library missForest (1.4) [84].

### Unsupervised Data Analysis

Each cohort was analyzed separately. We applied Multi-Omics Factor Analysis (MOFA) using the R/Bioconductor library MOFA2 (1.3.4) [63, 85] to obtain factors for the buccal DNA methylation and urine metabolomics data and applied Multiple Correspondence Analysis (MCA) [61] using the R library FactoMineR (2.4) [60] to obtain factors for the behavioral data.

To identify subgroups of individuals based on their buccal DNA methylation and urine metabolomics data we constructed integrated similarity networks with Similarity Network Fusion (SNF) [68]. The optimal numbers of clusters were determined using a built-in function of the Python library SNFpy [86] that uses the eigengap method [87] to find the optimal number of clusters. SNF first constructs sample similarity networks for each available data type and then fuses these into a single network comprising both the shared and unique information from each data type. The final fused network thus captures how each data type contributes to the similarity amongst the samples. We tested whether the behavioral dimension scores from MCA differ between children in the different SNF clusters, using Mann-Whitney U tests (two clusters) or Kruskal-Wallis tests (four clusters) in the Curium cohort, and with generalized estimation equation (GEE) models (with cluster as predictor and behavioral dimension score as outcome) in NTR.

We determined correlations amongst the obtained factors capturing the omics and behavioral data, respectively, using Spearman’s rank correlation, and additionally in the NTR cohort using GEE models. All GEE models were fitted with the R package GEE, with the following specifications: Gaussian link function (for continuous data), 100 iterations, and the ‘exchangeable’ option to account for the correlations in twin pairs. Statistical test were adjusted for multiple testing using false discovery rate (FDR) [88].

## Workflow Implementation

### Availability of source code and requirements

- Project name: X-omics ACTION demonstrator
- Project home page: [89]
- Operating system(s): Platform independent
- Programming language: Python, R
- Other requirements: Nextflow (22.04.0), Docker (19.03.1), Singularity (3.8.0)
- License: MIT

### Additional Files

- File name: Additional File 1
- File format: .html
- Title: Multiple Correspondence Analysis of CBCL behavioral data
- Description: Overview of externalizing behavior items of the Child Behavior Checklist (CBCL) of the Achenbach System of Empirically Based Assessment (ASEBA) in the ACTION-NTR cohort before and after imputation of missing values using random forests and results and visualizations of Multiple Correspondence Analysis (MCA).
- File name: Additional File 2
- File format: .html
- Title: Multiple Correspondence Analysis of CBCL behavioral data
- Description: Overview of externalizing behavior items of the Child Behavior Checklist (CBCL) of the Achenbach System of Empirically Based Assessment (ASEBA) in the LUMC-Curium cohort before and after imputation of missing values using random forests and results and visualizations of Multiple Correspondence Analysis (MCA).
- File name: Additional File 3
- File format: .html
- Title: MOFA downstream analysis report
- Description: Visualizations of Multi-omics Factor Analysis (MOFA) of buccal DNA methylation (Illumina EPIC array) and urine metabolomics data of the ACTION-NTR cohort. Also, associations between the MOFA factors and phenotypic data are tested with GEE models.
- File name: Additional File 4
- File format: .html
- Title: MOFA downstream analysis report
- Description: Visualizations of Multi-omics Factor Analysis (MOFA) of buccal DNA methylation (Illumina EPIC array) and urine metabolomics data of the LUMC-Curium cohort.
- File name: Additional File 5
- File format: .xlsx
- Title: EWAS atlas enrichment analysis
- Description: Enriched traits for CpGs with top 100 largest weights of ACTION-NTR MOFA factors 1-10.
- File name: Additional File 6
- File format: .xlsx
- Title: EWAS atlas enrichment analysis
- Description: Enriched traits for CpGs with top 100 largest weights of LUMC-Curium MOFA factors 1-10.
- File name: Additional File 7
- File format: .html
- Title: Similarity Network Fusion downstream analysis
- Description: Visualizations of Similarity Network Fusions (SNF) and subsequent spectral clustering of buccal DNA methylation (Illumina EPIC array) and urine metabolomics data of the ACTION-NTR cohort.
- File name: Additional File 8
- File format: .html
- Title: Similarity Network Fusion downstream analysis
- Description: Visualizations of Similarity Network Fusions (SNF) and subsequent spectral clustering of buccal DNA methylation (Illumina EPIC array) and urine metabolomics data of the LUMC-Curium cohort.
- File name: Additional File 9
- File format: .html
- Title: Similarity Network Fusion downstream analysis with GEE models
- Description: Associations between the SNF clusters and phenotypic data are tested with GEE models in the ACTION-NTR cohort.

## Data Availability

The data of the Netherlands Twin Register (NTR) ACTION Biomarker Study may be accessed, upon approval of the data access committee, through the NTR [90].

A synthetic data set representing the structure of the ACTION Biomarker Study data set is available as part of the workflow RO-Crate available at WorkflowHub [https://doi.org/10.48546/workflowhub.workflow.402.5].

## Declarations

## List of abbreviations

ACTION: Aggression in
Children: Unraveling gene-environment interplay to inform Treatment and InterventiON strategies;
ASEBA: Achenbach System of Empirically Based Assessment;
EBI: European Bioinformatics Institute;
EGA: European Genome-phenome Archive;
EMBL: European Molecular Biology Laboratory;
EWAS: epigenome-wide association study;
FAIR: Findable, Accessible, Interoperable, Reusable;
GC-MS: gas chromatography–mass spectrometry;
ISA: Investigation, Study, Assay;
LC-MS: liquid chromatography–mass spectrometry;
MOFA: Multi-Omics Factor Analysis;
NTR: Netherlands Twin Register;
RO-Crate: Research Object Crate;
SNF: Similarity Network Fusion

## Ethical Approval

The ACTION Biomarker study was conducted according to the guidelines of the Declaration of Helsinki and approved by the Central Ethics Committee on Research Involving Human Subjects of the VU University Medical Center, Amsterdam (NTR 25th of May 2007 and ACTION 2013/41 and 2014.252), an Institutional Review Board certified by the U.S. Office of Human Research Protections (IRB number IRB00002991 under Federal-wide Assurance- FWA00017598; IRB/institute codes), and the Medical Ethical Committee of Leiden University Medical Center (B17.031, B17.032 and B17.040).

Parents provided written informed consent for the participation of their children.

## Consent for publication

Not applicable

## Authors’ Contributions

A.J.G, P.A.C.H., D.I.B. and J.D. formulated overarching research aims (Conceptualization). R.P. and N.K. maintained and annotated the metabolomics data of the ACTION-NTR cohort (Data curation). C.V., A.N., P.K., and JD applied statistical methods to analyze the study data (Formal Analysis). A.J.G., P.A.C.H., R.R.J.M.V and D.I.B. acquired financial support for the projects leading to this publication (Funding acquisition). A.N., C.V., F.A.H., N.K., A.S.D.K., P.K., R.P. and J.D. conducted the research process (Investigation). A.N., C.V., F.A.H., N.K., A.S.D.K., P.K., R.P. and J.D. designed the methodology (Methodology). A.N. coordinated the research activity planning and execution (Project administration). F.A.H., R.P., J.D., R.R.J.M.V. and D.B. provided study samples and acquired data (Resources). C.V., P.K., R.P., J.D. and A.N. implemented the source code (Software). D.I.B., A.J.G. and P.A.C.H. provided mentorship to the research team (Supervision). A.N. wrote the initial draft of the manuscript (Writing – original draft). All authors critically reviewed and contributed to writing the manuscript (Writing – review & editing).

## Acknowledgements

We acknowledge the ACTION Consortium and thank the participants of the ACTION Biomarker Study. We thank Michael van Vliet for discussions on the FAIRification strategy and Philippe Rocca-Serra and his team for their support on ISA.

