## Additional File 9 for "A Multi-omics Data Analysis Workflow Packaged as a FAIR Digital Object": Additional File 9.html

SNF downstream analysis with GEE


### SNF downstream analysis with GEE

###### Casper de Visser^1^

##### Load SNF dataframe

```
print(params$input_file_snf_pheno)
```

```
## [1] "snf_analysis_out.csv"
```

```
df_snf_phenotypes <- read.csv(params$input_file_snf_pheno, row.names=1)
df_snf_phenotypes <- df_snf_phenotypes[order(df_snf_phenotypes$Familynumber),]
```

##### GEE model to correct for twin effect

```
library(gee)


#' Function for GEE model
#'
#' @param data Dataframe with pheno metadata
#' @param pheno_feature Feature to select as outcome
#' @param family Gaussian/Binomial
#' @return GEE model
geemodel <- function (data, pheno_feature, snf_label, family) {

f <- as.formula(
    paste(pheno_feature,
        paste(snf_label),
        sep = " ~ "))

invisible(gee(f, data=data, id=Familynumber, family=family, corstr="exchangeable", maxiter=100, na.action=na.omit, silent=T))
}


#' Function to run the GEE model with mofa factors as predictor for phenotypic feature
#'
#' @param data Dataframe with pheno metadata
#' @param pheno_feature Feature to select as outcome
#' @param family Gaussian/Binomial
#' @return end table with estimates and p-values
run_geemodel <- function(data, pheno_feature, snf_label, family=gaussian) {


# Make dataframes for storing statistics
estimates <- matrix(NA, 10, 2)
robustSE <- matrix(NA, 10, 2)
robustZ <- matrix(NA, 10, 2)
pval <- matrix(NA, 10, 2)


coeff <- summary(invisible(geemodel(data, pheno_feature, snf_label, family)))$coefficients
coeff <- data.frame(coeff)
coeff$pval <- 2*pnorm(-abs(coeff[,5]))
coeff$padjusted <- p.adjust(coeff$pval)

return(coeff)

}


### Set categorial variables to factors
```

```
df_snf_phenotypes$SNF_label_4 <- as.factor(df_snf_phenotypes$SNF_label_4) 
df_snf_phenotypes$SNF_label_2 <- as.factor(df_snf_phenotypes$SNF_label_2)
```

##### Rewrite male/female as 0/1

```
df_snf_phenotypes$Sex[df_snf_phenotypes$Sex == 'Male'] <- 0
df_snf_phenotypes$Sex[df_snf_phenotypes$Sex == 'Female'] <- 1
df_snf_phenotypes$Sex <- as.numeric(df_snf_phenotypes$Sex)
```

### GEE models with 4 SNF clusters as predictor

##### Aggression score and Age as outcome

```
geemodel_agg <- run_geemodel(df_snf_phenotypes, 'Aggression_Tscore', 'SNF_label_4')
geemodel_age <- run_geemodel(df_snf_phenotypes, 'Age', 'SNF_label_4')
```

##### Aggression score and Age as outcome

GEE model with Aggression T-score as outcome

|  | Estimate | Naive.S.E. | Naive.z | Robust.S.E. | Robust.z | pval | padjusted |
| --- | --- | --- | --- | --- | --- | --- | --- |
| (Intercept) | 52.0558379 | 0.5558342 | 93.6535404 | 0.5825707 | 89.3554042 | 0.0000000 | 0.0000000 |
| SNF\_label\_4SNF\_1 | -0.9686044 | 1.0185372 | -0.9509760 | 0.7995511 | -1.2114352 | 0.2257286 | 0.6771859 |
| SNF\_label\_4SNF\_2 | 0.0331735 | 0.7767149 | 0.0427100 | 0.7606449 | 0.0436123 | 0.9652135 | 1.0000000 |
| SNF\_label\_4SNF\_3 | -0.2162909 | 0.8609907 | -0.2512116 | 0.8899187 | -0.2430457 | 0.8079700 | 1.0000000 |

GEE model with Age as outcome

|  | Estimate | Naive.S.E. | Naive.z | Robust.S.E. | Robust.z | pval | padjusted |
| --- | --- | --- | --- | --- | --- | --- | --- |
| (Intercept) | 9.0859181 | 0.0804051 | 113.0017 | 0.0814033 | 111.6160344 | 0.0000000 | 0.0000000 |
| SNF\_label\_4SNF\_1 | -0.0065958 | NaN | NaN | 0.0162894 | -0.4049125 | 0.6855418 | 1.0000000 |
| SNF\_label\_4SNF\_2 | 0.0009252 | NaN | NaN | 0.0125712 | 0.0736008 | 0.9413281 | 1.0000000 |
| SNF\_label\_4SNF\_3 | 0.0603068 | NaN | NaN | 0.0562410 | 1.0722925 | 0.2835887 | 0.8507661 |

##### Male/Female as outcome

```
geemodel_sex <- run_geemodel(df_snf_phenotypes, 'Sex', 'SNF_label_4', 'binomial')
```

##### Male/Female as outcome

GEE model with Sex as outcome

|  | Estimate | Naive.S.E. | Naive.z | Robust.S.E. | Robust.z | pval | padjusted |
| --- | --- | --- | --- | --- | --- | --- | --- |
| (Intercept) | -0.1590829 | 0.0883589 | -1.8004170 | 0.0864617 | -1.8399237 | 0.0657794 | 0.2631178 |
| SNF\_label\_4SNF\_1 | -0.1083750 | 0.0933238 | -1.1612796 | 0.0737571 | -1.4693506 | 0.1417377 | 0.4252131 |
| SNF\_label\_4SNF\_2 | -0.0081028 | 0.0682986 | -0.1186385 | 0.0367946 | -0.2202186 | 0.8257009 | 0.8257009 |
| SNF\_label\_4SNF\_3 | -0.0499806 | 0.0760359 | -0.6573292 | 0.0549224 | -0.9100227 | 0.3628105 | 0.7256211 |

##### MCA dimension coordinates as outcome

```
geemodel_mca1 <- run_geemodel(df_snf_phenotypes, 'Dim_1', 'SNF_label_4')
geemodel_mca2 <-run_geemodel(df_snf_phenotypes, 'Dim_2', 'SNF_label_4')
geemodel_mca3 <-run_geemodel(df_snf_phenotypes, 'Dim_3', 'SNF_label_4')
geemodel_mca4 <-run_geemodel(df_snf_phenotypes, 'Dim_4', 'SNF_label_4')
geemodel_mca5 <-run_geemodel(df_snf_phenotypes, 'Dim_5', 'SNF_label_4')
geemodel_mca6 <-run_geemodel(df_snf_phenotypes, 'Dim_6', 'SNF_label_4')
geemodel_mca7 <-run_geemodel(df_snf_phenotypes, 'Dim_7', 'SNF_label_4')
geemodel_mca8 <-run_geemodel(df_snf_phenotypes, 'Dim_8', 'SNF_label_4')
geemodel_mca9 <-run_geemodel(df_snf_phenotypes, 'Dim_9', 'SNF_label_4')
geemodel_mca10 <-run_geemodel(df_snf_phenotypes, 'Dim_10', 'SNF_label_4')
```

##### Print GEE models

GEE model with MCA dimension 1 as outcome

|  | Estimate | Naive.S.E. | Naive.z | Robust.S.E. | Robust.z | pval | padjusted |
| --- | --- | --- | --- | --- | --- | --- | --- |
| (Intercept) | 0.0130994 | 0.0262644 | 0.4987501 | 0.0272114 | 0.4813924 | 0.6302377 | 1.0000000 |
| SNF\_label\_4SNF\_1 | -0.0429187 | 0.0499502 | -0.8592294 | 0.0350874 | -1.2231934 | 0.2212567 | 0.8850267 |
| SNF\_label\_4SNF\_2 | -0.0126438 | 0.0382894 | -0.3302164 | 0.0359064 | -0.3521314 | 0.7247397 | 1.0000000 |
| SNF\_label\_4SNF\_3 | -0.0075180 | 0.0424343 | -0.1771683 | 0.0436915 | -0.1720703 | 0.8633822 | 1.0000000 |

GEE model with MCA dimension 2 as outcome

|  | Estimate | Naive.S.E. | Naive.z | Robust.S.E. | Robust.z | pval | padjusted |
| --- | --- | --- | --- | --- | --- | --- | --- |
| (Intercept) | 0.0005117 | 0.0192017 | 0.0266491 | 0.0211547 | 0.0241888 | 0.9807020 | 1 |
| SNF\_label\_4SNF\_1 | -0.0252961 | 0.0455229 | -0.5556773 | 0.0327886 | -0.7714904 | 0.4404163 | 1 |
| SNF\_label\_4SNF\_2 | -0.0108700 | 0.0366296 | -0.2967540 | 0.0280996 | -0.3868373 | 0.6988767 | 1 |
| SNF\_label\_4SNF\_3 | -0.0223005 | 0.0407679 | -0.5470116 | 0.0314661 | -0.7087167 | 0.4785003 | 1 |

GEE model with MCA dimension 3 as outcome

|  | Estimate | Naive.S.E. | Naive.z | Robust.S.E. | Robust.z | pval | padjusted |
| --- | --- | --- | --- | --- | --- | --- | --- |
| (Intercept) | 0.0121847 | 0.0159622 | 0.7633454 | 0.0166482 | 0.7318925 | 0.4642342 | 1 |
| SNF\_label\_4SNF\_1 | 0.0022983 | 0.0351718 | 0.0653450 | 0.0256874 | 0.0894721 | 0.9287067 | 1 |
| SNF\_label\_4SNF\_2 | -0.0176608 | 0.0277413 | -0.6366252 | 0.0262490 | -0.6728182 | 0.5010630 | 1 |
| SNF\_label\_4SNF\_3 | -0.0146461 | 0.0307540 | -0.4762323 | 0.0234485 | -0.6246054 | 0.5322301 | 1 |

GEE model with MCA dimension 4 as outcome

|  | Estimate | Naive.S.E. | Naive.z | Robust.S.E. | Robust.z | pval | padjusted |
| --- | --- | --- | --- | --- | --- | --- | --- |
| (Intercept) | 0.0074012 | 0.0153573 | 0.4819315 | 0.0187987 | 0.3937055 | 0.6937985 | 1 |
| SNF\_label\_4SNF\_1 | -0.0022851 | 0.0364107 | -0.0627584 | 0.0259640 | -0.0880095 | 0.9298692 | 1 |
| SNF\_label\_4SNF\_2 | -0.0101754 | 0.0292979 | -0.3473086 | 0.0186368 | -0.5459867 | 0.5850751 | 1 |
| SNF\_label\_4SNF\_3 | -0.0112515 | 0.0326081 | -0.3450531 | 0.0224535 | -0.5011040 | 0.6162980 | 1 |

GEE model with MCA dimension 5 as outcome

|  | Estimate | Naive.S.E. | Naive.z | Robust.S.E. | Robust.z | pval | padjusted |
| --- | --- | --- | --- | --- | --- | --- | --- |
| (Intercept) | -0.0021967 | 0.0116341 | -0.1888168 | 0.0112045 | -0.1960565 | 0.8445659 | 1 |
| SNF\_label\_4SNF\_1 | 0.0088195 | 0.0238910 | 0.3691545 | 0.0163463 | 0.5395394 | 0.5895147 | 1 |
| SNF\_label\_4SNF\_2 | -0.0148768 | 0.0185541 | -0.8018071 | 0.0180396 | -0.8246737 | 0.4095569 | 1 |
| SNF\_label\_4SNF\_3 | -0.0301931 | 0.0205573 | -1.4687288 | 0.0271691 | -1.1113040 | 0.2664375 | 1 |

GEE model with MCA dimension 6 as outcome

|  | Estimate | Naive.S.E. | Naive.z | Robust.S.E. | Robust.z | pval | padjusted |
| --- | --- | --- | --- | --- | --- | --- | --- |
| (Intercept) | 0.0123764 | 0.0109322 | 1.1321073 | 0.0114453 | 1.0813473 | 0.2795426 | 1 |
| SNF\_label\_4SNF\_1 | -0.0020667 | 0.0248311 | -0.0832316 | 0.0212359 | -0.0973224 | 0.9224704 | 1 |
| SNF\_label\_4SNF\_2 | -0.0161579 | 0.0197402 | -0.8185294 | 0.0175580 | -0.9202606 | 0.3574366 | 1 |
| SNF\_label\_4SNF\_3 | -0.0253440 | 0.0219040 | -1.1570506 | 0.0230580 | -1.0991450 | 0.2717048 | 1 |

GEE model with MCA dimension 7 as outcome

|  | Estimate | Naive.S.E. | Naive.z | Robust.S.E. | Robust.z | pval | padjusted |
| --- | --- | --- | --- | --- | --- | --- | --- |
| (Intercept) | 0.0093735 | 0.0100402 | 0.9335948 | 0.0108303 | 0.8654879 | 0.3867711 | 1 |
| SNF\_label\_4SNF\_1 | -0.0155501 | 0.0231030 | -0.6730755 | 0.0167006 | -0.9311106 | 0.3517963 | 1 |
| SNF\_label\_4SNF\_2 | -0.0186805 | 0.0184322 | -1.0134739 | 0.0171847 | -1.0870429 | 0.2770178 | 1 |
| SNF\_label\_4SNF\_3 | -0.0009790 | 0.0204656 | -0.0478339 | 0.0159114 | -0.0615250 | 0.9509411 | 1 |

GEE model with MCA dimension 8 as outcome

|  | Estimate | Naive.S.E. | Naive.z | Robust.S.E. | Robust.z | pval | padjusted |
| --- | --- | --- | --- | --- | --- | --- | --- |
| (Intercept) | -0.0074307 | 0.0106107 | -0.7003083 | 0.0113420 | -0.6551535 | 0.5123689 | 1 |
| SNF\_label\_4SNF\_1 | 0.0098908 | 0.0221863 | 0.4458059 | 0.0188834 | 0.5237825 | 0.6004298 | 1 |
| SNF\_label\_4SNF\_2 | -0.0108337 | 0.0172926 | -0.6264964 | 0.0149514 | -0.7245967 | 0.4686995 | 1 |
| SNF\_label\_4SNF\_3 | 0.0281611 | 0.0191601 | 1.4697790 | 0.0270401 | 1.0414575 | 0.2976633 | 1 |

GEE model with MCA dimension 9 as outcome

|  | Estimate | Naive.S.E. | Naive.z | Robust.S.E. | Robust.z | pval | padjusted |
| --- | --- | --- | --- | --- | --- | --- | --- |
| (Intercept) | 0.0127300 | 0.0104009 | 1.2239277 | 0.0108449 | 1.1738190 | 0.2404675 | 0.7214026 |
| SNF\_label\_4SNF\_1 | 0.0080819 | 0.0224982 | 0.3592228 | 0.0151345 | 0.5340030 | 0.5933395 | 1.0000000 |
| SNF\_label\_4SNF\_2 | -0.0338236 | 0.0176666 | -1.9145536 | 0.0149971 | -2.2553417 | 0.0241119 | 0.0964476 |
| SNF\_label\_4SNF\_3 | -0.0011724 | 0.0195793 | -0.0598816 | 0.0229500 | -0.0510869 | 0.9592563 | 1.0000000 |

GEE model with MCA dimension 10 as outcome

|  | Estimate | Naive.S.E. | Naive.z | Robust.S.E. | Robust.z | pval | padjusted |
| --- | --- | --- | --- | --- | --- | --- | --- |
| (Intercept) | -0.0087390 | 0.0101703 | -0.8592658 | 0.0106898 | -0.8175064 | 0.4136391 | 1.0000000 |
| SNF\_label\_4SNF\_1 | 0.0343186 | 0.0231616 | 1.4817009 | 0.0185250 | 1.8525600 | 0.0639454 | 0.2557818 |
| SNF\_label\_4SNF\_2 | 0.0056874 | 0.0184262 | 0.3086597 | 0.0143452 | 0.3964692 | 0.6917589 | 1.0000000 |
| SNF\_label\_4SNF\_3 | 0.0113915 | 0.0204484 | 0.5570845 | 0.0182222 | 0.6251415 | 0.5318782 | 1.0000000 |

### GEE models with 2 SNF clusters as predictor

##### Aggression score as outcome

```
geemodel_agg <- run_geemodel(df_snf_phenotypes, 'Aggression_Tscore', 'SNF_label_2')
geemodel_age <- run_geemodel(df_snf_phenotypes, 'Age', 'SNF_label_2')
```

##### Aggression score as outcome

GEE model with Aggression T-score as outcome

|  | Estimate | Naive.S.E. | Naive.z | Robust.S.E. | Robust.z | pval | padjusted |
| --- | --- | --- | --- | --- | --- | --- | --- |
| (Intercept) | 51.7627798 | 0.7214374 | 71.7495090 | 0.7525637 | 68.7819283 | 0.0000000 | 0.0000000 |
| SNF\_label\_2SNF\_1 | 0.2055225 | 0.6983032 | 0.2943169 | 0.7657876 | 0.2683805 | 0.7884064 | 0.7884064 |

GEE model with Aggression T-score as outcome

|  | Estimate | Naive.S.E. | Naive.z | Robust.S.E. | Robust.z | pval | padjusted |
| --- | --- | --- | --- | --- | --- | --- | --- |
| (Intercept) | 9.1383932 | 0.0774236 | 118.0311 | 0.0846003 | 108.018511 | 0.0000000 | 0.0000000 |
| SNF\_label\_2SNF\_1 | -0.0582866 | NaN | NaN | 0.0405686 | -1.436741 | 0.1507916 | 0.1507916 |

##### Male/Female as outcome

```
geemodel_sex <- run_geemodel(df_snf_phenotypes, 'Sex', 'SNF_label_2', 'binomial')
```

##### Male/Female as outcome

GEE model with Sex as outcome

|  | Estimate | Naive.S.E. | Naive.z | Robust.S.E. | Robust.z | pval | padjusted |
| --- | --- | --- | --- | --- | --- | --- | --- |
| (Intercept) | -0.1806008 | 0.0979279 | -1.8442224 | 0.0905499 | -1.9944893 | 0.0460986 | 0.0921972 |
| SNF\_label\_2SNF\_1 | 0.0000923 | 0.0636263 | 0.0014505 | 0.0383636 | 0.0024057 | 0.9980806 | 0.9980806 |

```
geemodel_mca1 <- run_geemodel(df_snf_phenotypes, 'Dim_1', 'SNF_label_2')
geemodel_mca2 <-run_geemodel(df_snf_phenotypes, 'Dim_2', 'SNF_label_2')
geemodel_mca3 <-run_geemodel(df_snf_phenotypes, 'Dim_3', 'SNF_label_2')
geemodel_mca4 <-run_geemodel(df_snf_phenotypes, 'Dim_4', 'SNF_label_2')
geemodel_mca5 <-run_geemodel(df_snf_phenotypes, 'Dim_5', 'SNF_label_2')
geemodel_mca6 <-run_geemodel(df_snf_phenotypes, 'Dim_6', 'SNF_label_2')
geemodel_mca7 <-run_geemodel(df_snf_phenotypes, 'Dim_7', 'SNF_label_2')
geemodel_mca8 <-run_geemodel(df_snf_phenotypes, 'Dim_8', 'SNF_label_2')
geemodel_mca9 <-run_geemodel(df_snf_phenotypes, 'Dim_9', 'SNF_label_2')
geemodel_mca10 <-run_geemodel(df_snf_phenotypes, 'Dim_10', 'SNF_label_2')
```

##### Print GEE models

GEE model with MCA dimension 1 as outcome

|  | Estimate | Naive.S.E. | Naive.z | Robust.S.E. | Robust.z | pval | padjusted |
| --- | --- | --- | --- | --- | --- | --- | --- |
| (Intercept) | 0.0148916 | 0.0346458 | 0.4298234 | 0.0420593 | 0.3540621 | 0.7232923 | 1 |
| SNF\_label\_2SNF\_1 | -0.0133989 | 0.0343088 | -0.3905396 | 0.0441071 | -0.3037815 | 0.7612944 | 1 |

GEE model with MCA dimension 2 as outcome

|  | Estimate | Naive.S.E. | Naive.z | Robust.S.E. | Robust.z | pval | padjusted |
| --- | --- | --- | --- | --- | --- | --- | --- |
| (Intercept) | 0.032820 | 0.0285811 | 1.148314 | 0.0479460 | 0.6845208 | 0.4936464 | 0.5862301 |
| SNF\_label\_2SNF\_1 | -0.053235 | 0.0323508 | -1.645556 | 0.0506367 | -1.0513126 | 0.2931150 | 0.5862301 |

GEE model with MCA dimension 3 as outcome

|  | Estimate | Naive.S.E. | Naive.z | Robust.S.E. | Robust.z | pval | padjusted |
| --- | --- | --- | --- | --- | --- | --- | --- |
| (Intercept) | 0.0188258 | 0.0227272 | 0.8283416 | 0.0284213 | 0.6623860 | 0.5077239 | 1 |
| SNF\_label\_2SNF\_1 | -0.0155635 | 0.0246393 | -0.6316552 | 0.0305083 | -0.5101399 | 0.6099534 | 1 |

GEE model with MCA dimension 4 as outcome

|  | Estimate | Naive.S.E. | Naive.z | Robust.S.E. | Robust.z | pval | padjusted |
| --- | --- | --- | --- | --- | --- | --- | --- |
| (Intercept) | -0.0105635 | 0.0228886 | -0.4615166 | 0.0132171 | -0.7992291 | 0.4241576 | 0.6414387 |
| SNF\_label\_2SNF\_1 | 0.0185885 | 0.0259024 | 0.7176346 | 0.0187199 | 0.9929807 | 0.3207193 | 0.6414387 |

GEE model with MCA dimension 5 as outcome

|  | Estimate | Naive.S.E. | Naive.z | Robust.S.E. | Robust.z | pval | padjusted |
| --- | --- | --- | --- | --- | --- | --- | --- |
| (Intercept) | -0.0116773 | 0.0159570 | -0.7317937 | 0.0197459 | -0.5913772 | 0.5542677 | 1 |
| SNF\_label\_2SNF\_1 | 0.0043620 | 0.0165868 | 0.2629777 | 0.0204496 | 0.2133021 | 0.8310913 | 1 |

GEE model with MCA dimension 6 as outcome

|  | Estimate | Naive.S.E. | Naive.z | Robust.S.E. | Robust.z | pval | padjusted |
| --- | --- | --- | --- | --- | --- | --- | --- |
| (Intercept) | -0.0029090 | 0.0158633 | -0.1833788 | 0.0148308 | -0.1961450 | 0.8444967 | 1 |
| SNF\_label\_2SNF\_1 | 0.0109413 | 0.0175065 | 0.6249866 | 0.0173375 | 0.6310807 | 0.5279877 | 1 |

GEE model with MCA dimension 7 as outcome

|  | Estimate | Naive.S.E. | Naive.z | Robust.S.E. | Robust.z | pval | padjusted |
| --- | --- | --- | --- | --- | --- | --- | --- |
| (Intercept) | 0.0125796 | 0.0146896 | 0.8563626 | 0.0193921 | 0.6486978 | 0.5165337 | 1 |
| SNF\_label\_2SNF\_1 | -0.0114091 | 0.0163272 | -0.6987802 | 0.0213200 | -0.5351362 | 0.5925557 | 1 |

GEE model with MCA dimension 8 as outcome

|  | Estimate | Naive.S.E. | Naive.z | Robust.S.E. | Robust.z | pval | padjusted |
| --- | --- | --- | --- | --- | --- | --- | --- |
| (Intercept) | 0.0008134 | 0.0146883 | 0.0553794 | 0.0174124 | 0.0467156 | 0.9627399 | 1 |
| SNF\_label\_2SNF\_1 | -0.0066509 | 0.0154496 | -0.4304912 | 0.0190934 | -0.3483361 | 0.7275878 | 1 |

GEE model with MCA dimension 9 as outcome

|  | Estimate | Naive.S.E. | Naive.z | Robust.S.E. | Robust.z | pval | padjusted |
| --- | --- | --- | --- | --- | --- | --- | --- |
| (Intercept) | 0.0049722 | 0.0146841 | 0.3386117 | 0.0162054 | 0.3068249 | 0.7589767 | 1 |
| SNF\_label\_2SNF\_1 | 0.0028418 | 0.0157545 | 0.1803826 | 0.0178397 | 0.1592982 | 0.8734340 | 1 |

GEE model with MCA dimension 10 as outcome

|  | Estimate | Naive.S.E. | Naive.z | Robust.S.E. | Robust.z | pval | padjusted |
| --- | --- | --- | --- | --- | --- | --- | --- |
| (Intercept) | -0.0167499 | 0.0147755 | -1.133627 | 0.0127523 | -1.313483 | 0.1890202 | 0.372601 |
| SNF\_label\_2SNF\_1 | 0.0192731 | 0.0163387 | 1.179601 | 0.0145831 | 1.321603 | 0.1863005 | 0.372601 |

```
sessionInfo()
```

```
## R version 4.0.2 (2020-06-22)
## Platform: x86_64-pc-linux-gnu (64-bit)
## Running under: Debian GNU/Linux bullseye/sid
## 
## Matrix products: default
## BLAS:   /usr/lib/x86_64-linux-gnu/openblas-pthread/libblas.so.3
## LAPACK: /usr/lib/x86_64-linux-gnu/openblas-pthread/libopenblasp-r0.3.10.so
## 
## locale:
##  [1] LC_CTYPE=en_US.UTF-8       LC_NUMERIC=C              
##  [3] LC_TIME=en_US.UTF-8        LC_COLLATE=en_US.UTF-8    
##  [5] LC_MONETARY=en_US.UTF-8    LC_MESSAGES=en_US.UTF-8   
##  [7] LC_PAPER=en_US.UTF-8       LC_NAME=C                 
##  [9] LC_ADDRESS=C               LC_TELEPHONE=C            
## [11] LC_MEASUREMENT=en_US.UTF-8 LC_IDENTIFICATION=C       
## 
## attached base packages:
## [1] stats     graphics  grDevices utils     datasets  methods   base     
## 
## other attached packages:
## [1] gee_4.13-25
## 
## loaded via a namespace (and not attached):
##  [1] digest_0.6.31   R6_2.5.1        jsonlite_1.8.4  evaluate_0.20  
##  [5] rlang_1.1.0     cachem_1.0.7    cli_3.6.1       jquerylib_0.1.4
##  [9] bslib_0.4.2     rmarkdown_2.21  tools_4.0.2     xfun_0.38      
## [13] yaml_2.3.7      fastmap_1.1.1   compiler_4.0.2  htmltools_0.5.5
## [17] knitr_1.42      sass_0.4.5
```

---

1. Radboud University Medical Center,↩︎
