## Additional File 8 for "A Multi-omics Data Analysis Workflow Packaged as a FAIR Digital Object": Additional File 8.html

snf\_analysis\_out


In [1]:

```
# Parameters
snf_matrix_path = "snf.csv"
phenotypes_covariates_path = "phenotype_covariates_data_hr.csv"
mca_dims_path = "cbcl_mca_coord.csv"
output_dir_plots = "output_curium"
outdir_df = "snf_analysis_out.csv"
```

### Similarity Network Fusion downstream analysis¶

Authors: Casper de Visser

#### Introduction¶

This notebook contains the downstream analysis of the fused sample similarity matrix that was constructed with SNF (Wang 2014). Spectral clustering is performed on these sample similarities and these clusters are compared with the behavioral data and phenotypic covariates.

In [2]:

```
import sys
import numpy as np
import pandas as pd
import matplotlib.pylab as plt
from sklearn.cluster import spectral_clustering
from sklearn.metrics import v_measure_score
import snf
from snf import metrics
import seaborn as sns
import scipy.stats as stats
import statsmodels.api as sm
import statsmodels
```

#### Input file paths¶

snf\_matrix\_path   
phenotypes\_covariates\_path   
metabolomics\_path   
mca\_dims\_path   
output\_dir\_plots

#### Load numpy array of fused network¶

In [3]:

```
# Load Pandas DataFrame of fused network
fused_df = pd.read_csv(snf_matrix_path, index_col=0)

# Convert to numpy array
fused_network = fused_df.to_numpy()
```

#### Perform spectral clustering on fused network¶

In [4]:

```
# Spectral clustering

# determine optimal number of clusters (estimated via an eigengap approach)
best, second = snf.get_n_clusters(fused_network)

# Perform spectral clustering on the fused network
labels = spectral_clustering(fused_network, n_clusters=best)
labels_second = spectral_clustering(fused_network, n_clusters=second)
```

In [5]:

```
# Functions


def sort_fused_network(fused_network, labels_array):
    # Make Pandas Dataframes
    df = pd.DataFrame(fused_network)
    df_labels = pd.DataFrame(labels_array)
    df_labels.columns = ["Label"]
    # sort label df
    df_labels = df_labels.sort_values(by=['Label'])
    # sort fused network df with sorted labels
    df = df.reindex(df_labels.index)
    df = df[df_labels.index]
    array = df.to_numpy()
    np.fill_diagonal(array, 0)
    return(array)

def make_heatmap(array, n_clusters):
    # Create heatmap
    heatmap = plt.imshow(array, cmap='hot', interpolation='nearest')

    # Set axis names, title etc.
    plt.xlabel('samples')
    plt.ylabel('samples') 
    cbar = plt.colorbar(heatmap)
    cbar.ax.set_ylabel('sample correlations', loc="top")
    plt.suptitle('Fused network: sample correlations\nNumber of clusters: {:.2f}'.format(round(n_clusters)))
    plt.show()

    return(plt)
    
# Sort Fused networks according to labels found by spectral clustering
sorted_fused_network_best = sort_fused_network(fused_network, labels)
sorted_fused_network_second = sort_fused_network(fused_network, labels_second)
```

In [6]:

```
make_heatmap(sorted_fused_network_best,  best)
```

Out[6]:

```
<module 'matplotlib.pylab' from '/opt/conda/lib/python3.9/site-packages/matplotlib/pylab.py'>
```

In [7]:

```
make_heatmap(sorted_fused_network_second, second)
```

Out[7]:

```
<module 'matplotlib.pylab' from '/opt/conda/lib/python3.9/site-packages/matplotlib/pylab.py'>
```

In [8]:

```
# Evaluation metrics

# Determine V-measure score (requiring true lables)
#v_score_1 = v_measure_score(labels, true_labels)
#v_score_2 = v_measure_score(labels_second, true_labels)

# Silhouette score
np.fill_diagonal(fused_network, 0)
sil = metrics.silhouette_score(fused_network, labels)
sil2 = metrics.silhouette_score(fused_network, labels_second)

# Affinity Z-score
zscore =  metrics.affinity_zscore(fused_network, labels)
zscore2 = metrics.affinity_zscore(fused_network, labels_second)
```

#### Compare SNF clusters to phenotype data¶

In [9]:

```
phenotypes_data = pd.read_csv(phenotypes_covariates_path , index_col=0) #phenotype_covariates_data.csv
```

In [10]:

```
# Phenotypes process out

phenotypes_data = pd.read_csv(phenotypes_covariates_path , index_col=0) #phenotype_covariates_data.csv
phenotypes_data = phenotypes_data[phenotypes_data.index.isin(fused_df.index)]


# Add cluster labels from SNF

SNF_label_best_colname =  "SNF_label_" + str(best)
SNF_label_second_colname =  "SNF_label_" + str(second)

phenotypes_data[SNF_label_best_colname] = labels
phenotypes_data[SNF_label_second_colname] = labels_second

for i in phenotypes_data.index:
    phenotypes_data.at[i, SNF_label_best_colname] =  "SNF_"+ str(phenotypes_data.at[i, SNF_label_best_colname])
    phenotypes_data.at[i, SNF_label_second_colname] =  "SNF_"+ str(phenotypes_data.at[i, SNF_label_second_colname])
```

#### Generate mosaic plots, comparing SNF clusters with phenotypic covariates¶

In [11]:

```
# Function to generate the mosaic plot

from statsmodels.graphics.mosaicplot import mosaic

colors = ['#e69F00', '#56b4e9', '#009e73', '#f0e442', '#0072b2', '#d55e00', '#cc79a7', '#000000']

def make_mosaic(df, col1, col2, out_dir):
    
    #Sort df on pheno value that is plotted against the clusters
    df = df.sort_values(by=[col1])
    
    
    #Adjust plot size to value counts
    number_of_pheno_values = len(df[col1].value_counts())
    number_of_clusters = len(df[col2].value_counts())
    
    
    if number_of_pheno_values < 2:
        print('No differences are observed in this phenotypic feature among the subjects')
    
    else:
    
        if number_of_pheno_values < 4:
            number_of_pheno_values = 4
    
        # Figure size
        fig, ax = plt.subplots(figsize=(number_of_pheno_values*2,number_of_clusters*1.5))      
        
        
        # Figure color palette
        props= {}
        e = 0
        a =  0.6 - number_of_clusters/10 
        for i in df[col1].unique():
            for j in df[col2].unique():
                props[(str(i), str(j))] = {'color': colors[e], 'alpha' : a}
                a += (0.6 - number_of_clusters/10)
            e += 1
            a = (0.6 - number_of_clusters/10)
        
        # Figure lables (percentages)
        labels_dict={}
        for i in df[col1].unique():
            for j in df[col2].unique():
                samples = len(df[(df[col1] == i) & (df[col2] == j)])
                percentage = round(samples/len(df.index) * 100, 2)
                labels_dict[(str(i), str(j))] = str(i) + '; ' + str(j) + '\n' + str(percentage) + '%'

        # Generate plot
        mosaic(df, 
               [col1, col2], 
               ax=ax, 
               axes_label=False,
               properties = props,
               labelizer = lambda k: labels_dict[k])
        plt.xlabel(col1, fontsize=20)
        plt.ylabel(col2, fontsize=20)
        plt.savefig(out_dir)
        plt.show()
        plt.close()

        return(plt)
```

In [12]:

```
make_mosaic(phenotypes_data, 'Age', SNF_label_best_colname, str(output_dir_plots) + 'Age_l1.png')
```

Out[12]:

```
<module 'matplotlib.pylab' from '/opt/conda/lib/python3.9/site-packages/matplotlib/pylab.py'>
```

In [13]:

```
make_mosaic(phenotypes_data, 'Age', SNF_label_second_colname, str(output_dir_plots) + 'Age_l2.png')
```

Out[13]:

```
<module 'matplotlib.pylab' from '/opt/conda/lib/python3.9/site-packages/matplotlib/pylab.py'>
```

In [14]:

```
make_mosaic(phenotypes_data, 'Sex', SNF_label_best_colname, str(output_dir_plots) + 'sex_l1.png')
```

Out[14]:

```
<module 'matplotlib.pylab' from '/opt/conda/lib/python3.9/site-packages/matplotlib/pylab.py'>
```

In [15]:

```
make_mosaic(phenotypes_data, 'Sex', SNF_label_second_colname, str(output_dir_plots) + 'sex_l2.png')
```

Out[15]:

```
<module 'matplotlib.pylab' from '/opt/conda/lib/python3.9/site-packages/matplotlib/pylab.py'>
```

In [16]:

```
make_mosaic(phenotypes_data, 'Sick', SNF_label_best_colname, str(output_dir_plots) + 'Sick_l1.png')
```

```
No differences are observed in this phenotypic feature among the subjects
```

In [17]:

```
make_mosaic(phenotypes_data, 'Sick', SNF_label_second_colname, str(output_dir_plots) + 'Sick_l2.png')
```

```
No differences are observed in this phenotypic feature among the subjects
```

In [18]:

```
make_mosaic(phenotypes_data, 'Menstruation', SNF_label_best_colname, str(output_dir_plots) + 'Menstruation_l.png')
```

```
No differences are observed in this phenotypic feature among the subjects
```

In [19]:

```
make_mosaic(phenotypes_data, 'Menstruation', SNF_label_second_colname, str(output_dir_plots) + 'Menstruation_l2.png')
```

```
No differences are observed in this phenotypic feature among the subjects
```

In [20]:

```
make_mosaic(phenotypes_data, 'Vitamines', SNF_label_best_colname, str(output_dir_plots) + 'Vitamines_l.png')
```

Out[20]:

```
<module 'matplotlib.pylab' from '/opt/conda/lib/python3.9/site-packages/matplotlib/pylab.py'>
```

In [21]:

```
make_mosaic(phenotypes_data, 'Vitamines', SNF_label_second_colname, str(output_dir_plots) + 'Vitamines_l2.png')
```

Out[21]:

```
<module 'matplotlib.pylab' from '/opt/conda/lib/python3.9/site-packages/matplotlib/pylab.py'>
```

#### Chi square testing for categorical variables¶

In [22]:

```
import scipy.stats
from scipy.stats import chi2

categorical_vars = ['Sex', 'Age', 'Vitamines']

#phenotypes_data_nonan_test = phenotypes_data[phenotypes_data['Sex'].notna()]

chisquare_best_list = []

for i in categorical_vars:
    row = []
    ct_table = pd.crosstab(phenotypes_data[i], phenotypes_data[SNF_label_best_colname])
    chi2_stat, p, dof, expected = scipy.stats.chi2_contingency(ct_table)
    row.append(i)
    row.append(round(chi2_stat, 4))
    row.append(round(p, 4))
    row.append(dof)
    chisquare_best_list.append(row)
    
a = np.array(chisquare_best_list)
df = pd.DataFrame(a[:,1:], index = a[:,0], columns = ['chi2 test', 'p_value', 'degrees of freedom'])
df
```

Out[22]:

|  | chi2 test | p\_value | degrees of freedom |
| --- | --- | --- | --- |
| Sex | 5.5137 | 0.0189 | 1 |
| Age | 7.0686 | 0.4218 | 7 |
| Vitamines | 0.4332 | 0.5104 | 1 |

In [23]:

```
chisquare_second_list = []

for i in categorical_vars:
    row = []
    ct_table = pd.crosstab(phenotypes_data[i], phenotypes_data[SNF_label_second_colname])
    chi2_stat, p, dof, expected = scipy.stats.chi2_contingency(ct_table)
    row.append(i)
    row.append(round(chi2_stat, 4))
    row.append(round(p, 4))
    row.append(dof)
    chisquare_second_list.append(row)
    
a = np.array(chisquare_second_list)
df = pd.DataFrame(a[:,1:], index = a[:,0], columns = ['chi2 test', 'p_value', 'degrees of freedom'])
df
```

Out[23]:

|  | chi2 test | p\_value | degrees of freedom |
| --- | --- | --- | --- |
| Sex | 0.8725 | 0.8321 | 3 |
| Age | 15.27 | 0.8092 | 21 |
| Vitamines | 5.7114 | 0.1265 | 3 |

### Compare clusters on MCA dimensions¶

In [24]:

```
# Load in MCA dimensions

mca_coordinates = pd.read_csv(mca_dims_path, index_col=0)
mca_coordinates = mca_coordinates[mca_coordinates.index.isin(phenotypes_data.index)]
phenotypes_data = phenotypes_data[phenotypes_data.index.isin(mca_coordinates.index)]


# Add cluster labels from SNF

mca_coordinates[SNF_label_best_colname] = phenotypes_data[SNF_label_best_colname]
mca_coordinates[SNF_label_second_colname] = phenotypes_data[SNF_label_second_colname]


for i in mca_coordinates.index:
    mca_coordinates.at[i, SNF_label_best_colname] =  str(mca_coordinates.at[i, SNF_label_best_colname])
    mca_coordinates.at[i, SNF_label_second_colname] =  str(mca_coordinates.at[i, SNF_label_second_colname])

mca_coordinates.columns = mca_coordinates.columns.str.replace(' ', '_')
```

#### Functions used for statistics¶

In [25]:

```
def make_significant_bold(x):
    bold = 'bold' if x < 0.05 else ''
    return 'font-weight: %s' % bold


def make_pvalue_table(p_value_list):
    a = np.array(p_value_list)
    df = pd.DataFrame(a[:,1:], index = a[:,0], columns = ['test statistic', 'p-value'])
    df['test statistic'] = pd.to_numeric(df['test statistic'])
    df['p-value'] = pd.to_numeric(df['p-value'])
    p_values = np.asarray(df['p-value'].values.tolist())
    corrected_p_values = statsmodels.stats.multitest.fdrcorrection(p_values)
    df['FDR corrected p-value'] = corrected_p_values[1].tolist()
    df.style.applymap(make_significant_bold)
    return(df)


def man_whitney(group_list, df):
    
    # Calculate Mann-Whitney U tests per MCA dimension
    p_value_list = []
    
    for i in df.columns[0:-2]:
        row = []
        statistics = stats.mannwhitneyu(group_list[0][i], group_list[1][i])
        row.append(i)
        row.append(list(statistics)[0])
        row.append(list(statistics)[1])
        p_value_list.append(row)
        
    # Make nicer looking table for p-values
    df = make_pvalue_table(p_value_list)
    return(df)


def kruskal_wallis(groups_list, df):

    # Make list of the different snf groups, per mca dimension        
    groups_vs_mca_dims = []

    for i in df.columns[0:-2]:
        
        groups_vs_mca_dim = []
        for j in groups_list:
            groups_vs_mca_dim.append(j[i])
        groups_vs_mca_dims.append(groups_vs_mca_dim)
    
    # Calculate Kruskal-Wallis statistic over the different groups
    p_value_list = []
    dim_num = 0
    
    for i in groups_vs_mca_dims:
        dim_num += 1
        row = []
        statistics = stats.kruskal(*i)
        row.append("Dim_" + str(dim_num))
        row.append(list(statistics)[0])
        row.append(list(statistics)[1])
        p_value_list.append(row)
        
    # Make nice looking table
    df = make_pvalue_table(p_value_list)
    return(df)


def perform_statistical_testing(n_clusters, df):
    
    # Make list of groups of different SNF clusters (best or second)
    snf_groups = []
    
    for i in range(n_clusters):
        snf_label = 'SNF_' + str(i)
        snf_groups.append(df[df["SNF_label_" + str(n_clusters)] == snf_label])
    
    # Perform statistical test based on number of clusters
    if len(snf_groups) > 2:
        p_values = kruskal_wallis(snf_groups, df)

    else:
        p_values = man_whitney(snf_groups, df)
                            
    return(p_values)


def perform_statistical_testing_aggression(n_clusters, df):

    snf_groups = []

    for i in range(n_clusters):
        snf_label = 'SNF_' + str(i)
        group = df[df["SNF_label_" + str(n_clusters)] == snf_label]
        snf_groups.append(group["Aggression_Tscore"])


    if len(snf_groups) > 2:
        for i in snf_groups:
            p_values = stats.kruskal(*i)
    
    else:
        p_values = stats.mannwhitneyu(snf_groups[0], snf_groups[1])
        
    return(p_values)
```

#### Shapiro test for MCA dimensions¶

In [26]:

```
# Shapiro tests show that MCA dimensions are not normalliy distributed
p_value_list = []

for i in mca_coordinates.columns[0:-2]:
    row = []
    statistics = stats.shapiro(mca_coordinates[i])
    row.append(i)
    row.append(list(statistics)[0])
    row.append(list(statistics)[1])
    p_value_list.append(row)

df = make_pvalue_table(p_value_list)
df.style.applymap(make_significant_bold)
df
```

Out[26]:

|  | test statistic | p-value | FDR corrected p-value |
| --- | --- | --- | --- |
| Dim\_1 | 0.964185 | 2.926199e-04 | 9.753997e-04 |
| Dim\_2 | 0.889399 | 9.429951e-10 | 9.429951e-09 |
| Dim\_3 | 0.995970 | 9.380884e-01 | 9.380884e-01 |
| Dim\_4 | 0.988087 | 1.760247e-01 | 2.851989e-01 |
| Dim\_5 | 0.985788 | 9.133694e-02 | 2.075845e-01 |
| Dim\_6 | 0.986232 | 1.037922e-01 | 2.075845e-01 |
| Dim\_7 | 0.988929 | 2.225651e-01 | 2.851989e-01 |
| Dim\_8 | 0.921603 | 8.790787e-08 | 4.395394e-07 |
| Dim\_9 | 0.989019 | 2.281592e-01 | 2.851989e-01 |
| Dim\_10 | 0.992731 | 5.772696e-01 | 6.414107e-01 |

In [27]:

```
from IPython.display import display, Markdown
display(Markdown("## Comparing " + str(best) + " clusters on MCA dimensions"))
```

#### Comparing 2 clusters on MCA dimensions¶

In [28]:

```
results = perform_statistical_testing(best, mca_coordinates)
results.style.applymap(make_significant_bold)
if best > 2: 
    print('Kruskal-Wallis test\n\n') 
else: 
    print('Mann-Whitney U test\n\n')
results
```

```
Mann-Whitney U test
```

Out[28]:

|  | test statistic | p-value | FDR corrected p-value |
| --- | --- | --- | --- |
| Dim\_1 | 3329.0 | 0.825727 | 0.825727 |
| Dim\_2 | 3485.0 | 0.775318 | 0.825727 |
| Dim\_3 | 3704.0 | 0.317406 | 0.555754 |
| Dim\_4 | 3219.0 | 0.562586 | 0.773585 |
| Dim\_5 | 3727.0 | 0.282451 | 0.555754 |
| Dim\_6 | 2603.0 | 0.009642 | 0.096423 |
| Dim\_7 | 2839.0 | 0.068978 | 0.229925 |
| Dim\_8 | 2701.0 | 0.023285 | 0.116426 |
| Dim\_9 | 3100.0 | 0.333452 | 0.555754 |
| Dim\_10 | 3550.0 | 0.618868 | 0.773585 |

In [29]:

```
display(Markdown("## Comparing " + str(second) + " clusters on MCA dimensions"))
```

#### Comparing 4 clusters on MCA dimensions¶

In [30]:

```
results = perform_statistical_testing(second, mca_coordinates)
results.style.applymap(make_significant_bold)
if second > 2: 
    print('Kruskal-Wallis test\n\n') 
else: 
    print('Mann-Whitney U test\n\n')
results
```

```
Kruskal-Wallis test
```

Out[30]:

|  | test statistic | p-value | FDR corrected p-value |
| --- | --- | --- | --- |
| Dim\_1 | 0.420380 | 0.936001 | 0.936001 |
| Dim\_2 | 0.607391 | 0.894738 | 0.936001 |
| Dim\_3 | 2.268283 | 0.518624 | 0.864374 |
| Dim\_4 | 7.404629 | 0.060060 | 0.200201 |
| Dim\_5 | 0.931893 | 0.817725 | 0.936001 |
| Dim\_6 | 11.703935 | 0.008469 | 0.071746 |
| Dim\_7 | 2.454744 | 0.483525 | 0.864374 |
| Dim\_8 | 10.561554 | 0.014349 | 0.071746 |
| Dim\_9 | 1.216769 | 0.748985 | 0.936001 |
| Dim\_10 | 5.467877 | 0.140573 | 0.351432 |

#### Aggression T-scores¶

In [31]:

```
# Remove samples with NA values for Aggression score

phenotypes_data_nonan = phenotypes_data[phenotypes_data['Aggression_Tscore'].notna()]

# Perform shapiro test for normality
shapiro = stats.shapiro(phenotypes_data_nonan["Aggression_Tscore"])

if float(list(shapiro)[1]) < 0.05:
    print('Shapiro test shows that Aggression T-scores are not normally distributed')
    print('p-value', list(shapiro)[1])
    
else:
    print('Shapiro test shows that Aggression T-scores are normally distributed')
    print('p-value', list(shapiro)[1])
```

```
Shapiro test shows that Aggression T-scores are not normally distributed
p-value 0.0002661454782355577
```

In [32]:

```
#Compare SNF clustering to Aggression T-scores

results = perform_statistical_testing_aggression(best, phenotypes_data_nonan)
results_second = perform_statistical_testing_aggression(second, phenotypes_data_nonan)

print('Comparing best SNF clustering on Aggresion T-score\n')
if best > 2: 
    print('Kruskal-Wallis test\n') 
else: 
    print('Mann-Whitney U test\n')
print('p-value:' , str(list(results)[1]))
print('\n\nComparing second best SNF clustering on Aggresion T-score\n')
if second > 2: 
    print('Kruskal-Wallis test\n') 
else: 
    print('Mann-Whitney U test\n')
print('p-value:' + str(list(results_second)[1]))
```

```
Comparing best SNF clustering on Aggresion T-score

Mann-Whitney U test

p-value: 0.9817624226188104


Comparing second best SNF clustering on Aggresion T-score

Kruskal-Wallis test

p-value:0.46310474709968097
```

#### Write Dataframe to csv for GEE model¶

In [33]:

```
mca_coordinates = mca_coordinates.join(phenotypes_data_nonan["Aggression_Tscore"])
mca_coordinates = mca_coordinates.join(phenotypes_data_nonan["Age"])
mca_coordinates = mca_coordinates.join(phenotypes_data_nonan["Sex"])
mca_coordinates = mca_coordinates.join(phenotypes_data_nonan["Vitamines"])
if 'Familynumber' in phenotypes_data_nonan.columns:
    mca_coordinates = mca_coordinates.join(phenotypes_data_nonan["Familynumber"])
mca_coordinates.to_csv(outdir_df)
```
