## Additional File 7 for "A Multi-omics Data Analysis Workflow Packaged as a FAIR Digital Object": Additional File 7.html

snf\_analysis\_out


In [1]:

```
# Parameters
snf_matrix_path = "snf.csv"
phenotypes_covariates_path = "phenotype_covariates_data_hr.csv"
mca_dims_path = "cbcl_mca_coord.csv"
output_dir_plots = "output_ntr"
outdir_df = "snf_analysis_out.csv"
```

In [12]:

```
make_mosaic(phenotypes_data, 'Age', SNF_label_best_colname, str(output_dir_plots) + 'Age_l1.png')
```

Out[12]:

```
<module 'matplotlib.pylab' from '/opt/conda/lib/python3.9/site-packages/matplotlib/pylab.py'>
```

In [13]:

```
make_mosaic(phenotypes_data, 'Age', SNF_label_second_colname, str(output_dir_plots) + 'Age_l2.png')
```

Out[13]:

```
<module 'matplotlib.pylab' from '/opt/conda/lib/python3.9/site-packages/matplotlib/pylab.py'>
```

In [14]:

```
make_mosaic(phenotypes_data, 'Sex', SNF_label_best_colname, str(output_dir_plots) + 'sex_l1.png')
```

Out[14]:

```
<module 'matplotlib.pylab' from '/opt/conda/lib/python3.9/site-packages/matplotlib/pylab.py'>
```

In [15]:

```
make_mosaic(phenotypes_data, 'Sex', SNF_label_second_colname, str(output_dir_plots) + 'sex_l2.png')
```

Out[15]:

```
<module 'matplotlib.pylab' from '/opt/conda/lib/python3.9/site-packages/matplotlib/pylab.py'>
```

In [16]:

```
make_mosaic(phenotypes_data, 'Sick', SNF_label_best_colname, str(output_dir_plots) + 'Sick_l1.png')
```

```
No differences are observed in this phenotypic feature among the subjects
```

In [17]:

```
make_mosaic(phenotypes_data, 'Sick', SNF_label_second_colname, str(output_dir_plots) + 'Sick_l2.png')
```

```
No differences are observed in this phenotypic feature among the subjects
```

In [18]:

```
make_mosaic(phenotypes_data, 'Menstruation', SNF_label_best_colname, str(output_dir_plots) + 'Menstruation_l.png')
```

```
No differences are observed in this phenotypic feature among the subjects
```

In [19]:

```
make_mosaic(phenotypes_data, 'Menstruation', SNF_label_second_colname, str(output_dir_plots) + 'Menstruation_l2.png')
```

```
No differences are observed in this phenotypic feature among the subjects
```

In [20]:

```
make_mosaic(phenotypes_data, 'Vitamines', SNF_label_best_colname, str(output_dir_plots) + 'Vitamines_l.png')
```

Out[20]:

```
<module 'matplotlib.pylab' from '/opt/conda/lib/python3.9/site-packages/matplotlib/pylab.py'>
```

In [21]:

```
make_mosaic(phenotypes_data, 'Vitamines', SNF_label_second_colname, str(output_dir_plots) + 'Vitamines_l2.png')
```

Out[21]:

```
<module 'matplotlib.pylab' from '/opt/conda/lib/python3.9/site-packages/matplotlib/pylab.py'>
```

#### Chi square testing for categorical variables¶

In [22]:

```
import scipy.stats
from scipy.stats import chi2

categorical_vars = ['Sex', 'Age', 'Vitamines']

#phenotypes_data_nonan_test = phenotypes_data[phenotypes_data['Sex'].notna()]

chisquare_best_list = []

for i in categorical_vars:
    row = []
    ct_table = pd.crosstab(phenotypes_data[i], phenotypes_data[SNF_label_best_colname])
    chi2_stat, p, dof, expected = scipy.stats.chi2_contingency(ct_table)
    row.append(i)
    row.append(round(chi2_stat, 4))
    row.append(round(p, 4))
    row.append(dof)
    chisquare_best_list.append(row)
    
a = np.array(chisquare_best_list)
df = pd.DataFrame(a[:,1:], index = a[:,0], columns = ['chi2 test', 'p_value', 'degrees of freedom'])
df
```

Out[22]:

|  | chi2 test | p\_value | degrees of freedom |
| --- | --- | --- | --- |
| Sex | 2.5857 | 0.46 | 3 |
| Age | 31.2439 | 0.0697 | 21 |
| Vitamines | 0.8001 | 0.8494 | 3 |

In [23]:

```
chisquare_second_list = []

for i in categorical_vars:
    row = []
    ct_table = pd.crosstab(phenotypes_data[i], phenotypes_data[SNF_label_second_colname])
    chi2_stat, p, dof, expected = scipy.stats.chi2_contingency(ct_table)
    row.append(i)
    row.append(round(chi2_stat, 4))
    row.append(round(p, 4))
    row.append(dof)
    chisquare_second_list.append(row)
    
a = np.array(chisquare_second_list)
df = pd.DataFrame(a[:,1:], index = a[:,0], columns = ['chi2 test', 'p_value', 'degrees of freedom'])
df
```

Out[23]:

|  | chi2 test | p\_value | degrees of freedom |
| --- | --- | --- | --- |
| Sex | 2.8365 | 0.0921 | 1 |
| Age | 23.0853 | 0.0016 | 7 |
| Vitamines | 0.0432 | 0.8354 | 1 |

### Compare clusters on MCA dimensions¶

In [24]:

```
# Load in MCA dimensions

mca_coordinates = pd.read_csv(mca_dims_path, index_col=0)
mca_coordinates = mca_coordinates[mca_coordinates.index.isin(phenotypes_data.index)]
phenotypes_data = phenotypes_data[phenotypes_data.index.isin(mca_coordinates.index)]


# Add cluster labels from SNF

mca_coordinates[SNF_label_best_colname] = phenotypes_data[SNF_label_best_colname]
mca_coordinates[SNF_label_second_colname] = phenotypes_data[SNF_label_second_colname]


for i in mca_coordinates.index:
    mca_coordinates.at[i, SNF_label_best_colname] =  str(mca_coordinates.at[i, SNF_label_best_colname])
    mca_coordinates.at[i, SNF_label_second_colname] =  str(mca_coordinates.at[i, SNF_label_second_colname])

mca_coordinates.columns = mca_coordinates.columns.str.replace(' ', '_')
```

Out[26]:

|  | test statistic | p-value | FDR corrected p-value |
| --- | --- | --- | --- |
| Dim\_1 | 0.781586 | 2.509019e-33 | 3.136274e-33 |
| Dim\_2 | 0.528872 | 4.904545e-44 | 2.452272e-43 |
| Dim\_3 | 0.740012 | 1.283412e-35 | 3.208530e-35 |
| Dim\_4 | 0.195504 | 0.000000e+00 | 0.000000e+00 |
| Dim\_5 | 0.817525 | 4.784094e-31 | 5.315660e-31 |
| Dim\_6 | 0.827764 | 2.478981e-30 | 2.478981e-30 |
| Dim\_7 | 0.596048 | 9.076210e-42 | 3.025403e-41 |
| Dim\_8 | 0.757255 | 1.047688e-34 | 2.095375e-34 |
| Dim\_9 | 0.781468 | 2.469075e-33 | 3.136274e-33 |
| Dim\_10 | 0.763065 | 2.183649e-34 | 3.639414e-34 |

In [27]:

```
from IPython.display import display, Markdown
display(Markdown("## Comparing " + str(best) + " clusters on MCA dimensions"))
```

#### Comparing 4 clusters on MCA dimensions¶

In [28]:

```
results = perform_statistical_testing(best, mca_coordinates)
results.style.applymap(make_significant_bold)
if best > 2: 
    print('Kruskal-Wallis test\n\n') 
else: 
    print('Mann-Whitney U test\n\n')
results
```

```
Kruskal-Wallis test
```

Out[28]:

|  | test statistic | p-value | FDR corrected p-value |
| --- | --- | --- | --- |
| Dim\_1 | 0.916453 | 0.821456 | 0.972034 |
| Dim\_2 | 0.233348 | 0.972034 | 0.972034 |
| Dim\_3 | 0.269599 | 0.965641 | 0.972034 |
| Dim\_4 | 4.191639 | 0.241500 | 0.643860 |
| Dim\_5 | 5.343192 | 0.148324 | 0.643860 |
| Dim\_6 | 0.638678 | 0.887524 | 0.972034 |
| Dim\_7 | 0.595839 | 0.897384 | 0.972034 |
| Dim\_8 | 1.560763 | 0.668319 | 0.972034 |
| Dim\_9 | 4.036553 | 0.257544 | 0.643860 |
| Dim\_10 | 6.401652 | 0.093623 | 0.643860 |

In [29]:

```
display(Markdown("## Comparing " + str(second) + " clusters on MCA dimensions"))
```

#### Comparing 2 clusters on MCA dimensions¶

In [30]:

```
results = perform_statistical_testing(second, mca_coordinates)
results.style.applymap(make_significant_bold)
if second > 2: 
    print('Kruskal-Wallis test\n\n') 
else: 
    print('Mann-Whitney U test\n\n')
results
```

```
Mann-Whitney U test
```

Out[30]:

|  | test statistic | p-value | FDR corrected p-value |
| --- | --- | --- | --- |
| Dim\_1 | 78687.5 | 0.595088 | 0.814794 |
| Dim\_2 | 74071.0 | 0.413525 | 0.814794 |
| Dim\_3 | 80625.5 | 0.272225 | 0.814794 |
| Dim\_4 | 77595.0 | 0.832104 | 0.832104 |
| Dim\_5 | 78899.5 | 0.553019 | 0.814794 |
| Dim\_6 | 77686.5 | 0.811293 | 0.832104 |
| Dim\_7 | 79550.0 | 0.433426 | 0.814794 |
| Dim\_8 | 75324.5 | 0.651835 | 0.814794 |
| Dim\_9 | 74204.0 | 0.436208 | 0.814794 |
| Dim\_10 | 81765.5 | 0.152489 | 0.814794 |

#### Aggression T-scores¶

In [31]:

```
# Remove samples with NA values for Aggression score

phenotypes_data_nonan = phenotypes_data[phenotypes_data['Aggression_Tscore'].notna()]

# Perform shapiro test for normality
shapiro = stats.shapiro(phenotypes_data_nonan["Aggression_Tscore"])

if float(list(shapiro)[1]) < 0.05:
    print('Shapiro test shows that Aggression T-scores are not normally distributed')
    print('p-value', list(shapiro)[1])
    
else:
    print('Shapiro test shows that Aggression T-scores are normally distributed')
    print('p-value', list(shapiro)[1])
```

```
Shapiro test shows that Aggression T-scores are not normally distributed
p-value 2.5653841258334235e-29
```

In [32]:

```
#Compare SNF clustering to Aggression T-scores

results = perform_statistical_testing_aggression(best, phenotypes_data_nonan)
results_second = perform_statistical_testing_aggression(second, phenotypes_data_nonan)

print('Comparing best SNF clustering on Aggresion T-score\n')
if best > 2: 
    print('Kruskal-Wallis test\n') 
else: 
    print('Mann-Whitney U test\n')
print('p-value:' , str(list(results)[1]))
print('\n\nComparing second best SNF clustering on Aggresion T-score\n')
if second > 2: 
    print('Kruskal-Wallis test\n') 
else: 
    print('Mann-Whitney U test\n')
print('p-value:' + str(list(results_second)[1]))
```

```
Comparing best SNF clustering on Aggresion T-score

Kruskal-Wallis test

p-value: 0.4835674963574282


Comparing second best SNF clustering on Aggresion T-score

Mann-Whitney U test

p-value:0.7009162273494289
```

#### Write Dataframe to csv for GEE model¶

In [33]:

```
mca_coordinates = mca_coordinates.join(phenotypes_data_nonan["Aggression_Tscore"])
mca_coordinates = mca_coordinates.join(phenotypes_data_nonan["Age"])
mca_coordinates = mca_coordinates.join(phenotypes_data_nonan["Sex"])
mca_coordinates = mca_coordinates.join(phenotypes_data_nonan["Vitamines"])
if 'Familynumber' in phenotypes_data_nonan.columns:
    mca_coordinates = mca_coordinates.join(phenotypes_data_nonan["Familynumber"])
mca_coordinates.to_csv(outdir_df)
```
