## Additional File 4 for "A Multi-omics Data Analysis Workflow Packaged as a FAIR Digital Object": Additional File 4.html

MOFA downstream analysis report


### MOFA downstream analysis report

###### Purva Kulkarni^1^

###### Casper de Visser^2^

###### Anna Niehues^3^

#### Introduction

This notebook contains the downstream analysis steps and
visualizations in the form of plots generated using a MOFA (Multi-omics
Factor Analysis) model. The text explaining these analyses steps is
adapted from “MOFA+: downstream analysis (in R)” by Ricard Argelaguet
and Britta Velten (2020).

#### List input parameters

```
print(params)
```

```
## $MOFA_factor_top100_epigenomics_weights_outdir
## [1] "MOFA_factor_top100_epigenomics_weights"
## 
## $modelFile
## [1] "MOFAmodel.hdf5"
## 
## $metaData
## [1] "phenotype_covariates_data_hr.csv"
## 
## $mcaCoord
## [1] "cbcl_mca_coord.csv"
## 
## $epic_anno
## [1] "epigenomics_annotation_out.RData"
```

#### Load libraries

```
library(ggplot2)
library(MOFA2)
```

#### Load the mofa model

```
model <- load_model(params$modelFile)
```

```
## Warning in .quality_control(object, verbose = verbose): Factor(s) 1 are strongly correlated with the total number of expressed features for at least one of your omics. Such factors appear when there are differences in the total 'levels' between your samples, *sometimes* because of poor normalisation in the preprocessing steps.
```

#### Read the meta data file

```
metaData <- read.csv(params$metaData)
#head(metaData)
```

#### Load DNA methylation annotation file

```
load(params$epic_anno)
```

#### Read the file resulting from the multiple correspondance analysis

```
cbcl_mca_coord <- read.csv(params$mcaCoord)
#head(cbcl_mca_coord)
```

#### Plot data overview

Obtain an overview of the input omics data. It shows how many views
(rows) and how many groups (columns) exist, what are their corresponding
dimensionalities and how many missing information they have (grey
bars).

```
plot_data_overview(model)
```

#### Add meta data to the model

The metadata is stored as a data.frame object in model@samples\_metadata,
and it requires at least the column sample

```
names(metaData)[names(metaData) == 'X'] <- 'sample' 
metaData_cor <- metaData
metaData <- metaData[metaData$sample %in% colnames(model@data$metabolomics$group1),]
samples_metadata(model) <- metaData
```

#### Variance decomposition

This step quantifies the amount of variance explained (\(R^2\)) by each factor in each data
modality.

##### Total variance explained per view

```
knitr::kable(head(get_variance_explained(model)$r2_total[[1]]))
```

|  | x |
| --- | --- |
| epigenomics | 75.909562 |
| metabolomics | 1.897284 |

##### Variance explained for every factor in per view

```
knitr::kable(head(get_variance_explained(model)$r2_per_factor[[1]]))
```

|  | epigenomics | metabolomics |
| --- | --- | --- |
| Factor1 | 72.5644570 | 0.0000305 |
| Factor2 | 0.1024144 | 1.8769112 |
| Factor3 | 0.9012006 | 0.0002667 |
| Factor4 | 0.5126855 | 0.0054320 |
| Factor5 | 0.4224853 | 0.0080295 |
| Factor6 | 0.4241417 | 0.0029281 |

##### Plot the variance explained estimates

```
plot_variance_explained(model, x="view", y="factor") +
  # add explained variance values
  geom_text(label = round(c(
    get_variance_explained(model)$r2_per_factor$group1), 5))
```

```
plot_variance_explained(model, x="view", y="factor", plot_total = TRUE)[[2]]
```

#### Visualisations of factors

The MOFA factors capture the global sources of variability in the
data. Mathematically, each factor ordinates cells along a
one-dimensional axis centered at zero. The value per se is not
important, only the relative positioning of samples matters. Samples
with different signs manifest opposite “effects” along the inferred axis
of variation, with higher absolute value indicating a stronger
effect.

##### Plot factors

Individual factors can be plotted and visualized in combination with
other factors in the form of scatter plots and colored based on specific
condition, for example, in the plot below factors 1 to 3 have been
visualized and are colored based on Sex.

```
 plot_factors(model, factors = 1:3, color_by = "Sex")
```

```
## Registered S3 method overwritten by 'GGally':
##   method from   
##   +.gg   ggplot2
```

##### Plot factor correlations

```
plot_factor_cor(model)
```

#### Visualisation of feature weights

The weights provide a score for how strong each feature relates to
each factor. Features with no association with the factor have values
close to zero, while features with strong association with the factor
have large absolute values. The sign of the weight indicates the
direction of the effect: a positive weight indicates that the feature
has higher levels in the cells with positive factor values, and vice
versa.

##### Plot top weights per factor epigenomics

Display highest loading features Tables below each plot show CpG
locations on the genome

```
require(knitr)
```

```
## Loading required package: knitr
```

```
dir.create(params$MOFA_factor_top100_epigenomics_weights_outdir)

for (MOFA_factor in 1:10) {
  # plot top10 weights
  plot(plot_top_weights(model, factors = MOFA_factor, view = 1))
  # retrieve weights
  df_weights <- get_weights(
    model, factors = MOFA_factor, view = 1, abs = TRUE, as.data.frame = TRUE)
  row.names(df_weights) <- df_weights$feature
  # sort weights
  df_weights <- df_weights[order(df_weights$value, decreasing = TRUE), ]
  # get EPIC annotations
  columns <- c("chromosome", "position", "gene.symbol", "gene.accession")
  mapped_CpG_sites <- epicanno[rownames(df_weights)[1:100], columns]
  mapped_CpG_sites$factor.weight <- df_weights$value[1:100]
  # save top100 weights to csv
  write.csv(mapped_CpG_sites,
            file.path(params$MOFA_factor_top100_epigenomics_weights_outdir,
                      paste0("factor", MOFA_factor, "_weights.csv")))
  # display top10 weights
  print(knitr::kable(mapped_CpG_sites[1:10, ]))
  cat("\n\n\n")
}
```

|  | chromosome | position | gene.symbol | gene.accession | factor.weight |
| --- | --- | --- | --- | --- | --- |
| cg02787560 | chr16 | 3998777 |  |  | 0.2982292 |
| cg13340765 | chr8 | 142276287 |  |  | 0.2902006 |
| cg04770195 | chr16 | 53114571 | CHD9 | NM\_025134 | 0.2891673 |
| cg13964405 | chr1 | 29009447 | GMEB1;GMEB1 | NM\_024482;NM\_006582 | 0.2874957 |
| cg24716664 | chr11 | 1006121 | AP2A2 | NM\_012305 | 0.2865425 |
| cg14431175 | chr17 | 35806368 | TADA2A;TADA2A;TADA2A;TADA2A | NM\_133439;NM\_001291918;NM\_001166105;NM\_001488 | 0.2852947 |
| cg26084376 | chr5 | 127185815 |  |  | 0.2846209 |
| cg06715204 | chr3 | 171594651 | TMEM212-AS1 | NR\_046852 | 0.2846123 |
| cg04737166 | chr9 | 34372878 | KIAA1161 | NM\_020702 | 0.2829073 |
| cg23884120 | chr12 | 121198406 |  |  | 0.2812711 |

|  | chromosome | position | gene.symbol | gene.accession | factor.weight |
| --- | --- | --- | --- | --- | --- |
| cg21923568 | chr7 | 98721688 | SMURF1;SMURF1;SMURF1 | NM\_001199847;NM\_020429;NM\_181349 | 4.082707 |
| cg13355472 | chr6 | 17431988 | CAP2 | NM\_006366 | 3.647331 |
| cg16942644 | chr21 | 46867867 | COL18A1 | NM\_130445 | 3.572141 |
| cg07729025 | chr16 | 56883034 |  |  | 3.566086 |
| cg11873492 | chr4 | 84140721 |  |  | 3.471508 |
| cg18615437 | chr10 | 106094821 | ITPRIP;ITPRIP | NM\_033397;NM\_001272013 | 3.437669 |
| cg04346791 | chr12 | 32607254 |  |  | 3.274109 |
| cg14696311 | chr13 | 114855198 | RASA3 | NM\_007368 | 3.234331 |
| cg15269794 | chr5 | 68824022 | OCLN;OCLN;OCLN | NM\_001205254;NM\_002538;NM\_001205255 | 3.226033 |
| cg05210373 | chr14 | 105173786 | INF2;INF2 | NM\_022489;NM\_001031714 | 3.214714 |

|  | chromosome | position | gene.symbol | gene.accession | factor.weight |
| --- | --- | --- | --- | --- | --- |
| cg11209549 | chr1 | 223314605 | TLR5 | NM\_003268 | 0.6307081 |
| cg21617353 | chr17 | 43319371 | FMNL1 | NM\_005892 | 0.6053237 |
| cg24290948 | chr3 | 12800802 | TMEM40;TMEM40 | NM\_018306;NM\_018306 | 0.5900655 |
| cg19452633 | chr17 | 42164476 | HDAC5;HDAC5 | NM\_001015053;NM\_005474 | 0.5727998 |
| cg09268718 | chr17 | 1546403 | SCARF1;SCARF1;SCARF1;SCARF1;SCARF1 | NM\_145352;NR\_028075;NM\_003693;NR\_028076;NM\_145350 | 0.5682427 |
| cg19917235 | chr5 | 131594961 | PDLIM4;PDLIM4 | NM\_001131027;NM\_003687 | 0.5624443 |
| cg06511312 | chr1 | 3459949 | MEGF6 | NM\_001409 | 0.5462586 |
| cg19955173 | chr3 | 195603697 | TNK2;TNK2 | NM\_001010938;NM\_005781 | 0.5391736 |
| cg09490371 | chr2 | 233253024 | ECEL1P2 | NR\_028501 | 0.5288945 |
| cg20408776 | chr17 | 43319382 | FMNL1 | NM\_005892 | 0.5286901 |

|  | chromosome | position | gene.symbol | gene.accession | factor.weight |
| --- | --- | --- | --- | --- | --- |
| cg20695297 | chr19 | 3178844 | S1PR4 | NM\_003775 | 2.194112 |
| cg08482436 | chr7 | 96647888 |  |  | 1.791103 |
| cg00011225 | chr2 | 219738314 | WNT6 | NM\_006522 | 1.787890 |
| cg00200734 | chr9 | 970379 |  |  | 1.733350 |
| cg13577556 | chr9 | 975059 |  |  | 1.658289 |
| cg24792289 | chr5 | 134825895 |  |  | 1.654354 |
| cg08854008 | chr1 | 9714397 | C1orf200;PIK3CD | NR\_027045;NM\_005026 | 1.649250 |
| cg22344703 | chr2 | 219736312 | WNT6 | NM\_006522 | 1.625491 |
| cg00548708 | chr17 | 48041547 |  |  | 1.620436 |
| cg00211208 | chr16 | 88770157 | RNF166;RNF166;RNF166 | NM\_001171816;NM\_178841;NM\_001171815 | 1.617484 |

|  | chromosome | position | gene.symbol | gene.accession | factor.weight |
| --- | --- | --- | --- | --- | --- |
| cg08058408 | chr14 | 64557064 | SYNE2;SYNE2 | NM\_182914;NM\_015180 | 3.682701 |
| cg03652396 | chr3 | 194425225 |  |  | 3.584276 |
| cg15556865 | chr11 | 94202317 | MRE11A;MRE11A | NM\_005591;NM\_005590 | 3.575799 |
| cg16626437 | chr6 | 111799485 | REV3L;REV3L;REV3L | NM\_002912;NM\_001286431;NM\_001286432 | 3.351998 |
| cg04695939 | chr3 | 43573746 | ANO10;ANO10;ANO10;ANO10;ANO10 | NM\_001204834;NM\_001204833;NM\_001204832;NM\_018075;NM\_001204831 | 3.343796 |
| cg06019956 | chr1 | 207213607 |  |  | 3.343628 |
| cg21546601 | chr1 | 31982773 | LOC284551 | NR\_027085 | 3.332251 |
| cg22939408 | chr20 | 13070768 | SPTLC3 | NM\_018327 | 3.307591 |
| cg13333107 | chr12 | 15931214 | EPS8 | NM\_004447 | 3.223621 |
| cg05736186 | chr11 | 7633488 | PPFIBP2 | NM\_003621 | 3.220366 |

|  | chromosome | position | gene.symbol | gene.accession | factor.weight |
| --- | --- | --- | --- | --- | --- |
| cg18598642 | chr4 | 152497130 | FAM160A1 | NM\_001109977 | 4.061499 |
| cg08343295 | chr4 | 41162279 | APBB2;APBB2;APBB2 | NM\_173075;NM\_001166050;NM\_004307 | 3.796105 |
| cg08670693 | chr4 | 41162197 | APBB2;APBB2;APBB2 | NM\_004307;NM\_173075;NM\_001166050 | 3.698694 |
| cg08418664 | chr22 | 26056515 | ADRBK2 | NM\_005160 | 3.689437 |
| cg21123577 | chr15 | 84513792 | ADAMTSL3 | NM\_207517 | 3.559855 |
| cg25190663 | chr10 | 52687485 |  |  | 3.480379 |
| cg26434964 | chr12 | 90647590 |  |  | 3.470519 |
| cg22156534 | chr17 | 560837 | VPS53;VPS53 | NM\_018289;NM\_001128159 | 3.459970 |
| cg05941378 | chr15 | 64456365 | PPIB | NM\_000942 | 3.363764 |
| cg17302532 | chr8 | 48533808 | SPIDR;SPIDR;SPIDR;SPIDR | NR\_104581;NM\_001080394;NM\_001282916;NM\_001282919 | 3.352831 |

|  | chromosome | position | gene.symbol | gene.accession | factor.weight |
| --- | --- | --- | --- | --- | --- |
| cg12526997 | chr2 | 242974096 |  |  | 5.724378 |
| cg14270434 | chr9 | 100613647 |  |  | 5.194678 |
| cg16403049 | chr13 | 97888209 | MBNL2;MBNL2;MBNL2 | NM\_001306070;NM\_144778;NM\_207304 | 4.851909 |
| cg09997225 | chr5 | 74105810 | FAM169A;FAM169A | NR\_046462;NM\_015566 | 4.703815 |
| cg02722539 | chr2 | 68654573 |  |  | 4.628353 |
| cg16434331 | chr17 | 70712540 | SLC39A11;SLC39A11 | NM\_001159770;NM\_139177 | 4.584806 |
| cg01393939 | chr1 | 87803705 | LMO4 | NM\_006769 | 4.488239 |
| cg04255230 | chr2 | 74727010 | LBX2 | NM\_001009812 | 4.453754 |
| cg21333377 | chr5 | 40052049 | LINC00603 | NR\_104633 | 4.412990 |
| cg09746470 | chr10 | 115317373 | HABP2;HABP2 | NM\_001177660;NM\_004132 | 4.405577 |

|  | chromosome | position | gene.symbol | gene.accession | factor.weight |
| --- | --- | --- | --- | --- | --- |
| cg06251719 | chr7 | 7285084 | C1GALT1 | NM\_020156 | 2.491637 |
| cg12869334 | chr8 | 37699360 | GPR124 | NM\_032777 | 2.405931 |
| cg19452633 | chr17 | 42164476 | HDAC5;HDAC5 | NM\_001015053;NM\_005474 | 2.398783 |
| cg07001918 | chr7 | 7189008 |  |  | 2.349755 |
| cg08101036 | chr7 | 27153655 | HOXA3;HOXA3;HOXA3 | NM\_153631;NM\_030661;NM\_153632 | 2.313163 |
| cg20408776 | chr17 | 43319382 | FMNL1 | NM\_005892 | 2.277941 |
| cg21617353 | chr17 | 43319371 | FMNL1 | NM\_005892 | 2.197374 |
| cg00668519 | chr3 | 11597941 | VGLL4;ATG7;VGLL4;ATG7;VGLL4;ATG7;VGLL4 | NM\_014667;NM\_001144912;NM\_001128220;NM\_001136031;NM\_001128219;NM\_006395;NM\_001128221 | 2.169151 |
| cg00469814 | chr4 | 188953677 |  |  | 2.145529 |
| cg07277884 | chr15 | 50010605 |  |  | 2.134241 |

|  | chromosome | position | gene.symbol | gene.accession | factor.weight |
| --- | --- | --- | --- | --- | --- |
| cg00883027 | chr9 | 5455361 | CD274;CD274;CD274 | NM\_014143;NM\_001267706;NR\_052005 | 4.589917 |
| cg18220699 | chr5 | 16754962 | MYO10 | NM\_012334 | 4.564961 |
| cg03243701 | chr3 | 112348304 | CCDC80;CCDC80 | NM\_199512;NM\_199511 | 4.557195 |
| cg26739766 | chr10 | 117205016 | ATRNL1 | NM\_207303 | 4.543064 |
| cg12642480 | chr22 | 40985395 | MKL1;MKL1;MKL1 | NM\_001282662;NM\_020831;NM\_001282661 | 4.536200 |
| cg20477546 | chr4 | 126973204 |  |  | 4.533130 |
| cg02790526 | chr15 | 101470215 | LRRK1 | NM\_024652 | 4.529696 |
| cg21699506 | chr4 | 80786707 |  |  | 4.528421 |
| cg12803053 | chr8 | 126142264 | NSMCE2 | NM\_173685 | 4.519864 |
| cg02703190 | chr5 | 93137476 | FAM172A;FAM172A;FAM172A;FAM172A | NR\_028080;NM\_032042;NM\_001163418;NM\_001163417 | 4.517979 |

|  | chromosome | position | gene.symbol | gene.accession | factor.weight |
| --- | --- | --- | --- | --- | --- |
| cg13340765 | chr8 | 142276287 |  |  | 2.524403 |
| cg00697916 | chr3 | 65342644 | MAGI1;MAGI1 | NM\_001033057;NM\_015520 | 2.305225 |
| cg10838410 | chr12 | 6659524 | IFFO1;IFFO1;IFFO1 | NM\_080730;NM\_080731;NM\_001039670 | 2.269952 |
| cg23235622 | chr20 | 34039349 |  |  | 2.216436 |
| cg04865531 | chr22 | 51159147 | SHANK3 | NM\_001080420 | 2.167248 |
| cg09363375 | chr6 | 1247581 |  |  | 2.073260 |
| cg16802439 | chr16 | 88907184 | GALNS | NM\_000512 | 2.069004 |
| cg18616175 | chr20 | 39321473 |  |  | 2.045027 |
| cg00880290 | chr19 | 14550999 | PKN1;PKN1 | NM\_213560;NM\_002741 | 2.038073 |
| cg08464513 | chr16 | 30136024 | MAPK3;MAPK3;MAPK3 | NM\_001040056;NM\_001109891;NM\_002746 | 2.038061 |

##### Plot top weights per factor metabolomics

Display only the top features with highest loading

```
lapply(1:10, function(f) {
  plot_top_weights(model, factors = f, view = 2) 
})
```

```
## [[1]]
```

```
## 
## [[2]]
```

```
## 
## [[3]]
```

```
## 
## [[4]]
```

```
## 
## [[5]]
```

```
## 
## [[6]]
```

```
## 
## [[7]]
```

```
## 
## [[8]]
```

```
## 
## [[9]]
```

```
## 
## [[10]]
```

##### Correlation with behavorial data

```
library(corrplot)
```

```
## corrplot 0.92 loaded
```

```
factors <- MOFA2::get_factors(model, factors = "all")
rownames(cbcl_mca_coord)<- cbcl_mca_coord[,1]
cbcl_mca_coord<-cbcl_mca_coord[,-1]
factors <- as.data.frame(factors)

common_rows <- intersect(rownames(cbcl_mca_coord), rownames(factors))
cbcl_mca_coord_subset <- cbcl_mca_coord[rownames(cbcl_mca_coord) %in% common_rows,]
colnames(cbcl_mca_coord_subset) <- gsub(
  "Dim.", "Dim", paste0("MCA.", colnames(cbcl_mca_coord_subset)))
factors_subset <- factors[rownames(factors) %in% common_rows,]
colnames(factors_subset) <- gsub("group1", "MOFA", colnames(factors_subset))

correlation_cbcl_MOFA <- cor(
  cbcl_mca_coord_subset, factors_subset, 
  use = "pairwise.complete.obs", method = "spearman") #default: pearson
my_lim <- max(abs(c(min(correlation_cbcl_MOFA), max(correlation_cbcl_MOFA))))

# significance test - FDR-corrected p-values (Benjamini-Hochberg)
sigTest = psych::corr.test(
  cbcl_mca_coord_subset, factors_subset,
  use = "pairwise",
  method = "spearman",
  adjust = "BH")

corrplot(
  correlation_cbcl_MOFA,
  is.corr = FALSE,
  col = COL2("RdBu", 200),
  col.lim = c(-my_lim, my_lim),
  p.mat = sigTest$p.adj
)
```

##### Correlation with phenotype covariates

```
rownames(metaData_cor) <- metaData_cor[,1]
metaData_cor <- metaData_cor[,-1]

commons_rows <- intersect(rownames(metaData_cor), rownames(factors))
metaData_subset <- metaData_cor[rownames(metaData_cor) %in% common_rows,]
metaData_subset <- metaData_subset[, c('Age', 'Aggression_Tscore')]

print(dim(metaData_cor))
```

```
## [1] 189  18
```

```
print(dim(factors_subset))
```

```
## [1] 165  10
```

```
# correlations
correlation_metadata_MOFA <- cor(
  metaData_subset, factors_subset, use = "pairwise.complete.obs",
  method = "spearman") #default: pearson
my_lim <- max(abs(c(min(correlation_metadata_MOFA), 
                    max(correlation_metadata_MOFA))))

# significance test - FDR-corrected p-values (Benjamini-Hochberg)
sigTest = psych::corr.test(
  metaData_subset, factors_subset,
  use = "pairwise",
  method = "spearman",
  adjust = "BH")

# plot
corrplot(
  correlation_metadata_MOFA,
  is.corr = FALSE,
  col = COL2("RdBu", 200),
  col.lim = c(-my_lim, my_lim),
  p.mat = sigTest$p.adj
)
```

##### Boxplot Factors vs. Sex

```
# get factor values per male/female
factors_male <- get_factors(
  model, factor = 1:10)$group1[metaData[metaData$Sex == "Male", ]$sample, ]
factors_female <- get_factors(
  model, factor = 1:10)$group1[metaData[metaData$Sex == "Male", ]$sample, ]
# Wilcoxon rank sum test
wilcox_MOFA_sex <- do.call(rbind, lapply(1:10, function(x) {
  data.frame(
    factor = x,
    p.value = wilcox.test(factors_male[, x], factors_female[, x],
                          alternative = "two.sided")$p.value)
}))
wilcox_MOFA_sex$p.adj <- p.adjust(wilcox_MOFA_sex$p.value, method = "BH")
knitr::kable(wilcox_MOFA_sex)
```

| factor | p.value | p.adj |
| --- | --- | --- |
| 1 | 1 | 1 |
| 2 | 1 | 1 |
| 3 | 1 | 1 |
| 4 | 1 | 1 |
| 5 | 1 | 1 |
| 6 | 1 | 1 |
| 7 | 1 | 1 |
| 8 | 1 | 1 |
| 9 | 1 | 1 |
| 10 | 1 | 1 |

```
# box plot
plot_factor(
  model,
  factor = 1:10,
  color_by = "Sex",
  dodge = TRUE,
  add_boxplot = TRUE
) +
  facet_wrap(~factor, nrow = 2, scales = "free",
             labeller = labeller(factor = wilcox_MOFA_sex))
```

```
## Warning: `fct_explicit_na()` was deprecated in forcats 1.0.0.
## ℹ Please use `fct_na_value_to_level()` instead.
## ℹ The deprecated feature was likely used in the MOFA2 package.
##   Please report the issue at <https://github.com/bioFAM/MOFA2>.
```

##### Check potential confounding factors

##### Boxplot Factors vs. Sick

```
plot_factor(
  model,
  factor = 1:10,
  color_by = "Sick",
  dodge= TRUE,
  add_boxplot = TRUE
) +
  facet_wrap(~factor, nrow = 2, scales = "free")
```

##### Boxplot Factors vs. Menstruation

```
plot_factor(
  model,
  factor = 1:10,
  color_by = "Menstruation",
  dodge= TRUE,
  add_boxplot = TRUE
) +
  facet_wrap(~factor, nrow = 2, scales = "free")
```

##### Boxplot Factors vs. Vitamine use

```
plot_factor(
  model,
  factor = 1:10,
  color_by = "Vitamines",
  dodge= TRUE,
  add_boxplot = TRUE
) +
  facet_wrap(~factor, nrow = 2, scales = "free")
```

```
sessionInfo()
```

```
## R version 4.0.2 (2020-06-22)
## Platform: x86_64-pc-linux-gnu (64-bit)
## Running under: Debian GNU/Linux bullseye/sid
## 
## Matrix products: default
## BLAS:   /usr/lib/x86_64-linux-gnu/openblas-pthread/libblas.so.3
## LAPACK: /usr/lib/x86_64-linux-gnu/openblas-pthread/libopenblasp-r0.3.10.so
## 
## locale:
##  [1] LC_CTYPE=en_US.UTF-8       LC_NUMERIC=C              
##  [3] LC_TIME=en_US.UTF-8        LC_COLLATE=en_US.UTF-8    
##  [5] LC_MONETARY=en_US.UTF-8    LC_MESSAGES=en_US.UTF-8   
##  [7] LC_PAPER=en_US.UTF-8       LC_NAME=C                 
##  [9] LC_ADDRESS=C               LC_TELEPHONE=C            
## [11] LC_MEASUREMENT=en_US.UTF-8 LC_IDENTIFICATION=C       
## 
## attached base packages:
## [1] stats     graphics  grDevices utils     datasets  methods   base     
## 
## other attached packages:
## [1] corrplot_0.92 knitr_1.42    MOFA2_1.3.4   ggplot2_3.4.0
## 
## loaded via a namespace (and not attached):
##  [1] MatrixGenerics_1.2.1 sass_0.4.5           tidyr_1.3.0         
##  [4] jsonlite_1.8.4       bslib_0.4.2          highr_0.10          
##  [7] stats4_4.0.2         yaml_2.3.7           ggrepel_0.9.1       
## [10] pillar_1.8.1         lattice_0.20-41      glue_1.6.2          
## [13] reticulate_1.20      digest_0.6.31        RColorBrewer_1.1-3  
## [16] colorspace_2.1-0     cowplot_1.1.1        htmltools_0.5.4     
## [19] Matrix_1.2-18        plyr_1.8.6           psych_2.2.9         
## [22] pkgconfig_2.0.3      pheatmap_1.0.12      purrr_1.0.1         
## [25] scales_1.2.1         HDF5Array_1.18.1     Rtsne_0.15          
## [28] tibble_3.1.8         generics_0.1.3       farver_2.1.1        
## [31] IRanges_2.24.1       cachem_1.0.6         withr_2.5.0         
## [34] BiocGenerics_0.36.1  cli_3.6.0            mnormt_2.1.1        
## [37] magrittr_2.0.3       evaluate_0.20        GGally_2.1.2        
## [40] fansi_1.0.4          nlme_3.1-149         forcats_1.0.0       
## [43] tools_4.0.2          lifecycle_1.0.3      matrixStats_0.60.0  
## [46] basilisk.utils_1.2.2 stringr_1.5.0        Rhdf5lib_1.12.1     
## [49] S4Vectors_0.28.1     munsell_0.5.0        DelayedArray_0.16.3 
## [52] compiler_4.0.2       jquerylib_0.1.4      rlang_1.0.6         
## [55] rhdf5_2.34.0         grid_4.0.2           rhdf5filters_1.2.1  
## [58] rappdirs_0.3.3       labeling_0.4.2       rmarkdown_2.20      
## [61] basilisk_1.2.1       gtable_0.3.1         reshape_0.8.8       
## [64] reshape2_1.4.4       R6_2.5.1             dplyr_1.1.0         
## [67] fastmap_1.1.0        uwot_0.1.10          utf8_1.2.2          
## [70] filelock_1.0.2       stringi_1.7.12       parallel_4.0.2      
## [73] Rcpp_1.0.7           vctrs_0.5.2          png_0.1-7           
## [76] tidyselect_1.2.0     xfun_0.36
```

---

1. Radboud University Medical Center,↩︎
2. Radboud University Medical Center,↩︎
3. Radboud University Medical Center,↩︎
