## Additional File 3 for "A Multi-omics Data Analysis Workflow Packaged as a FAIR Digital Object": Additional File 3.html

|  | x |
| --- | --- |
| epigenomics | 24.6626003 |
| metabolomics | 0.0009651 |

### Variance explained for every factor in per view

```
knitr::kable(head(get_variance_explained(model)$r2_per_factor[[1]]))
```

|  | epigenomics | metabolomics |
| --- | --- | --- |
| Factor1 | 10.4938285 | 0.0000026 |
| Factor2 | 7.9004061 | 0.0000493 |
| Factor3 | 1.2878250 | 0.0000009 |
| Factor4 | 1.1675614 | 0.0002908 |
| Factor5 | 0.8807488 | 0.0005940 |
| Factor6 | 0.8139437 | 0.0000054 |

|  | chromosome | position | gene.symbol | gene.accession | factor.weight |
| --- | --- | --- | --- | --- | --- |
| cg11541587 | chr18 | 34533108 | KIAA1328 | NM\_020776 | 0.0380038 |
| cg00830121 | chr16 | 85904858 |  |  | 0.0375090 |
| cg08910976 | chr3 | 141639452 | ATP1B3 | NM\_001679 | 0.0370445 |
| cg16954236 | chr18 | 8638786 | RAB12 | NM\_001025300 | 0.0367357 |
| cg00433611 | chr1 | 46184922 | IPP;IPP | NM\_001145349;NM\_005897 | 0.0363010 |
| cg13777782 | chr16 | 14275130 | MKL2;MKL2 | NM\_014048;NM\_001308142 | 0.0361360 |
| cg23194094 | chr12 | 7449465 |  |  | 0.0358521 |
| cg18306106 | chr10 | 13350494 |  |  | 0.0358483 |
| cg01759026 | chr14 | 90178213 |  |  | 0.0356217 |
| cg10587895 | chr8 | 61555498 |  |  | 0.0354675 |

|  | chromosome | position | gene.symbol | gene.accession | factor.weight |
| --- | --- | --- | --- | --- | --- |
| cg05367846 | chr22 | 18507520 | MICAL3;MICAL3;MICAL3 | NM\_015241;NM\_001136004;NM\_001122731 | 0.0587785 |
| cg12543831 | chr8 | 22550284 | EGR3;EGR3;EGR3 | NM\_004430;NM\_001199881;NM\_001199880 | 0.0584619 |
| cg17202948 | chr17 | 72772736 | NAT9;TMEM104 | NM\_015654;NM\_017728 | 0.0578962 |
| cg12697442 | chr11 | 101981167 | YAP1;YAP1 | NM\_001130145;NM\_006106 | 0.0542597 |
| cg16146432 | chr3 | 167967220 | C3orf50 | NR\_021485 | 0.0518771 |
| cg05898482 | chr7 | 152373203 | XRCC2;XRCC2 | NM\_005431;NM\_005431 | 0.0502534 |
| cg26393261 | chr6 | 16467598 | ATXN1;ATXN1 | NM\_001128164;NM\_000332 | 0.0498565 |
| cg25706478 | chr22 | 45636726 | KIAA0930 | NM\_001009880 | 0.0498524 |
| cg26547526 | chr10 | 2666743 |  |  | 0.0497938 |
| cg27521571 | chr22 | 19938424 | COMT;COMT;COMT | NM\_000754;NM\_001135162;NM\_001135161 | 0.0487930 |

|  | chromosome | position | gene.symbol | gene.accession | factor.weight |
| --- | --- | --- | --- | --- | --- |
| cg08861957 | chr5 | 130596889 |  |  | 0.0360841 |
| cg06620193 | chr3 | 78803218 | ROBO1;ROBO1;ROBO1 | NM\_001145845;NM\_133631;NM\_002941 | 0.0358957 |
| cg05615397 | chr1 | 198202503 | NEK7 | NM\_133494 | 0.0354983 |
| cg26531716 | chr1 | 216897565 | ESRRG;ESRRG;ESRRG;ESRRG;ESRRG;ESRRG;ESRRG;ESRRG;ESRRG;ESRRG;ESRRG;ESRRG;ESRRG;ESRRG;ESRRG;ESRRG | NM\_001438;NM\_001243519;NM\_001243518;NM\_001243515;NM\_001243514;NM\_001243505;NM\_001243506;NM\_001243513;NM\_001243512;NM\_001243509;NM\_001243507;NM\_206595;NM\_206594;NM\_001134285;NM\_001243511;NM\_001243510 | 0.0349324 |
| cg16907986 | chr3 | 63947642 | ATXN7;ATXN7 | NM\_000333;NM\_001177387 | 0.0348529 |
| cg04346791 | chr12 | 32607254 |  |  | 0.0347745 |
| cg21449990 | chr4 | 167013271 | TLL1 | NM\_012464 | 0.0344427 |
| cg21626732 | chr13 | 75445077 |  |  | 0.0342927 |
| cg12142326 | chr4 | 74089113 | ANKRD17;ANKRD17;ANKRD17 | NM\_001286771;NM\_198889;NM\_032217 | 0.0341664 |
| cg11373565 | chr12 | 31359011 |  |  | 0.0338619 |

|  | chromosome | position | gene.symbol | gene.accession | factor.weight |
| --- | --- | --- | --- | --- | --- |
| cg04777726 | chr19 | 49340489 | PLEKHA4;PLEKHA4;HSD17B14 | NM\_001161354;NM\_020904;NM\_016246 | 0.0570381 |
| cg18121224 | chr5 | 176559563 | NSD1;NSD1 | NM\_022455;NM\_172349 | 0.0537834 |
| cg18691800 | chr6 | 17281015 | RBM24;RBM24 | NM\_001143942;NM\_001143941 | 0.0534553 |
| cg05690644 | chr8 | 97158015 | GDF6 | NM\_001001557 | 0.0533533 |
| cg16425038 | chr14 | 24641194 | REC8;REC8 | NM\_001048205;NM\_005132 | 0.0524984 |
| cg15605704 | chr1 | 4770676 | AJAP1;AJAP1 | NM\_018836;NM\_001042478 | 0.0524899 |
| cg05825244 | chr20 | 2730488 | EBF4 | NM\_001110514 | 0.0519255 |
| cg12749132 | chr1 | 151811364 | LOC100132111;C2CD4D | NR\_024237;NM\_001136003 | 0.0515498 |
| cg04015962 | chr1 | 10949192 |  |  | 0.0500643 |
| cg15384383 | chr10 | 99798415 |  |  | 0.0492006 |

|  | chromosome | position | gene.symbol | gene.accession | factor.weight |
| --- | --- | --- | --- | --- | --- |
| cg08599635 | chr16 | 78496508 | WWOX | NM\_016373 | 0.0552797 |
| cg07541744 | chr16 | 55363277 | IRX6 | NM\_024335 | 0.0515566 |
| cg19033383 | chr3 | 152217682 |  |  | 0.0498585 |
| cg05753543 | chr7 | 51189600 | COBL;COBL;COBL | NM\_015198;NM\_001287436;NM\_001287438 | 0.0495604 |
| cg02722539 | chr2 | 68654573 |  |  | 0.0484607 |
| cg14030258 | chr1 | 29591301 | PTPRU;PTPRU;PTPRU | NM\_133178;NM\_133177;NM\_005704 | 0.0484429 |
| cg12938003 | chr4 | 109092724 | LOC641518;LOC641518 | NR\_029373;NR\_029374 | 0.0479900 |
| cg16032803 | chr12 | 12009167 | ETV6;RNU6-19P | NM\_001987;NR\_046487 | 0.0476469 |
| cg22726079 | chr10 | 71880348 | AIFM2;AIFM2;AIFM2;AIFM2 | NM\_001198696;NM\_032797;NM\_001198696;NM\_032797 | 0.0468169 |
| cg02318232 | chr5 | 88599237 |  |  | 0.0468036 |

|  | chromosome | position | gene.symbol | gene.accession | factor.weight |
| --- | --- | --- | --- | --- | --- |
| cg00087792 | chr16 | 4000986 |  |  | 0.0328660 |
| cg09674340 | chr1 | 202509286 | PPP1R12B;PPP1R12B;PPP1R12B;PPP1R12B | NM\_001197131;NM\_032104;NM\_032103;NM\_002481 | 0.0323340 |
| cg08853285 | chr10 | 122942756 |  |  | 0.0316681 |
| cg12691488 | chr1 | 243053673 |  |  | 0.0315809 |
| cg04775572 | chr8 | 29356671 |  |  | 0.0305277 |
| cg12803754 | chr5 | 14676460 | FAM105B | NM\_138348 | 0.0300223 |
| cg16207263 | chr14 | 99413386 |  |  | 0.0299420 |
| cg12651173 | chr6 | 31823344 |  |  | 0.0298578 |
| cg16600370 | chr3 | 49851721 | UBA7 | NM\_003335 | 0.0297123 |
| cg18384588 | chr22 | 46463747 |  |  | 0.0295880 |

|  | chromosome | position | gene.symbol | gene.accession | factor.weight |
| --- | --- | --- | --- | --- | --- |
| cg03652396 | chr3 | 194425225 |  |  | 0.0704232 |
| cg04155451 | chr22 | 22805066 |  |  | 0.0600308 |
| cg06129556 | chr15 | 64634880 | CSNK1G1 | NM\_022048 | 0.0597010 |
| cg09072064 | chr22 | 41065457 |  |  | 0.0570018 |
| cg04093108 | chr6 | 15264508 | JARID2;JARID2 | NM\_004973;NM\_001267040 | 0.0568236 |
| cg06066640 | chr16 | 66692347 | CMTM4;CMTM4 | NM\_178818;NM\_181521 | 0.0566115 |
| cg02917016 | chr13 | 46401058 | SIAH3 | NM\_198849 | 0.0558689 |
| cg15556865 | chr11 | 94202317 | MRE11A;MRE11A | NM\_005591;NM\_005590 | 0.0552667 |
| cg24575378 | chr1 | 24583120 |  |  | 0.0546020 |
| cg03611990 | chr6 | 96980568 | UFL1 | NM\_015323 | 0.0537473 |

|  | chromosome | position | gene.symbol | gene.accession | factor.weight |
| --- | --- | --- | --- | --- | --- |
| cg19846154 | chr8 | 74332112 |  |  | 0.0834817 |
| cg07651316 | chr16 | 3641320 | BTBD12 | NM\_032444 | 0.0830885 |
| cg05080979 | chr12 | 107766282 | BTBD11 | NM\_001018072 | 0.0779408 |
| cg04720277 | chr2 | 192482590 |  |  | 0.0772774 |
| cg08738297 | chr7 | 144539618 |  |  | 0.0757809 |
| cg07532919 | chr2 | 235313794 |  |  | 0.0750867 |
| cg22867816 | chr4 | 16081205 | PROM1;PROM1 | NM\_001145848;NM\_001145847 | 0.0748724 |
| cg22375763 | chr19 | 4540003 | LRG1 | NM\_052972 | 0.0738526 |
| cg19652019 | chr2 | 101790485 |  |  | 0.0721332 |
| cg16887696 | chr20 | 50361345 | ATP9A | NM\_006045 | 0.0706330 |

|  | chromosome | position | gene.symbol | gene.accession | factor.weight |
| --- | --- | --- | --- | --- | --- |
| cg15096079 | chr17 | 5233746 | RABEP1;RABEP1;RABEP1;RABEP1 | NM\_001291582;NM\_004703;NM\_001083585;NM\_001291581 | 0.0289822 |
| cg20264491 | chr8 | 19649875 |  |  | 0.0283670 |
| cg11921583 | chr2 | 143910340 | ARHGAP15 | NM\_018460 | 0.0276012 |
| cg01097027 | chr22 | 30598622 |  |  | 0.0271762 |
| cg05579179 | chr9 | 125765159 | RABGAP1 | NM\_012197 | 0.0266792 |
| cg23368104 | chr10 | 74271297 | MICU1;MICU1;MICU1 | NM\_001195519;NM\_001195518;NM\_006077 | 0.0266787 |
| cg01761653 | chr3 | 155860011 | KCNAB1;KCNAB1;KCNAB1 | NM\_003471;NM\_001308217;NM\_172160 | 0.0259206 |
| cg22615730 | chr2 | 69997225 | ANXA4 | NM\_001153 | 0.0258751 |
| cg17493526 | chr12 | 20524048 | PDE3A | NM\_000921 | 0.0256017 |
| cg05851077 | chr9 | 29802712 |  |  | 0.0250036 |

|  | chromosome | position | gene.symbol | gene.accession | factor.weight |
| --- | --- | --- | --- | --- | --- |
| cg15077985 | chr18 | 59639286 |  |  | 0.0395679 |
| cg14388155 | chr8 | 125315228 |  |  | 0.0356046 |
| cg08742889 | chr14 | 28175496 |  |  | 0.0355356 |
| cg17097142 | chr4 | 135053718 |  |  | 0.0352952 |
| cg25337491 | chr2 | 21224422 | APOB | NM\_000384 | 0.0343363 |
| cg10232752 | chr2 | 180051001 | SESTD1 | NM\_178123 | 0.0337829 |
| cg13079633 | chr8 | 90766731 |  |  | 0.0336940 |
| cg20506071 | chr2 | 217985565 |  |  | 0.0335970 |
| cg11025439 | chr14 | 41817479 |  |  | 0.0320826 |
| cg11835442 | chr20 | 23470461 | CST8 | NM\_005492 | 0.0313926 |

### Plot top weights per factor metabolomics

Display only the top features with highest loading

```
lapply(1:10, function(f) {
  plot_top_weights(model, factors = f, view = 2) 
})
```

```
## [[1]]
```

```
## 
## [[2]]
```

```
## 
## [[3]]
```

```
## 
## [[4]]
```

```
## 
## [[5]]
```

```
## 
## [[6]]
```

```
## 
## [[7]]
```

```
## 
## [[8]]
```

```
## 
## [[9]]
```

```
## 
## [[10]]
```

### Correlation with behavorial data

```
library(corrplot)
```

```
## corrplot 0.92 loaded
```

```
factors <- MOFA2::get_factors(model, factors = "all")
rownames(cbcl_mca_coord)<- cbcl_mca_coord[,1]
cbcl_mca_coord<-cbcl_mca_coord[,-1]
factors <- as.data.frame(factors)

common_rows <- intersect(rownames(cbcl_mca_coord), rownames(factors))
cbcl_mca_coord_subset <- cbcl_mca_coord[rownames(cbcl_mca_coord) %in% common_rows,]
colnames(cbcl_mca_coord_subset) <- gsub(
  "Dim.", "Dim", paste0("MCA.", colnames(cbcl_mca_coord_subset)))
factors_subset <- factors[rownames(factors) %in% common_rows,]
colnames(factors_subset) <- gsub("group1", "MOFA", colnames(factors_subset))

correlation_cbcl_MOFA <- cor(
  cbcl_mca_coord_subset, factors_subset, 
  use = "pairwise.complete.obs", method = "spearman") #default: pearson
my_lim <- max(abs(c(min(correlation_cbcl_MOFA), max(correlation_cbcl_MOFA))))

# significance test - FDR-corrected p-values (Benjamini-Hochberg)
sigTest = psych::corr.test(
  cbcl_mca_coord_subset, factors_subset,
  use = "pairwise",
  method = "spearman",
  adjust = "BH")

corrplot(
  correlation_cbcl_MOFA,
  is.corr = FALSE,
  col = COL2("RdBu", 200),
  col.lim = c(-my_lim, my_lim),
  p.mat = sigTest$p.adj
)
```

### Correlation with phenotype covariates

```
rownames(metaData_cor) <- metaData_cor[,1]
metaData_cor <- metaData_cor[,-1]

commons_rows <- intersect(rownames(metaData_cor), rownames(factors))
metaData_subset <- metaData_cor[rownames(metaData_cor) %in% common_rows,]
metaData_subset2 <- metaData_subset[, c('Age', 'Aggression_Tscore', 'Familynumber')]
metaData_subset <- metaData_subset[, c('Age', 'Aggression_Tscore')]

# correlations
correlation_metadata_MOFA <- cor(
  metaData_subset, factors_subset, use = "pairwise.complete.obs",
  method = "spearman") #default: pearson
my_lim <- max(abs(c(min(correlation_metadata_MOFA), 
                    max(correlation_metadata_MOFA))))

# significance test - FDR-corrected p-values (Benjamini-Hochberg)
sigTest = psych::corr.test(
  metaData_subset, factors_subset,
  use = "pairwise",
  method = "spearman",
  adjust = "BH")

# plot
corrplot(
  correlation_metadata_MOFA,
  is.corr = FALSE,
  col = COL2("RdBu", 200),
  col.lim = c(-my_lim, my_lim),
  p.mat = sigTest$p.adj
)
```

### Boxplot Factors vs. Sex

```
# get factor values per male/female
factors_male <- get_factors(
  model, factor = 1:10)$group1[metaData[metaData$Sex == "Male", ]$sample, ]
factors_female <- get_factors(
  model, factor = 1:10)$group1[metaData[metaData$Sex == "Male", ]$sample, ]
# Wilcoxon rank sum test
wilcox_MOFA_sex <- do.call(rbind, lapply(1:10, function(x) {
  data.frame(
    factor = x,
    p.value = wilcox.test(factors_male[, x], factors_female[, x],
                          alternative = "two.sided")$p.value)
}))
wilcox_MOFA_sex$p.adj <- p.adjust(wilcox_MOFA_sex$p.value, method = "BH")
knitr::kable(wilcox_MOFA_sex)
```

| factor | p.value | p.adj |
| --- | --- | --- |
| 1 | 1 | 1 |
| 2 | 1 | 1 |
| 3 | 1 | 1 |
| 4 | 1 | 1 |
| 5 | 1 | 1 |
| 6 | 1 | 1 |
| 7 | 1 | 1 |
| 8 | 1 | 1 |
| 9 | 1 | 1 |
| 10 | 1 | 1 |

```
# box plot
plot_factor(
  model,
  factor = 1:10,
  color_by = "Sex",
  dodge = TRUE,
  add_boxplot = TRUE
) +
  facet_wrap(~factor, nrow = 2, scales = "free",
             labeller = labeller(factor = wilcox_MOFA_sex))
```

```
## Warning: `fct_explicit_na()` was deprecated in forcats 1.0.0.
## ℹ Please use `fct_na_value_to_level()` instead.
## ℹ The deprecated feature was likely used in the MOFA2 package.
##   Please report the issue at <https://github.com/bioFAM/MOFA2>.
## This warning is displayed once every 8 hours.
## Call `lifecycle::last_lifecycle_warnings()` to see where this warning was
## generated.
```

### Check potential confounding factors

### Boxplot Factors vs. Sick

```
plot_factor(
  model,
  factor = 1:10,
  color_by = "Sick",
  dodge= TRUE,
  add_boxplot = TRUE
) +
  facet_wrap(~factor, nrow = 2, scales = "free")
```

### Boxplot Factors vs. Menstruation

```
plot_factor(
  model,
  factor = 1:10,
  color_by = "Menstruation",
  dodge= TRUE,
  add_boxplot = TRUE
) +
  facet_wrap(~factor, nrow = 2, scales = "free")
```

### Boxplot Factors vs. Vitamine use

```
plot_factor(
  model,
  factor = 1:10,
  color_by = "Vitamines",
  dodge= TRUE,
  add_boxplot = TRUE
) +
  facet_wrap(~factor, nrow = 2, scales = "free")
```

```
sessionInfo()
```

```
## R version 4.0.2 (2020-06-22)
## Platform: x86_64-pc-linux-gnu (64-bit)
## Running under: Debian GNU/Linux bullseye/sid
## 
## Matrix products: default
## BLAS:   /usr/lib/x86_64-linux-gnu/openblas-pthread/libblas.so.3
## LAPACK: /usr/lib/x86_64-linux-gnu/openblas-pthread/libopenblasp-r0.3.10.so
## 
## locale:
##  [1] LC_CTYPE=en_US.UTF-8       LC_NUMERIC=C              
##  [3] LC_TIME=en_US.UTF-8        LC_COLLATE=en_US.UTF-8    
##  [5] LC_MONETARY=en_US.UTF-8    LC_MESSAGES=en_US.UTF-8   
##  [7] LC_PAPER=en_US.UTF-8       LC_NAME=C                 
##  [9] LC_ADDRESS=C               LC_TELEPHONE=C            
## [11] LC_MEASUREMENT=en_US.UTF-8 LC_IDENTIFICATION=C       
## 
## attached base packages:
## [1] stats     graphics  grDevices utils     datasets  methods   base     
## 
## other attached packages:
## [1] corrplot_0.92 knitr_1.42    MOFA2_1.3.4   ggplot2_3.4.2
## 
## loaded via a namespace (and not attached):
##  [1] MatrixGenerics_1.2.1 sass_0.4.5           tidyr_1.3.0         
##  [4] jsonlite_1.8.4       bslib_0.4.2          highr_0.10          
##  [7] stats4_4.0.2         yaml_2.3.7           ggrepel_0.9.1       
## [10] pillar_1.9.0         lattice_0.20-41      glue_1.6.2          
## [13] reticulate_1.20      digest_0.6.31        RColorBrewer_1.1-3  
## [16] colorspace_2.1-0     cowplot_1.1.1        htmltools_0.5.5     
## [19] Matrix_1.2-18        plyr_1.8.6           psych_2.3.3         
## [22] pkgconfig_2.0.3      pheatmap_1.0.12      purrr_1.0.1         
## [25] scales_1.2.1         HDF5Array_1.18.1     Rtsne_0.15          
## [28] tibble_3.2.1         generics_0.1.3       farver_2.1.1        
## [31] IRanges_2.24.1       cachem_1.0.7         withr_2.5.0         
## [34] BiocGenerics_0.36.1  cli_3.6.1            mnormt_2.1.1        
## [37] crayon_1.5.2         magrittr_2.0.3       evaluate_0.20       
## [40] GGally_2.1.2         fansi_1.0.4          nlme_3.1-149        
## [43] forcats_1.0.0        tools_4.0.2          lifecycle_1.0.3     
## [46] matrixStats_0.60.0   basilisk.utils_1.2.2 stringr_1.5.0       
## [49] Rhdf5lib_1.12.1      S4Vectors_0.28.1     munsell_0.5.0       
## [52] DelayedArray_0.16.3  compiler_4.0.2       jquerylib_0.1.4     
## [55] rlang_1.1.0          rhdf5_2.34.0         grid_4.0.2          
## [58] rhdf5filters_1.2.1   rappdirs_0.3.3       labeling_0.4.2      
## [61] rmarkdown_2.21       basilisk_1.2.1       gtable_0.3.3        
## [64] reshape_0.8.8        reshape2_1.4.4       R6_2.5.1            
## [67] dplyr_1.1.1          fastmap_1.1.1        uwot_0.1.10         
## [70] utf8_1.2.3           filelock_1.0.2       stringi_1.7.12      
## [73] parallel_4.0.2       Rcpp_1.0.7           vctrs_0.6.1         
## [76] png_0.1-7            tidyselect_1.2.0     xfun_0.38
```

---

1. Radboud University Medical Center,↩︎
2. Radboud University Medical Center,↩︎
3. Radboud University Medical Center,↩︎
