## Additional File 2 for "A Multi-omics Data Analysis Workflow Packaged as a FAIR Digital Object": Additional File 2.html

Multiple Correspondence analysis of CBCL behavioral data


### Multiple Correspondence analysis of CBCL behavioral data

###### Anna Niehues^1^

This notebook performs filtering based on missingness, missing-value imputation, and multiple correspondence analysis (MCA). Visualizations of data and results are plotted.

```
print(params)
```

```
## $missing_value_cutoff
## [1] 0.5
## 
## $cbcl_infile
## [1] "cbcl_data.csv"
## 
## $cbcl_labels
## [1] "cbcl_labels.csv"
## 
## $cbcl_filtered_outfile
## [1] "cbcl_filtered.csv"
## 
## $cbcl_imputed_outfile
## [1] "cbcl_imputed.csv"
## 
## $cbcl_mca_coord_outfile
## [1] "cbcl_mca_coord.csv"
## 
## $cbcl_mca_outfile
## [1] "cbcl_mca.RData"
## 
## $imputation_method
## [1] "RF"
```

Read phenotypic data (samples in rows, survey elements in columns).

```
# read phenotypic data
pheno_in <- read.csv(params$cbcl_infile, row.names = 1)
vars_in <- colnames(pheno_in)
# labels 
pheno_labels <- read.csv(params$cbcl_labels)
knitr::kable(pheno_labels)
```

| X | variable | label |
| --- | --- | --- |
| 1 | CBCL6-18\_q3 | CBCL6\_18 - 3 Argues a lot |
| 2 | CBCL6-18\_q16 | CBCL6\_18 - 16 Cruel to others |
| 3 | CBCL6-18\_q19 | CBCL6\_18 - 19 Demands a lot of attention |
| 4 | CBCL6-18\_q20 | CBCL6\_18 - 20 Destroys own things |
| 5 | CBCL6-18\_q21 | CBCL6\_18 - 21 Destroys others things |
| 6 | CBCL6-18\_q22 | CBCL6\_18 - 22 Disobedient at home |
| 7 | CBCL6-18\_q23 | CBCL6\_18 - 23 Disobedient at school |
| 8 | CBCL6-18\_q37 | CBCL6\_18 - 37 Gets in fights |
| 9 | CBCL6-18\_q57 | CBCL6\_18 - 57 Attacks others |
| 10 | CBCL6-18\_q68 | CBCL6\_18 - 68 Screams a lot |
| 11 | CBCL6-18\_q86 | CBCL6\_18 - 86 Stubborn, sullen |
| 12 | CBCL6-18\_q87 | CBCL6\_18 - 87 Mood changes |
| 13 | CBCL6-18\_q88 | CBCL6\_18 - 88 Sulks |
| 14 | CBCL6-18\_q89 | CBCL6\_18 - 89 Suspicious |
| 15 | CBCL6-18\_q94 | CBCL6\_18 - 94 Teases a lot |
| 16 | CBCL6-18\_q95 | CBCL6\_18 - 95 Temper |
| 17 | CBCL6-18\_q97 | CBCL6\_18 - 97 Threatens others |
| 18 | CBCL6-18\_q104 | CBCL6\_18 - 104 Loud |

```
cat('\n\n')
```

```
# convert 999999 to NA
pheno_in[pheno_in == "999999"] <- NA
# convert to factor
pheno_in[, vars_in] <- lapply(pheno_in[, vars_in], as.factor)
```

#### Overview of input data

```
summary(pheno_in)
```

```
##  CBCL6.18_q3 CBCL6.18_q16 CBCL6.18_q19 CBCL6.18_q20 CBCL6.18_q21 CBCL6.18_q22
##  0   :49     0   :111     0   :28      0   :122     0   :123     0   :44     
##  1   :83     1   : 58     1   :60      1   : 52     1   : 58     1   :90     
##  2   :53     2   : 15     2   :97      2   : 10     2   :  4     2   :51     
##  NA's: 4     NA's:  5     NA's: 4      NA's:  5     NA's:  4     NA's: 4     
##  CBCL6.18_q23 CBCL6.18_q37 CBCL6.18_q57 CBCL6.18_q68 CBCL6.18_q86 CBCL6.18_q87
##  0   :96      0   :138     0   :132     0   :72      0   :23      0   :45     
##  1   :61      1   : 39     1   : 42     1   :68      1   :84      1   :63     
##  2   :28      2   :  8     2   : 11     2   :45      2   :78      2   :77     
##  NA's: 4      NA's:  4     NA's:  4     NA's: 4      NA's: 4      NA's: 4     
##  CBCL6.18_q88 CBCL6.18_q89 CBCL6.18_q94 CBCL6.18_q95 CBCL6.18_q97 CBCL6.18_q104
##  0   :58      0   :110     0   :129     0   :57      0   :153     0   :75      
##  1   :80      1   : 56     1   : 50     1   :54      1   : 29     1   :65      
##  2   :47      2   : 19     2   :  6     2   :74      2   :  3     2   :45      
##  NA's: 4      NA's:  4     NA's:  4     NA's: 4      NA's:  4     NA's: 4
```

```
# distribution of missing values across samples and features
# histogram of missing values per sample
hist(rowSums(is.na(pheno_in)))
```

```
# histogram of missing values per feature
hist(colSums(is.na(pheno_in)))
```

#### Filter phenotypic data

Filter features: remove non-informative (invariant) features

```
pheno_filtfeat <- pheno_in[, sapply(pheno_in, nlevels) > 1]
message("[FILTERING] Removed ", dim(pheno_in)[2]-dim(pheno_filtfeat)[2], 
        " invariant feature(s).")
```

```
## [FILTERING] Removed 0 invariant feature(s).
```

Filter samples: remove all samples that have missing values across most features.

```
pheno_filtsamp <- pheno_filtfeat[
  rowSums(is.na(pheno_filtfeat)) < params$missing_value_cutoff*dim(pheno_filtfeat)[2],]
message("[FILTERING] Removed ", dim(pheno_filtfeat)[1]-dim(pheno_filtsamp)[1], 
        " samples with more than ", params$missing_value_cutoff*100, 
        "% missing values.")
```

```
## [FILTERING] Removed 4 samples with more than 50% missing values.
```

```
# remove invariant features
pheno_filt <- pheno_filtsamp[, sapply(pheno_filtsamp, nlevels) > 1]
message("[FILTERING] Removed ", dim(pheno_filtsamp)[2]-dim(pheno_filt)[2], 
        " invariant feature(s).")
```

```
## [FILTERING] Removed 0 invariant feature(s).
```

#### Overview of filtered data

```
message("[FILTERING] Filtered data set contains ", dim(pheno_filt)[1], 
        " samples, ", dim(pheno_filt)[2], " features, and ", 
        sum(is.na(pheno_filt)), " missing values.")
```

```
## [FILTERING] Filtered data set contains 185 samples, 18 features, and 2 missing values.
```

```
print(summary(pheno_filt))
```

```
##  CBCL6.18_q3 CBCL6.18_q16 CBCL6.18_q19 CBCL6.18_q20 CBCL6.18_q21 CBCL6.18_q22
##  0:49        0   :111     0:28         0   :122     0:123        0:44        
##  1:83        1   : 58     1:60         1   : 52     1: 58        1:90        
##  2:53        2   : 15     2:97         2   : 10     2:  4        2:51        
##              NA's:  1                  NA's:  1                              
##  CBCL6.18_q23 CBCL6.18_q37 CBCL6.18_q57 CBCL6.18_q68 CBCL6.18_q86 CBCL6.18_q87
##  0:96         0:138        0:132        0:72         0:23         0:45        
##  1:61         1: 39        1: 42        1:68         1:84         1:63        
##  2:28         2:  8        2: 11        2:45         2:78         2:77        
##                                                                               
##  CBCL6.18_q88 CBCL6.18_q89 CBCL6.18_q94 CBCL6.18_q95 CBCL6.18_q97 CBCL6.18_q104
##  0:58         0:110        0:129        0:57         0:153        0:75         
##  1:80         1: 56        1: 50        1:54         1: 29        1:65         
##  2:47         2: 19        2:  6        2:74         2:  3        2:45         
##
```

```
# histogram of missing values per sample
hist(rowSums(is.na(pheno_filt)))
```

```
# histogram of missing values per feature
hist(colSums(is.na(pheno_filt)))
```

```
# write filtered data to output file
write.csv(pheno_filt, params$cbcl_filtered_outfile)
```

#### Impute missing values

Random forests imputation

```
#' Non-parametric missing value imputation in survey data using random forests
#' 
#' @param df Data frame containing subjects in rows and questions in columns 
impute_cbcl_rf <- function(df) {
  library(missForest)
  stopifnot(all(sapply(df, is.factor)))
  # hyperparameters: ntree, mtry, replace, sampsize, nodesize, maxnodes
  # suggestion: could also make use of additional phenotype information 
  # impute values
  df_imp <- missForest(
    df,
    ntree = 1000,
    variablewise = TRUE)
  return(df_imp)
}
```

Imputation using MCA

```
#' Imputing missing categorical data using Multiple Correspondence Analysis
#'  
#' @param df Data frame containing subjects in rows and questions in columns
#' @return list of different plots of MCA
impute_cbcl_mca <- function(df){
  library(missMDA)
  stopifnot(all(sapply(df, is.factor)))
  # estimate the number of dimensions for MCA by cross-validation
  ncp_est <- estim_ncpMCA(
    df, 
    ncp.min = 0, 
    ncp.max = 7,
    method = "Regularized", 
    method.cv = "Kfold",
    verbose = TRUE) 
  print(ncp_est)
  # impute missing values using estimated number of dimensions
  df_imputed <- imputeMCA(df, ncp = ncp_est$ncp)
  return(df_imputed)
}
```

Perform imputation based using chosen method

```
if (params$imputation_method == "RF") {
  pheno_rfimp_res <- impute_cbcl_rf(pheno_filt)
  names(pheno_rfimp_res)
  # out-of-bag (OOB) imputation error estimate
  pheno_rfimp_res$OOBerror
  # extract imputed (complete) data
  pheno_imp <- pheno_rfimp_res$ximp
} else if (params$imputation_method == "MCA") {
  pheno_mcaimp_res <- impute_cbcl_mca(pheno_filt)
  names(pheno_mcaimp_res)
  # extract imputed (complete) data
  pheno_imp <- pheno_mcaimp_res$completeObs
  summary(pheno_imp)
}
```

```
## Loading required package: randomForest
```

```
## randomForest 4.6-14
```

```
## Type rfNews() to see new features/changes/bug fixes.
```

```
## Loading required package: foreach
```

```
## Loading required package: itertools
```

```
## Loading required package: iterators
```

```
##   missForest iteration 1 in progress...done!
##   missForest iteration 2 in progress...done!
##   missForest iteration 3 in progress...done!
```

```
summary(pheno_imp)
```

```
##  CBCL6.18_q3 CBCL6.18_q16 CBCL6.18_q19 CBCL6.18_q20 CBCL6.18_q21 CBCL6.18_q22
##  0:49        0:111        0:28         0:123        0:123        0:44        
##  1:83        1: 59        1:60         1: 52        1: 58        1:90        
##  2:53        2: 15        2:97         2: 10        2:  4        2:51        
##  CBCL6.18_q23 CBCL6.18_q37 CBCL6.18_q57 CBCL6.18_q68 CBCL6.18_q86 CBCL6.18_q87
##  0:96         0:138        0:132        0:72         0:23         0:45        
##  1:61         1: 39        1: 42        1:68         1:84         1:63        
##  2:28         2:  8        2: 11        2:45         2:78         2:77        
##  CBCL6.18_q88 CBCL6.18_q89 CBCL6.18_q94 CBCL6.18_q95 CBCL6.18_q97 CBCL6.18_q104
##  0:58         0:110        0:129        0:57         0:153        0:75         
##  1:80         1: 56        1: 50        1:54         1: 29        1:65         
##  2:47         2: 19        2:  6        2:74         2:  3        2:45
```

```
write.csv(pheno_imp, params$cbcl_imputed_outfile)
```

```
# prepare data for plot
library(reshape2)
library(stringr)
tmp_pheno_list <- list(filtered = pheno_filt, imputed = pheno_imp)
tmp_pheno_list <- lapply(names(tmp_pheno_list), function(x) {
  df <- tmp_pheno_list[[x]]
  df$data <- x
  df
})
pheno_all_wide <- do.call(rbind, tmp_pheno_list)
pheno_all_long <- melt(pheno_all_wide, id.vars = c("data"))
pheno_labels$variable_names <- make.names(pheno_labels$variable)
pheno_all_long$label <- apply(pheno_all_long, 1, function(x) {
  label <- pheno_labels[
    pheno_labels$variable_names == x[["variable"]], ][["label"]]
  str_wrap(gsub("CBCL6_18 - ", "", label), width = 12)
})

rm(tmp_pheno_list)
rm(pheno_all_wide)

# plot distribution of values, before and after imputation
library(ggplot2)
```

```
## 
## Attaching package: 'ggplot2'
```

```
## The following object is masked from 'package:randomForest':
## 
##     margin
```

```
cbf_palette <- c("#000000", "#E69F00", "#56B4E9", "#009E73", "#F0E442", "#0072B2", "#D55E00", "#CC79A7")
lapply(split(names(pheno_filt), ceiling(seq_along(names(pheno_filt)) / 4)), 
       function(x) {
         ggplot(pheno_all_long[pheno_all_long$variable %in% x,]) +
           geom_bar(aes(x = value, fill = value)) +
           scale_fill_manual(values = cbf_palette) +
           facet_grid(label ~ data, scales = "free_y")
})
```

```
## $`1`
```

```
## 
## $`2`
```

```
## 
## $`3`
```

```
## 
## $`4`
```

```
## 
## $`5`
```

#### Perform multiple correspondence analysis

Multiple correspondence analysis (MCA) is a dimension reduction technique that can be used for categorical data sets. The results are numerical variables (factors) representing the data.

```
#' Multiple Correspondence Analysis of survey question data with missing values
#'  
#' @param df Data frame containing subjects in rows and questions in columns
#' @return list of different plots of MCA
perform_mca <- function(df){
  library(FactoMineR)
  stopifnot(all(sapply(df, is.factor)))
  mca <- MCA(
    df, 
    ncp = 10, 
    graph = FALSE,
    method = "Indicator" # "Burt" 
  )
  return(mca)
}

names(pheno_imp) <- lapply(names(pheno_imp), function(x) {
  label <- pheno_labels[
    pheno_labels$variable_names == x, ][["label"]]
  str_wrap(gsub("CBCL6_18 - ", "", label), width = 16)
})

mca_pheno <- perform_mca(pheno_imp)
```

##### Quality of dimension reduction

Scree plot - percentage of variation (inertia) explained by dimensions.

```
library(factoextra)
```

```
## Welcome! Want to learn more? See two factoextra-related books at https://goo.gl/ve3WBa
```

```
knitr::kable(
  head(get_eigenvalue(mca_pheno), n = 10),
  caption = "Eigenvalues and explained variance for MCA dimensions 1-10.")
```

Eigenvalues and explained variance for MCA dimensions 1-10.

|  | eigenvalue | variance.percent | cumulative.variance.percent |
| --- | --- | --- | --- |
| Dim.1 | 0.4131722 | 20.658610 | 20.65861 |
| Dim.2 | 0.1871495 | 9.357473 | 30.01608 |
| Dim.3 | 0.1204884 | 6.024421 | 36.04050 |
| Dim.4 | 0.0916004 | 4.580021 | 40.62052 |
| Dim.5 | 0.0846851 | 4.234257 | 44.85478 |
| Dim.6 | 0.0785226 | 3.926129 | 48.78091 |
| Dim.7 | 0.0728193 | 3.640966 | 52.42188 |
| Dim.8 | 0.0672527 | 3.362633 | 55.78451 |
| Dim.9 | 0.0648551 | 3.242753 | 59.02726 |
| Dim.10 | 0.0612564 | 3.062820 | 62.09008 |

```
cat("\n\n")
```

```
fviz_screeplot(mca_pheno, addlabels = TRUE) +
  geom_point()
```

##### Relative contribution of individuals and variables

Biplot of individuals and behavior variable categories

```
fviz_mca_biplot(
  mca_pheno,
  repel = TRUE,
  label = "var",
  ggtheme = theme_minimal())
```

```
## Warning: ggrepel: 43 unlabeled data points (too many overlaps). Consider
## increasing max.overlaps
```

###### Correlation between **variables** and MCA dimensions

```
fviz_mca_var(
  mca_pheno, 
  choice = "mca.cor", 
  axes = c(1, 2),
  repel = TRUE, 
  ggtheme = theme_minimal())
```

###### Coordinates of **variable categories** and contribution to dimensions

```
fviz_mca_var(
  mca_pheno, 
  col.var = "contrib", # color by contribution to dimension
  repel = TRUE, 
  gradient.cols = c("#E69F00", "#56B4E9"),
  ggtheme = theme_minimal())
```

```
## Warning: ggrepel: 44 unlabeled data points (too many overlaps). Consider
## increasing max.overlaps
```

```
lapply(1:10, function(dim) {
  # contribution to MCA dimensions
  fviz_contrib(mca_pheno, choice = "var", axes = dim, top = 10)
})
```

```
## [[1]]
```

```
## 
## [[2]]
```

```
## 
## [[3]]
```

```
## 
## [[4]]
```

```
## 
## [[5]]
```

```
## 
## [[6]]
```

```
## 
## [[7]]
```

```
## 
## [[8]]
```

```
## 
## [[9]]
```

```
## 
## [[10]]
```

###### Degree of association of **variable categories** with dimensions

How well is a variable category represented by the dimensions?

```
fviz_mca_var(
  mca_pheno, 
  col.var = "cos2", # color by quality of representation
  repel = TRUE, 
  gradient.cols = c("#E69F00", "#56B4E9"),
  ggtheme = theme_minimal())
```

```
## Warning: ggrepel: 44 unlabeled data points (too many overlaps). Consider
## increasing max.overlaps
```

```
lapply(1:10, function(dim) {
  # quality of representation (cos2), squared cosine
  fviz_cos2(mca_pheno, choice = "var", axes = dim, top = 10)
})
```

```
## [[1]]
```

```
## 
## [[2]]
```

```
## 
## [[3]]
```

```
## 
## [[4]]
```

```
## 
## [[5]]
```

```
## 
## [[6]]
```

```
## 
## [[7]]
```

```
## 
## [[8]]
```

```
## 
## [[9]]
```

```
## 
## [[10]]
```

MCA dimension description

```
dimdesc_mca <- dimdesc(mca_pheno, axes = 1:10)
dimdesc_mca[paste0("Dim ", 1:10)]
```

```
## $`Dim 1`
## $quali
##                                       R2      p.value
## 3 Argues a lot                 0.5905476 5.134647e-36
## 95 Temper                      0.5506994 2.403620e-32
## 68 Screams a lot               0.5439541 9.327803e-32
## 22 Disobedient\nat home        0.5373319 3.463637e-31
## 97 Threatens\nothers           0.4345269 2.947726e-23
## 104 Loud                       0.4310561 5.144218e-23
## 86 Stubborn,\nsullen           0.4305103 5.613281e-23
## 16 Cruel to\nothers            0.4264527 1.071021e-22
## 87 Mood changes                0.4259191 1.165611e-22
## 21 Destroys\nothers things     0.4060125 2.592601e-21
## 94 Teases a lot                0.3889053 3.434585e-20
## 57 Attacks\nothers             0.3768012 2.046550e-19
## 88 Sulks                       0.3726917 3.721965e-19
## 19 Demands a lot\nof attention 0.3519597 7.174358e-18
## 23 Disobedient\nat school      0.3415476 3.059978e-17
## 37 Gets in\nfights             0.3411994 3.210787e-17
## 20 Destroys own\nthings        0.2449077 7.906871e-12
## 89 Suspicious                  0.2420763 1.111523e-11
## 
## $category
##                                                                    Estimate
## 3 Argues a lot=3 Argues a lot_2                                  0.69860679
## 95 Temper=95 Temper_2                                            0.59407624
## 22 Disobedient\nat home=22 Disobedient\nat home_2                0.70290863
## 87 Mood changes=87 Mood changes_2                                0.55090039
## 86 Stubborn,\nsullen=86 Stubborn,\nsullen_2                      0.62399241
## 97 Threatens\nothers=97 Threatens\nothers_1                      0.20072775
## 68 Screams a lot=68 Screams a lot_2                              0.56310080
## 19 Demands a lot\nof attention=19 Demands a lot\nof attention_2  0.52755600
## 88 Sulks=88 Sulks_2                                              0.56453130
## 104 Loud=104 Loud_2                                              0.50063558
## 23 Disobedient\nat school=23 Disobedient\nat school_2            0.57471217
## 16 Cruel to\nothers=16 Cruel to\nothers_2                        0.70000045
## 57 Attacks\nothers=57 Attacks\nothers_2                          0.63168194
## 37 Gets in\nfights=37 Gets in\nfights_2                          0.72800226
## 21 Destroys\nothers things=21 Destroys\nothers things_2          1.10266227
## 94 Teases a lot=94 Teases a lot_2                                0.85327155
## 57 Attacks\nothers=57 Attacks\nothers_1                          0.03427803
## 89 Suspicious=89 Suspicious_2                                    0.42969240
## 20 Destroys own\nthings=20 Destroys own\nthings_1                0.02637329
## 20 Destroys own\nthings=20 Destroys own\nthings_2                0.50760849
## 97 Threatens\nothers=97 Threatens\nothers_2                      0.66532342
## 89 Suspicious=89 Suspicious_1                                    0.03810237
## 68 Screams a lot=68 Screams a lot_1                              0.06408559
## 104 Loud=104 Loud_1                                              0.05079213
## 87 Mood changes=87 Mood changes_1                               -0.07258802
## 22 Disobedient\nat home=22 Disobedient\nat home_1               -0.13305310
## 19 Demands a lot\nof attention=19 Demands a lot\nof attention_1 -0.06058525
## 86 Stubborn,\nsullen=86 Stubborn,\nsullen_1                     -0.01010974
## 16 Cruel to\nothers=16 Cruel to\nothers_1                       -0.04264454
## 37 Gets in\nfights=37 Gets in\nfights_1                         -0.03005814
## 19 Demands a lot\nof attention=19 Demands a lot\nof attention_0 -0.46697075
## 94 Teases a lot=94 Teases a lot_1                               -0.07547334
## 21 Destroys\nothers things=21 Destroys\nothers things_1         -0.20722022
## 88 Sulks=88 Sulks_0                                             -0.47830845
## 89 Suspicious=89 Suspicious_0                                   -0.46779477
## 86 Stubborn,\nsullen=86 Stubborn,\nsullen_0                     -0.61388267
## 20 Destroys own\nthings=20 Destroys own\nthings_0               -0.53398178
## 23 Disobedient\nat school=23 Disobedient\nat school_0           -0.49594133
## 22 Disobedient\nat home=22 Disobedient\nat home_0               -0.56985553
## 87 Mood changes=87 Mood changes_0                               -0.47831238
## 37 Gets in\nfights=37 Gets in\nfights_0                         -0.69794412
## 21 Destroys\nothers things=21 Destroys\nothers things_0         -0.89544205
## 94 Teases a lot=94 Teases a lot_0                               -0.77779821
## 3 Argues a lot=3 Argues a lot_0                                 -0.61840933
## 57 Attacks\nothers=57 Attacks\nothers_0                         -0.66595996
## 16 Cruel to\nothers=16 Cruel to\nothers_0                       -0.65735591
## 104 Loud=104 Loud_0                                             -0.55142771
## 95 Temper=95 Temper_0                                           -0.53581275
## 97 Threatens\nothers=97 Threatens\nothers_0                     -0.86605117
## 68 Screams a lot=68 Screams a lot_0                             -0.62718639
##                                                                      p.value
## 3 Argues a lot=3 Argues a lot_2                                 2.607350e-27
## 95 Temper=95 Temper_2                                           7.174851e-27
## 22 Disobedient\nat home=22 Disobedient\nat home_2               1.531646e-26
## 87 Mood changes=87 Mood changes_2                               4.578420e-20
## 86 Stubborn,\nsullen=86 Stubborn,\nsullen_2                     1.659009e-18
## 97 Threatens\nothers=97 Threatens\nothers_1                     1.692437e-18
## 68 Screams a lot=68 Screams a lot_2                             2.351771e-17
## 19 Demands a lot\nof attention=19 Demands a lot\nof attention_2 1.708237e-16
## 88 Sulks=88 Sulks_2                                             3.634452e-16
## 104 Loud=104 Loud_2                                             5.907909e-14
## 23 Disobedient\nat school=23 Disobedient\nat school_2           1.881351e-13
## 16 Cruel to\nothers=16 Cruel to\nothers_2                       2.392836e-12
## 57 Attacks\nothers=57 Attacks\nothers_2                         4.181363e-09
## 37 Gets in\nfights=37 Gets in\nfights_2                         1.110393e-08
## 21 Destroys\nothers things=21 Destroys\nothers things_2         1.385609e-08
## 94 Teases a lot=94 Teases a lot_2                               2.534318e-08
## 57 Attacks\nothers=57 Attacks\nothers_1                         3.642214e-08
## 89 Suspicious=89 Suspicious_2                                   1.724170e-06
## 20 Destroys own\nthings=20 Destroys own\nthings_1               2.702638e-06
## 20 Destroys own\nthings=20 Destroys own\nthings_2               2.019300e-05
## 97 Threatens\nothers=97 Threatens\nothers_2                     2.321301e-04
## 89 Suspicious=89 Suspicious_1                                   2.374491e-04
## 68 Screams a lot=68 Screams a lot_1                             1.711531e-02
## 104 Loud=104 Loud_1                                             3.604367e-02
## 87 Mood changes=87 Mood changes_1                               1.433487e-02
## 22 Disobedient\nat home=22 Disobedient\nat home_1               8.966192e-03
## 19 Demands a lot\nof attention=19 Demands a lot\nof attention_1 2.513708e-04
## 86 Stubborn,\nsullen=86 Stubborn,\nsullen_1                     1.719473e-04
## 16 Cruel to\nothers=16 Cruel to\nothers_1                       4.977719e-06
## 37 Gets in\nfights=37 Gets in\nfights_1                         1.541384e-07
## 19 Demands a lot\nof attention=19 Demands a lot\nof attention_0 1.090102e-09
## 94 Teases a lot=94 Teases a lot_1                               6.024112e-10
## 21 Destroys\nothers things=21 Destroys\nothers things_1         1.140556e-10
## 88 Sulks=88 Sulks_0                                             6.337103e-11
## 89 Suspicious=89 Suspicious_0                                   3.528796e-11
## 86 Stubborn,\nsullen=86 Stubborn,\nsullen_0                     2.156716e-11
## 20 Destroys own\nthings=20 Destroys own\nthings_0               1.752745e-11
## 23 Disobedient\nat school=23 Disobedient\nat school_0           2.935097e-12
## 22 Disobedient\nat home=22 Disobedient\nat home_0               1.522935e-12
## 87 Mood changes=87 Mood changes_0                               4.453606e-13
## 37 Gets in\nfights=37 Gets in\nfights_0                         2.239534e-15
## 21 Destroys\nothers things=21 Destroys\nothers things_0         3.697573e-17
## 94 Teases a lot=94 Teases a lot_0                               1.530339e-17
## 3 Argues a lot=3 Argues a lot_0                                 7.223277e-18
## 57 Attacks\nothers=57 Attacks\nothers_0                         5.305387e-18
## 16 Cruel to\nothers=16 Cruel to\nothers_0                       3.009103e-18
## 104 Loud=104 Loud_0                                             1.629364e-19
## 95 Temper=95 Temper_0                                           1.515493e-20
## 97 Threatens\nothers=97 Threatens\nothers_0                     6.815112e-24
## 68 Screams a lot=68 Screams a lot_0                             5.817469e-26
## 
## attr(,"class")
## [1] "condes" "list"  
## 
## $`Dim 2`
## $quali
##                                        R2      p.value
## 16 Cruel to\nothers            0.35153174 7.618572e-18
## 94 Teases a lot                0.34950954 1.011404e-17
## 21 Destroys\nothers things     0.32286729 3.902251e-16
## 57 Attacks\nothers             0.31750408 8.000281e-16
## 37 Gets in\nfights             0.27564090 1.802215e-13
## 20 Destroys own\nthings        0.24295274 1.000443e-11
## 3 Argues a lot                 0.18741033 6.284752e-09
## 22 Disobedient\nat home        0.17438700 2.670937e-08
## 23 Disobedient\nat school      0.16939099 4.624810e-08
## 88 Sulks                       0.14442285 6.848336e-07
## 95 Temper                      0.13498578 1.858299e-06
## 68 Screams a lot               0.12908909 3.448337e-06
## 97 Threatens\nothers           0.12130935 7.745983e-06
## 86 Stubborn,\nsullen           0.11918990 9.644679e-06
## 87 Mood changes                0.10548549 3.930541e-05
## 104 Loud                       0.09427429 1.220954e-04
## 19 Demands a lot\nof attention 0.07916555 5.502014e-04
## 89 Suspicious                  0.04957333 9.785744e-03
## 
## $category
##                                                                    Estimate
## 94 Teases a lot=94 Teases a lot_2                                0.90607811
## 21 Destroys\nothers things=21 Destroys\nothers things_2          1.06522817
## 57 Attacks\nothers=57 Attacks\nothers_2                          0.67429617
## 37 Gets in\nfights=37 Gets in\nfights_2                          0.73313479
## 16 Cruel to\nothers=16 Cruel to\nothers_2                        0.54839553
## 20 Destroys own\nthings=20 Destroys own\nthings_2                0.60357681
## 22 Disobedient\nat home=22 Disobedient\nat home_0                0.21604241
## 86 Stubborn,\nsullen=86 Stubborn,\nsullen_0                      0.29011254
## 3 Argues a lot=3 Argues a lot_0                                  0.19256781
## 97 Threatens\nothers=97 Threatens\nothers_2                      0.70353986
## 95 Temper=95 Temper_0                                            0.19738327
## 87 Mood changes=87 Mood changes_0                                0.19686947
## 19 Demands a lot\nof attention=19 Demands a lot\nof attention_0  0.22709055
## 68 Screams a lot=68 Screams a lot_0                              0.13597505
## 23 Disobedient\nat school=23 Disobedient\nat school_2            0.22596804
## 88 Sulks=88 Sulks_0                                              0.12460136
## 104 Loud=104 Loud_0                                              0.12689202
## 89 Suspicious=89 Suspicious_0                                    0.12980472
## 23 Disobedient\nat school=23 Disobedient\nat school_0            0.04382986
## 88 Sulks=88 Sulks_2                                              0.09510282
## 89 Suspicious=89 Suspicious_1                                   -0.06783898
## 86 Stubborn,\nsullen=86 Stubborn,\nsullen_1                     -0.18705107
## 97 Threatens\nothers=97 Threatens\nothers_0                     -0.42877524
## 37 Gets in\nfights=37 Gets in\nfights_1                         -0.46580855
## 87 Mood changes=87 Mood changes_1                               -0.17613313
## 20 Destroys own\nthings=20 Destroys own\nthings_1               -0.39605589
## 21 Destroys\nothers things=21 Destroys\nothers things_1         -0.63459669
## 57 Attacks\nothers=57 Attacks\nothers_1                         -0.44847527
## 104 Loud=104 Loud_1                                             -0.17359764
## 94 Teases a lot=94 Teases a lot_1                               -0.57728709
## 95 Temper=95 Temper_1                                           -0.21257254
## 68 Screams a lot=68 Screams a lot_1                             -0.20711272
## 88 Sulks=88 Sulks_1                                             -0.21970417
## 22 Disobedient\nat home=22 Disobedient\nat home_1               -0.22488596
## 23 Disobedient\nat school=23 Disobedient\nat school_1           -0.26979791
## 3 Argues a lot=3 Argues a lot_1                                 -0.24269596
## 16 Cruel to\nothers=16 Cruel to\nothers_1                       -0.43363659
##                                                                      p.value
## 94 Teases a lot=94 Teases a lot_2                               4.923975e-15
## 21 Destroys\nothers things=21 Destroys\nothers things_2         1.726495e-14
## 57 Attacks\nothers=57 Attacks\nothers_2                         2.743608e-14
## 37 Gets in\nfights=37 Gets in\nfights_2                         1.293773e-12
## 16 Cruel to\nothers=16 Cruel to\nothers_2                       1.789311e-12
## 20 Destroys own\nthings=20 Destroys own\nthings_2               9.273138e-11
## 22 Disobedient\nat home=22 Disobedient\nat home_0               9.844146e-07
## 86 Stubborn,\nsullen=86 Stubborn,\nsullen_0                     3.607449e-06
## 3 Argues a lot=3 Argues a lot_0                                 5.326957e-06
## 97 Threatens\nothers=97 Threatens\nothers_2                     7.101797e-06
## 95 Temper=95 Temper_0                                           4.037724e-05
## 87 Mood changes=87 Mood changes_0                               8.341859e-05
## 19 Demands a lot\nof attention=19 Demands a lot\nof attention_0 1.059751e-04
## 68 Screams a lot=68 Screams a lot_0                             3.187521e-04
## 23 Disobedient\nat school=23 Disobedient\nat school_2           5.439903e-04
## 88 Sulks=88 Sulks_0                                             8.048630e-04
## 104 Loud=104 Loud_0                                             1.065099e-03
## 89 Suspicious=89 Suspicious_0                                   2.321115e-03
## 23 Disobedient\nat school=23 Disobedient\nat school_0           1.311529e-02
## 88 Sulks=88 Sulks_2                                             1.982820e-02
## 89 Suspicious=89 Suspicious_1                                   1.427169e-02
## 86 Stubborn,\nsullen=86 Stubborn,\nsullen_1                     6.422050e-03
## 97 Threatens\nothers=97 Threatens\nothers_0                     3.317806e-03
## 37 Gets in\nfights=37 Gets in\nfights_1                         1.060841e-03
## 87 Mood changes=87 Mood changes_1                               3.885390e-04
## 20 Destroys own\nthings=20 Destroys own\nthings_1               3.655156e-04
## 21 Destroys\nothers things=21 Destroys\nothers things_1         2.104835e-04
## 57 Attacks\nothers=57 Attacks\nothers_1                         9.690362e-05
## 104 Loud=104 Loud_1                                             3.695060e-05
## 94 Teases a lot=94 Teases a lot_1                               1.595223e-05
## 95 Temper=95 Temper_1                                           7.484310e-06
## 68 Screams a lot=68 Screams a lot_1                             7.346805e-07
## 88 Sulks=88 Sulks_1                                             1.023960e-07
## 22 Disobedient\nat home=22 Disobedient\nat home_1               8.846720e-08
## 23 Disobedient\nat school=23 Disobedient\nat school_1           5.995311e-08
## 3 Argues a lot=3 Argues a lot_1                                 4.143478e-09
## 16 Cruel to\nothers=16 Cruel to\nothers_1                       1.156625e-09
## 
## attr(,"class")
## [1] "condes" "list"  
## 
## $`Dim 3`
## $quali
##                                        R2      p.value
## 20 Destroys own\nthings        0.25460028 2.440168e-12
## 87 Mood changes                0.19732366 2.056752e-09
## 68 Screams a lot               0.18844443 5.597084e-09
## 19 Demands a lot\nof attention 0.16876109 4.955115e-08
## 89 Suspicious                  0.16401986 8.314428e-08
## 86 Stubborn,\nsullen           0.16005445 1.278949e-07
## 88 Sulks                       0.13558436 1.744850e-06
## 94 Teases a lot                0.13076331 2.894426e-06
## 104 Loud                       0.12498443 5.289761e-06
## 3 Argues a lot                 0.11327459 1.773426e-05
## 95 Temper                      0.10778631 3.109342e-05
## 21 Destroys\nothers things     0.07489562 8.382183e-04
## 57 Attacks\nothers             0.07372532 9.404183e-04
## 22 Disobedient\nat home        0.07242285 1.068695e-03
## 97 Threatens\nothers           0.06768039 1.699803e-03
## 37 Gets in\nfights             0.06484318 2.241219e-03
## 16 Cruel to\nothers            0.05497945 5.823095e-03
## 
## $category
##                                                                     Estimate
## 87 Mood changes=87 Mood changes_1                                0.218735592
## 20 Destroys own\nthings=20 Destroys own\nthings_2                0.451673987
## 68 Screams a lot=68 Screams a lot_1                              0.198603105
## 19 Demands a lot\nof attention=19 Demands a lot\nof attention_1  0.216862172
## 88 Sulks=88 Sulks_1                                              0.170016315
## 86 Stubborn,\nsullen=86 Stubborn,\nsullen_1                      0.202136200
## 94 Teases a lot=94 Teases a lot_2                                0.457560960
## 3 Argues a lot=3 Argues a lot_1                                  0.156361192
## 104 Loud=104 Loud_1                                              0.170674921
## 95 Temper=95 Temper_1                                            0.167527153
## 22 Disobedient\nat home=22 Disobedient\nat home_1                0.124039311
## 57 Attacks\nothers=57 Attacks\nothers_2                          0.268492936
## 89 Suspicious=89 Suspicious_1                                    0.218103021
## 37 Gets in\nfights=37 Gets in\nfights_2                          0.291364104
## 16 Cruel to\nothers=16 Cruel to\nothers_2                        0.193435485
## 21 Destroys\nothers things=21 Destroys\nothers things_2          0.310394559
## 86 Stubborn,\nsullen=86 Stubborn,\nsullen_2                      0.006890935
## 95 Temper=95 Temper_2                                           -0.074008999
## 88 Sulks=88 Sulks_2                                             -0.068383971
## 21 Destroys\nothers things=21 Destroys\nothers things_0         -0.078889514
## 95 Temper=95 Temper_0                                           -0.093518153
## 94 Teases a lot=94 Teases a lot_1                               -0.278114263
## 19 Demands a lot\nof attention=19 Demands a lot\nof attention_2 -0.027479056
## 3 Argues a lot=3 Argues a lot_0                                 -0.072758094
## 22 Disobedient\nat home=22 Disobedient\nat home_2               -0.069365581
## 87 Mood changes=87 Mood changes_2                               -0.061164267
## 97 Threatens\nothers=97 Threatens\nothers_0                     -0.038020190
## 3 Argues a lot=3 Argues a lot_2                                 -0.083603098
## 21 Destroys\nothers things=21 Destroys\nothers things_1         -0.231505045
## 97 Threatens\nothers=97 Threatens\nothers_1                     -0.252411418
## 104 Loud=104 Loud_2                                             -0.136071844
## 88 Sulks=88 Sulks_0                                             -0.101632343
## 19 Demands a lot\nof attention=19 Demands a lot\nof attention_0 -0.189383116
## 87 Mood changes=87 Mood changes_0                               -0.157571325
## 86 Stubborn,\nsullen=86 Stubborn,\nsullen_0                     -0.209027135
## 68 Screams a lot=68 Screams a lot_2                             -0.189320525
## 20 Destroys own\nthings=20 Destroys own\nthings_1               -0.340036996
## 89 Suspicious=89 Suspicious_2                                   -0.288366814
##                                                                      p.value
## 87 Mood changes=87 Mood changes_1                               9.411550e-10
## 20 Destroys own\nthings=20 Destroys own\nthings_2               6.104063e-09
## 68 Screams a lot=68 Screams a lot_1                             6.297686e-08
## 19 Demands a lot\nof attention=19 Demands a lot\nof attention_1 1.089176e-07
## 88 Sulks=88 Sulks_1                                             2.892969e-07
## 86 Stubborn,\nsullen=86 Stubborn,\nsullen_1                     9.971205e-07
## 94 Teases a lot=94 Teases a lot_2                               2.330123e-06
## 3 Argues a lot=3 Argues a lot_1                                 2.852695e-06
## 104 Loud=104 Loud_1                                             3.225813e-06
## 95 Temper=95 Temper_1                                           5.365309e-06
## 22 Disobedient\nat home=22 Disobedient\nat home_1               2.172695e-04
## 57 Attacks\nothers=57 Attacks\nothers_2                         2.309339e-04
## 89 Suspicious=89 Suspicious_1                                   2.584586e-04
## 37 Gets in\nfights=37 Gets in\nfights_2                         4.768605e-04
## 16 Cruel to\nothers=16 Cruel to\nothers_2                       2.502730e-03
## 21 Destroys\nothers things=21 Destroys\nothers things_2         1.236043e-02
## 86 Stubborn,\nsullen=86 Stubborn,\nsullen_2                     3.884431e-02
## 95 Temper=95 Temper_2                                           3.927471e-02
## 88 Sulks=88 Sulks_2                                             3.420089e-02
## 21 Destroys\nothers things=21 Destroys\nothers things_0         2.959431e-02
## 95 Temper=95 Temper_0                                           2.808960e-02
## 94 Teases a lot=94 Teases a lot_1                               2.714816e-02
## 19 Demands a lot\nof attention=19 Demands a lot\nof attention_2 2.430128e-02
## 3 Argues a lot=3 Argues a lot_0                                 1.895749e-02
## 22 Disobedient\nat home=22 Disobedient\nat home_2               1.823104e-02
## 87 Mood changes=87 Mood changes_2                               1.731065e-02
## 97 Threatens\nothers=97 Threatens\nothers_0                     1.525952e-02
## 3 Argues a lot=3 Argues a lot_2                                 5.883254e-03
## 21 Destroys\nothers things=21 Destroys\nothers things_1         2.578068e-03
## 97 Threatens\nothers=97 Threatens\nothers_1                     1.545741e-03
## 104 Loud=104 Loud_2                                             8.500470e-04
## 88 Sulks=88 Sulks_0                                             7.704590e-04
## 19 Demands a lot\nof attention=19 Demands a lot\nof attention_0 2.889174e-04
## 87 Mood changes=87 Mood changes_0                               1.530162e-04
## 86 Stubborn,\nsullen=86 Stubborn,\nsullen_0                     3.044104e-05
## 68 Screams a lot=68 Screams a lot_2                             1.273029e-06
## 20 Destroys own\nthings=20 Destroys own\nthings_1               9.925540e-07
## 89 Suspicious=89 Suspicious_2                                   5.974329e-07
## 
## attr(,"class")
## [1] "condes" "list"  
## 
## $`Dim 4`
## $quali
##                                        R2      p.value
## 87 Mood changes                0.24468246 8.124388e-12
## 94 Teases a lot                0.23790238 1.832168e-11
## 88 Sulks                       0.21589076 2.444938e-10
## 86 Stubborn,\nsullen           0.15639783 1.899020e-07
## 16 Cruel to\nothers            0.14428020 6.953023e-07
## 89 Suspicious                  0.12161064 7.507979e-06
## 37 Gets in\nfights             0.12041994 8.492958e-06
## 23 Disobedient\nat school      0.09151313 1.610637e-04
## 22 Disobedient\nat home        0.06470021 2.272615e-03
## 19 Demands a lot\nof attention 0.05805478 4.328463e-03
## 3 Argues a lot                 0.04739018 1.205785e-02
## 57 Attacks\nothers             0.04462128 1.570278e-02
## 95 Temper                      0.04141215 2.130613e-02
## 
## $category
##                                                                      Estimate
## 94 Teases a lot=94 Teases a lot_1                                0.1177611860
## 87 Mood changes=87 Mood changes_0                                0.2025818241
## 88 Sulks=88 Sulks_0                                              0.1940785032
## 16 Cruel to\nothers=16 Cruel to\nothers_1                        0.1032012361
## 37 Gets in\nfights=37 Gets in\nfights_1                          0.1380766543
## 89 Suspicious=89 Suspicious_0                                    0.1564149911
## 86 Stubborn,\nsullen=86 Stubborn,\nsullen_1                      0.0620196638
## 23 Disobedient\nat school=23 Disobedient\nat school_2            0.1386576965
## 3 Argues a lot=3 Argues a lot_2                                  0.0962546375
## 22 Disobedient\nat home=22 Disobedient\nat home_1                0.0833206770
## 57 Attacks\nothers=57 Attacks\nothers_1                          0.0666077075
## 19 Demands a lot\nof attention=19 Demands a lot\nof attention_0  0.1159681122
## 86 Stubborn,\nsullen=86 Stubborn,\nsullen_0                      0.1069966331
## 95 Temper=95 Temper_0                                            0.0662154529
## 68 Screams a lot=68 Screams a lot_1                              0.0682996320
## 89 Suspicious=89 Suspicious_1                                   -0.0005107064
## 95 Temper=95 Temper_2                                           -0.0783311263
## 19 Demands a lot\nof attention=19 Demands a lot\nof attention_1 -0.1070664477
## 57 Attacks\nothers=57 Attacks\nothers_0                         -0.0831329605
## 22 Disobedient\nat home=22 Disobedient\nat home_0               -0.1091876644
## 23 Disobedient\nat school=23 Disobedient\nat school_0           -0.1214222857
## 89 Suspicious=89 Suspicious_2                                   -0.1559042847
## 37 Gets in\nfights=37 Gets in\nfights_0                         -0.1199944288
## 88 Sulks=88 Sulks_2                                             -0.1763513523
## 16 Cruel to\nothers=16 Cruel to\nothers_0                       -0.1421105271
## 86 Stubborn,\nsullen=86 Stubborn,\nsullen_2                     -0.1690162970
## 87 Mood changes=87 Mood changes_2                               -0.1790867543
## 94 Teases a lot=94 Teases a lot_0                               -0.2064848948
##                                                                      p.value
## 94 Teases a lot=94 Teases a lot_1                               6.524391e-11
## 87 Mood changes=87 Mood changes_0                               3.069219e-10
## 88 Sulks=88 Sulks_0                                             4.414344e-09
## 16 Cruel to\nothers=16 Cruel to\nothers_1                       1.564728e-06
## 37 Gets in\nfights=37 Gets in\nfights_1                         2.147490e-06
## 89 Suspicious=89 Suspicious_0                                   9.996791e-06
## 86 Stubborn,\nsullen=86 Stubborn,\nsullen_1                     1.376992e-04
## 23 Disobedient\nat school=23 Disobedient\nat school_2           3.506978e-04
## 3 Argues a lot=3 Argues a lot_2                                 2.957207e-03
## 22 Disobedient\nat home=22 Disobedient\nat home_1               6.847634e-03
## 57 Attacks\nothers=57 Attacks\nothers_1                         7.300518e-03
## 19 Demands a lot\nof attention=19 Demands a lot\nof attention_0 8.736364e-03
## 86 Stubborn,\nsullen=86 Stubborn,\nsullen_0                     2.046790e-02
## 95 Temper=95 Temper_0                                           2.729873e-02
## 68 Screams a lot=68 Screams a lot_1                             3.787249e-02
## 89 Suspicious=89 Suspicious_1                                   2.195654e-02
## 95 Temper=95 Temper_2                                           9.068816e-03
## 19 Demands a lot\nof attention=19 Demands a lot\nof attention_1 7.785673e-03
## 57 Attacks\nothers=57 Attacks\nothers_0                         4.464613e-03
## 22 Disobedient\nat home=22 Disobedient\nat home_0               9.239078e-04
## 23 Disobedient\nat school=23 Disobedient\nat school_0           5.126771e-04
## 89 Suspicious=89 Suspicious_2                                   3.483335e-04
## 37 Gets in\nfights=37 Gets in\nfights_0                         3.639304e-06
## 88 Sulks=88 Sulks_2                                             6.483757e-07
## 16 Cruel to\nothers=16 Cruel to\nothers_0                       1.334364e-07
## 86 Stubborn,\nsullen=86 Stubborn,\nsullen_2                     3.213491e-08
## 87 Mood changes=87 Mood changes_2                               9.111706e-09
## 94 Teases a lot=94 Teases a lot_0                               1.962453e-12
## 
## attr(,"class")
## [1] "condes" "list"  
## 
## $`Dim 5`
## $quali
##                                    R2      p.value
## 20 Destroys own\nthings    0.31480428 1.145862e-15
## 21 Destroys\nothers things 0.26190886 9.954265e-13
## 37 Gets in\nfights         0.11073361 2.300917e-05
## 22 Disobedient\nat home    0.11030046 2.405173e-05
## 3 Argues a lot             0.10703064 3.358354e-05
## 97 Threatens\nothers       0.10208273 5.552718e-05
## 23 Disobedient\nat school  0.08792052 2.306610e-04
## 88 Sulks                   0.07771607 6.348667e-04
## 89 Suspicious              0.07470774 8.538518e-04
## 57 Attacks\nothers         0.05807623 4.319506e-03
## 68 Screams a lot           0.05203576 7.728016e-03
## 16 Cruel to\nothers        0.05157810 8.075011e-03
## 94 Teases a lot            0.05079397 8.705715e-03
## 
## $category
##                                                            Estimate
## 20 Destroys own\nthings=20 Destroys own\nthings_0        0.26903949
## 21 Destroys\nothers things=21 Destroys\nothers things_0  0.12839023
## 97 Threatens\nothers=97 Threatens\nothers_2              0.49102666
## 22 Disobedient\nat home=22 Disobedient\nat home_2        0.09695691
## 94 Teases a lot=94 Teases a lot_1                        0.09704688
## 3 Argues a lot=3 Argues a lot_0                          0.08468241
## 88 Sulks=88 Sulks_1                                      0.07987528
## 16 Cruel to\nothers=16 Cruel to\nothers_2                0.13818022
## 23 Disobedient\nat school=23 Disobedient\nat school_2    0.11882587
## 68 Screams a lot=68 Screams a lot_1                      0.08466210
## 89 Suspicious=89 Suspicious_1                            0.02842260
## 37 Gets in\nfights=37 Gets in\nfights_1                  0.20354262
## 89 Suspicious=89 Suspicious_2                            0.08693906
## 95 Temper=95 Temper_1                                    0.06563586
## 57 Attacks\nothers=57 Attacks\nothers_1                  0.13579849
## 16 Cruel to\nothers=16 Cruel to\nothers_0               -0.10116307
## 68 Screams a lot=68 Screams a lot_2                     -0.08812585
## 57 Attacks\nothers=57 Attacks\nothers_2                 -0.18436515
## 94 Teases a lot=94 Teases a lot_0                       -0.05082944
## 89 Suspicious=89 Suspicious_0                           -0.11536166
## 23 Disobedient\nat school=23 Disobedient\nat school_1   -0.13033808
## 88 Sulks=88 Sulks_0                                     -0.10831191
## 37 Gets in\nfights=37 Gets in\nfights_2                 -0.30672616
## 22 Disobedient\nat home=22 Disobedient\nat home_1       -0.12259751
## 3 Argues a lot=3 Argues a lot_1                         -0.12637271
## 20 Destroys own\nthings=20 Destroys own\nthings_2       -0.23894019
## 20 Destroys own\nthings=20 Destroys own\nthings_1       -0.03009929
## 21 Destroys\nothers things=21 Destroys\nothers things_1 -0.19401095
##                                                              p.value
## 20 Destroys own\nthings=20 Destroys own\nthings_0       2.192269e-15
## 21 Destroys\nothers things=21 Destroys\nothers things_0 7.202069e-13
## 97 Threatens\nothers=97 Threatens\nothers_2             1.641500e-05
## 22 Disobedient\nat home=22 Disobedient\nat home_2       3.083719e-04
## 94 Teases a lot=94 Teases a lot_1                       2.041027e-03
## 3 Argues a lot=3 Argues a lot_0                         2.535780e-03
## 88 Sulks=88 Sulks_1                                     3.124079e-03
## 16 Cruel to\nothers=16 Cruel to\nothers_2               5.437266e-03
## 23 Disobedient\nat school=23 Disobedient\nat school_2   6.333015e-03
## 68 Screams a lot=68 Screams a lot_1                     8.523135e-03
## 89 Suspicious=89 Suspicious_1                           1.423976e-02
## 37 Gets in\nfights=37 Gets in\nfights_1                 1.912342e-02
## 89 Suspicious=89 Suspicious_2                           2.914585e-02
## 95 Temper=95 Temper_1                                   3.532024e-02
## 57 Attacks\nothers=57 Attacks\nothers_1                 3.971541e-02
## 16 Cruel to\nothers=16 Cruel to\nothers_0               2.244947e-02
## 68 Screams a lot=68 Screams a lot_2                     8.424039e-03
## 57 Attacks\nothers=57 Attacks\nothers_2                 4.821508e-03
## 94 Teases a lot=94 Teases a lot_0                       4.260491e-03
## 89 Suspicious=89 Suspicious_0                           2.310486e-04
## 23 Disobedient\nat school=23 Disobedient\nat school_1   2.220575e-04
## 88 Sulks=88 Sulks_0                                     2.057117e-04
## 37 Gets in\nfights=37 Gets in\nfights_2                 2.949564e-05
## 22 Disobedient\nat home=22 Disobedient\nat home_1       8.764568e-06
## 3 Argues a lot=3 Argues a lot_1                         7.555172e-06
## 20 Destroys own\nthings=20 Destroys own\nthings_2       6.090039e-06
## 20 Destroys own\nthings=20 Destroys own\nthings_1       1.280517e-08
## 21 Destroys\nothers things=21 Destroys\nothers things_1 1.096286e-13
## 
## attr(,"class")
## [1] "condes" "list"  
## 
## $`Dim 6`
## $quali
##                                        R2      p.value
## 89 Suspicious                  0.20637157 7.331061e-10
## 86 Stubborn,\nsullen           0.14759661 4.883266e-07
## 104 Loud                       0.14152981 9.310970e-07
## 87 Mood changes                0.14133237 9.507874e-07
## 16 Cruel to\nothers            0.13224201 2.478990e-06
## 88 Sulks                       0.11306307 1.812337e-05
## 23 Disobedient\nat school      0.09480627 1.157390e-04
## 97 Threatens\nothers           0.07558891 7.829398e-04
## 68 Screams a lot               0.06926357 1.456233e-03
## 20 Destroys own\nthings        0.06164476 3.057897e-03
## 19 Demands a lot\nof attention 0.04452189 1.585213e-02
## 22 Disobedient\nat home        0.03968790 2.509160e-02
## 
## $category
##                                                                     Estimate
## 89 Suspicious=89 Suspicious_2                                    0.250470047
## 16 Cruel to\nothers=16 Cruel to\nothers_1                        0.166886017
## 88 Sulks=88 Sulks_2                                              0.141928954
## 86 Stubborn,\nsullen=86 Stubborn,\nsullen_2                      0.081981304
## 87 Mood changes=87 Mood changes_0                                0.116698817
## 23 Disobedient\nat school=23 Disobedient\nat school_1            0.070119170
## 104 Loud=104 Loud_1                                              0.095735443
## 22 Disobedient\nat home=22 Disobedient\nat home_1                0.068586055
## 68 Screams a lot=68 Screams a lot_0                              0.076375635
## 3 Argues a lot=3 Argues a lot_1                                  0.062977828
## 20 Destroys own\nthings=20 Destroys own\nthings_0                0.008347448
## 94 Teases a lot=94 Teases a lot_2                                0.147062723
## 19 Demands a lot\nof attention=19 Demands a lot\nof attention_1  0.035703454
## 21 Destroys\nothers things=21 Destroys\nothers things_1         -0.090539755
## 22 Disobedient\nat home=22 Disobedient\nat home_2               -0.061188922
## 95 Temper=95 Temper_2                                           -0.063930819
## 16 Cruel to\nothers=16 Cruel to\nothers_2                       -0.136259625
## 88 Sulks=88 Sulks_1                                             -0.083895229
## 19 Demands a lot\nof attention=19 Demands a lot\nof attention_2 -0.079871648
## 20 Destroys own\nthings=20 Destroys own\nthings_1               -0.127853798
## 16 Cruel to\nothers=16 Cruel to\nothers_0                       -0.030626393
## 68 Screams a lot=68 Screams a lot_2                             -0.110596779
## 97 Threatens\nothers=97 Threatens\nothers_2                     -0.405617146
## 23 Disobedient\nat school=23 Disobedient\nat school_0           -0.110785848
## 89 Suspicious=89 Suspicious_0                                   -0.175621766
## 87 Mood changes=87 Mood changes_1                               -0.147991389
## 104 Loud=104 Loud_2                                             -0.163376975
## 86 Stubborn,\nsullen=86 Stubborn,\nsullen_1                     -0.139312636
##                                                                      p.value
## 89 Suspicious=89 Suspicious_2                                   1.678865e-09
## 16 Cruel to\nothers=16 Cruel to\nothers_1                       1.088610e-06
## 88 Sulks=88 Sulks_2                                             3.395461e-06
## 86 Stubborn,\nsullen=86 Stubborn,\nsullen_2                     1.218478e-05
## 87 Mood changes=87 Mood changes_0                               4.778611e-04
## 23 Disobedient\nat school=23 Disobedient\nat school_1           7.326345e-04
## 104 Loud=104 Loud_1                                             7.671925e-03
## 22 Disobedient\nat home=22 Disobedient\nat home_1               1.079477e-02
## 68 Screams a lot=68 Screams a lot_0                             1.805730e-02
## 3 Argues a lot=3 Argues a lot_1                                 2.198739e-02
## 20 Destroys own\nthings=20 Destroys own\nthings_0               2.736868e-02
## 94 Teases a lot=94 Teases a lot_2                               4.597868e-02
## 19 Demands a lot\nof attention=19 Demands a lot\nof attention_1 4.636702e-02
## 21 Destroys\nothers things=21 Destroys\nothers things_1         3.275512e-02
## 22 Disobedient\nat home=22 Disobedient\nat home_2               2.295105e-02
## 95 Temper=95 Temper_2                                           2.274783e-02
## 16 Cruel to\nothers=16 Cruel to\nothers_2                       2.095037e-02
## 88 Sulks=88 Sulks_1                                             5.356514e-03
## 19 Demands a lot\nof attention=19 Demands a lot\nof attention_2 3.978051e-03
## 20 Destroys own\nthings=20 Destroys own\nthings_1               1.491989e-03
## 16 Cruel to\nothers=16 Cruel to\nothers_0                       1.112276e-03
## 68 Screams a lot=68 Screams a lot_2                             4.620217e-04
## 97 Threatens\nothers=97 Threatens\nothers_2                     2.525196e-04
## 23 Disobedient\nat school=23 Disobedient\nat school_0           2.270563e-05
## 89 Suspicious=89 Suspicious_0                                   8.413169e-06
## 87 Mood changes=87 Mood changes_1                               6.350390e-07
## 104 Loud=104 Loud_2                                             1.612594e-07
## 86 Stubborn,\nsullen=86 Stubborn,\nsullen_1                     7.288181e-08
## 
## attr(,"class")
## [1] "condes" "list"  
## 
## $`Dim 7`
## $quali
##                                        R2      p.value
## 57 Attacks\nothers             0.21856972 1.790653e-10
## 89 Suspicious                  0.14378646 7.327748e-07
## 37 Gets in\nfights             0.12246479 6.871857e-06
## 95 Temper                      0.10981765 2.526895e-05
## 16 Cruel to\nothers            0.09643645 9.822949e-05
## 22 Disobedient\nat home        0.07848962 5.881944e-04
## 94 Teases a lot                0.07820388 6.050254e-04
## 104 Loud                       0.07562194 7.803984e-04
## 19 Demands a lot\nof attention 0.06702686 1.811723e-03
## 20 Destroys own\nthings        0.06674683 1.861882e-03
## 86 Stubborn,\nsullen           0.05810920 4.305765e-03
## 23 Disobedient\nat school      0.05230067 7.533938e-03
## 88 Sulks                       0.03953579 2.545587e-02
## 
## $category
##                                                                    Estimate
## 57 Attacks\nothers=57 Attacks\nothers_1                          0.22216839
## 89 Suspicious=89 Suspicious_1                                    0.16928504
## 37 Gets in\nfights=37 Gets in\nfights_1                          0.19366972
## 16 Cruel to\nothers=16 Cruel to\nothers_0                        0.13089605
## 23 Disobedient\nat school=23 Disobedient\nat school_1            0.08458474
## 95 Temper=95 Temper_2                                            0.07900100
## 22 Disobedient\nat home=22 Disobedient\nat home_1                0.06640130
## 88 Sulks=88 Sulks_1                                              0.06845556
## 20 Destroys own\nthings=20 Destroys own\nthings_2                0.15027971
## 104 Loud=104 Loud_1                                              0.04921495
## 86 Stubborn,\nsullen=86 Stubborn,\nsullen_2                      0.02881419
## 21 Destroys\nothers things=21 Destroys\nothers things_2          0.19845524
## 3 Argues a lot=3 Argues a lot_0                                  0.06395507
## 97 Threatens\nothers=97 Threatens\nothers_0                     -0.06258033
## 89 Suspicious=89 Suspicious_0                                   -0.00186465
## 88 Sulks=88 Sulks_2                                             -0.06210540
## 37 Gets in\nfights=37 Gets in\nfights_2                         -0.18021059
## 23 Disobedient\nat school=23 Disobedient\nat school_0           -0.04953344
## 94 Teases a lot=94 Teases a lot_0                               -0.00980723
## 20 Destroys own\nthings=20 Destroys own\nthings_1               -0.13453788
## 37 Gets in\nfights=37 Gets in\nfights_0                         -0.01345913
## 86 Stubborn,\nsullen=86 Stubborn,\nsullen_1                     -0.09218620
## 19 Demands a lot\nof attention=19 Demands a lot\nof attention_1 -0.10592513
## 89 Suspicious=89 Suspicious_2                                   -0.16742039
## 94 Teases a lot=94 Teases a lot_1                               -0.15682832
## 16 Cruel to\nothers=16 Cruel to\nothers_2                       -0.15981340
## 104 Loud=104 Loud_0                                             -0.10090485
## 22 Disobedient\nat home=22 Disobedient\nat home_2               -0.10921008
## 95 Temper=95 Temper_1                                           -0.12862325
## 57 Attacks\nothers=57 Attacks\nothers_0                         -0.06872443
##                                                                      p.value
## 57 Attacks\nothers=57 Attacks\nothers_1                         3.724815e-11
## 89 Suspicious=89 Suspicious_1                                   3.656449e-06
## 37 Gets in\nfights=37 Gets in\nfights_1                         5.587046e-06
## 16 Cruel to\nothers=16 Cruel to\nothers_0                       4.703300e-04
## 23 Disobedient\nat school=23 Disobedient\nat school_1           1.804274e-03
## 95 Temper=95 Temper_2                                           3.981431e-03
## 22 Disobedient\nat home=22 Disobedient\nat home_1               7.863007e-03
## 88 Sulks=88 Sulks_1                                             1.258127e-02
## 20 Destroys own\nthings=20 Destroys own\nthings_2               2.170553e-02
## 104 Loud=104 Loud_1                                             2.539202e-02
## 86 Stubborn,\nsullen=86 Stubborn,\nsullen_2                     2.934757e-02
## 21 Destroys\nothers things=21 Destroys\nothers things_2         3.427995e-02
## 3 Argues a lot=3 Argues a lot_0                                 4.285142e-02
## 97 Threatens\nothers=97 Threatens\nothers_0                     4.674448e-02
## 89 Suspicious=89 Suspicious_0                                   3.373869e-02
## 88 Sulks=88 Sulks_2                                             2.969056e-02
## 37 Gets in\nfights=37 Gets in\nfights_2                         2.949601e-02
## 23 Disobedient\nat school=23 Disobedient\nat school_0           1.497991e-02
## 94 Teases a lot=94 Teases a lot_0                               9.093895e-03
## 20 Destroys own\nthings=20 Destroys own\nthings_1               2.784129e-03
## 37 Gets in\nfights=37 Gets in\nfights_0                         1.522794e-03
## 86 Stubborn,\nsullen=86 Stubborn,\nsullen_1                     1.118183e-03
## 19 Demands a lot\nof attention=19 Demands a lot\nof attention_1 6.217903e-04
## 89 Suspicious=89 Suspicious_2                                   5.628104e-04
## 94 Teases a lot=94 Teases a lot_1                               4.635301e-04
## 16 Cruel to\nothers=16 Cruel to\nothers_2                       3.862714e-04
## 104 Loud=104 Loud_0                                             1.518883e-04
## 22 Disobedient\nat home=22 Disobedient\nat home_2               1.276313e-04
## 95 Temper=95 Temper_1                                           5.076481e-06
## 57 Attacks\nothers=57 Attacks\nothers_0                         6.208942e-07
## 
## attr(,"class")
## [1] "condes" "list"  
## 
## $`Dim 8`
## $quali
##                                   R2      p.value
## 97 Threatens\nothers      0.45259536 1.534869e-24
## 104 Loud                  0.11501354 1.483314e-05
## 37 Gets in\nfights        0.09951171 7.202846e-05
## 89 Suspicious             0.09608882 1.017288e-04
## 57 Attacks\nothers        0.08375008 3.493570e-04
## 3 Argues a lot            0.07525013 8.094857e-04
## 88 Sulks                  0.07313495 9.965572e-04
## 16 Cruel to\nothers       0.05378994 6.529323e-03
## 23 Disobedient\nat school 0.04261351 1.900833e-02
## 
## $category
##                                                          Estimate      p.value
## 97 Threatens\nothers=97 Threatens\nothers_2            0.81595352 4.946712e-16
## 89 Suspicious=89 Suspicious_1                          0.12507428 2.231775e-05
## 3 Argues a lot=3 Argues a lot_1                        0.09511653 1.711501e-04
## 104 Loud=104 Loud_2                                    0.09797499 1.315361e-03
## 57 Attacks\nothers=57 Attacks\nothers_0                0.04813579 1.514750e-03
## 16 Cruel to\nothers=16 Cruel to\nothers_1              0.07862294 1.613849e-03
## 23 Disobedient\nat school=23 Disobedient\nat school_1  0.08022170 5.865760e-03
## 20 Destroys own\nthings=20 Destroys own\nthings_2      0.11453742 3.996347e-02
## 88 Sulks=88 Sulks_2                                    0.05219898 4.188075e-02
## 88 Sulks=88 Sulks_0                                    0.04238917 4.458371e-02
## 3 Argues a lot=3 Argues a lot_0                       -0.05731403 1.978504e-02
## 16 Cruel to\nothers=16 Cruel to\nothers_0             -0.05270035 4.486851e-03
## 89 Suspicious=89 Suspicious_0                         -0.04168115 3.187767e-03
## 97 Threatens\nothers=97 Threatens\nothers_0           -0.26970876 2.999170e-03
## 88 Sulks=88 Sulks_1                                   -0.09458815 2.010614e-04
## 57 Attacks\nothers=57 Attacks\nothers_1               -0.12773209 7.032374e-05
## 37 Gets in\nfights=37 Gets in\nfights_2               -0.26784502 2.827461e-05
## 104 Loud=104 Loud_1                                   -0.12221529 8.679849e-06
## 97 Threatens\nothers=97 Threatens\nothers_1           -0.54624476 3.548491e-09
## 
## attr(,"class")
## [1] "condes" "list"  
## 
## $`Dim 9`
## $quali
##                                        R2      p.value
## 89 Suspicious                  0.21221523 3.741879e-10
## 23 Disobedient\nat school      0.13103499 2.813251e-06
## 94 Teases a lot                0.10694565 3.387568e-05
## 21 Destroys\nothers things     0.10467373 4.268752e-05
## 19 Demands a lot\nof attention 0.10269139 5.220439e-05
## 37 Gets in\nfights             0.09239591 1.474267e-04
## 68 Screams a lot               0.09117543 1.666040e-04
## 86 Stubborn,\nsullen           0.08977179 1.917239e-04
## 87 Mood changes                0.05245173 7.425439e-03
## 57 Attacks\nothers             0.04582135 1.400562e-02
## 97 Threatens\nothers           0.04062675 2.295473e-02
## 104 Loud                       0.03296577 4.733779e-02
## 
## $category
##                                                                     Estimate
## 23 Disobedient\nat school=23 Disobedient\nat school_1            0.139584204
## 94 Teases a lot=94 Teases a lot_2                                0.303799928
## 89 Suspicious=89 Suspicious_2                                    0.186233751
## 86 Stubborn,\nsullen=86 Stubborn,\nsullen_1                      0.110494613
## 19 Demands a lot\nof attention=19 Demands a lot\nof attention_1  0.123039908
## 21 Destroys\nothers things=21 Destroys\nothers things_2          0.315443241
## 37 Gets in\nfights=37 Gets in\nfights_1                          0.167755864
## 68 Screams a lot=68 Screams a lot_2                              0.099415598
## 87 Mood changes=87 Mood changes_1                                0.070369187
## 89 Suspicious=89 Suspicious_0                                    0.007741786
## 97 Threatens\nothers=97 Threatens\nothers_1                      0.083065301
## 95 Temper=95 Temper_1                                            0.062995299
## 104 Loud=104 Loud_2                                              0.063122370
## 37 Gets in\nfights=37 Gets in\nfights_0                          0.018908281
## 94 Teases a lot=94 Teases a lot_1                               -0.184406441
## 23 Disobedient\nat school=23 Disobedient\nat school_0           -0.017466634
## 104 Loud=104 Loud_1                                             -0.058703161
## 88 Sulks=88 Sulks_1                                             -0.051582333
## 86 Stubborn,\nsullen=86 Stubborn,\nsullen_2                     -0.021609536
## 86 Stubborn,\nsullen=86 Stubborn,\nsullen_0                     -0.088885077
## 37 Gets in\nfights=37 Gets in\nfights_2                         -0.186664144
## 97 Threatens\nothers=97 Threatens\nothers_0                     -0.058284826
## 87 Mood changes=87 Mood changes_2                               -0.064380211
## 57 Attacks\nothers=57 Attacks\nothers_2                         -0.139982033
## 21 Destroys\nothers things=21 Destroys\nothers things_1         -0.211104263
## 19 Demands a lot\nof attention=19 Demands a lot\nof attention_0 -0.118365487
## 23 Disobedient\nat school=23 Disobedient\nat school_2           -0.122117570
## 68 Screams a lot=68 Screams a lot_1                             -0.099808312
## 89 Suspicious=89 Suspicious_1                                   -0.193975537
##                                                                      p.value
## 23 Disobedient\nat school=23 Disobedient\nat school_1           3.493944e-06
## 94 Teases a lot=94 Teases a lot_2                               2.132788e-05
## 89 Suspicious=89 Suspicious_2                                   4.903206e-05
## 86 Stubborn,\nsullen=86 Stubborn,\nsullen_1                     6.937533e-05
## 19 Demands a lot\nof attention=19 Demands a lot\nof attention_1 1.032971e-04
## 21 Destroys\nothers things=21 Destroys\nothers things_2         3.654020e-04
## 37 Gets in\nfights=37 Gets in\nfights_1                         4.259707e-04
## 68 Screams a lot=68 Screams a lot_2                             6.372650e-04
## 87 Mood changes=87 Mood changes_1                               3.997726e-03
## 89 Suspicious=89 Suspicious_0                                   5.536736e-03
## 97 Threatens\nothers=97 Threatens\nothers_1                     6.112142e-03
## 95 Temper=95 Temper_1                                           2.255173e-02
## 104 Loud=104 Loud_2                                             3.368104e-02
## 37 Gets in\nfights=37 Gets in\nfights_0                         3.977780e-02
## 94 Teases a lot=94 Teases a lot_1                               4.710511e-02
## 23 Disobedient\nat school=23 Disobedient\nat school_0           4.646738e-02
## 104 Loud=104 Loud_1                                             4.261329e-02
## 88 Sulks=88 Sulks_1                                             4.241632e-02
## 86 Stubborn,\nsullen=86 Stubborn,\nsullen_2                     1.848918e-02
## 86 Stubborn,\nsullen=86 Stubborn,\nsullen_0                     1.661373e-02
## 37 Gets in\nfights=37 Gets in\nfights_2                         9.437511e-03
## 97 Threatens\nothers=97 Threatens\nothers_0                     7.844509e-03
## 87 Mood changes=87 Mood changes_2                               6.550658e-03
## 57 Attacks\nothers=57 Attacks\nothers_2                         5.265153e-03
## 21 Destroys\nothers things=21 Destroys\nothers things_1         2.798426e-03
## 19 Demands a lot\nof attention=19 Demands a lot\nof attention_0 1.742391e-03
## 23 Disobedient\nat school=23 Disobedient\nat school_2           1.404199e-03
## 68 Screams a lot=68 Screams a lot_1                             3.184066e-04
## 89 Suspicious=89 Suspicious_1                                   6.015447e-09
## 
## attr(,"class")
## [1] "condes" "list"  
## 
## $`Dim 10`
## $quali
##                                        R2      p.value
## 68 Screams a lot               0.18323157 1.002314e-08
## 22 Disobedient\nat home        0.14773700 4.810618e-07
## 87 Mood changes                0.12318111 6.379706e-06
## 95 Temper                      0.11846048 1.039926e-05
## 89 Suspicious                  0.08776794 2.341992e-04
## 37 Gets in\nfights             0.07441836 8.784978e-04
## 23 Disobedient\nat school      0.07182590 1.133130e-03
## 16 Cruel to\nothers            0.05180485 7.901208e-03
## 19 Demands a lot\nof attention 0.04342803 1.759164e-02
## 94 Teases a lot                0.03915660 2.638685e-02
## 20 Destroys own\nthings        0.03717299 3.183408e-02
## 
## $category
##                                                                      Estimate
## 87 Mood changes=87 Mood changes_1                                0.1237280710
## 22 Disobedient\nat home=22 Disobedient\nat home_0                0.1193763604
## 68 Screams a lot=68 Screams a lot_0                              0.0958258136
## 89 Suspicious=89 Suspicious_1                                    0.1039611021
## 37 Gets in\nfights=37 Gets in\nfights_1                          0.0573960986
## 95 Temper=95 Temper_0                                            0.0935886699
## 23 Disobedient\nat school=23 Disobedient\nat school_2            0.1142409817
## 16 Cruel to\nothers=16 Cruel to\nothers_1                        0.0966983781
## 19 Demands a lot\nof attention=19 Demands a lot\nof attention_1  0.0781927143
## 94 Teases a lot=94 Teases a lot_1                                0.0984574602
## 20 Destroys own\nthings=20 Destroys own\nthings_1                0.0428782880
## 86 Stubborn,\nsullen=86 Stubborn,\nsullen_1                      0.0646782381
## 94 Teases a lot=94 Teases a lot_0                               -0.0005054165
## 97 Threatens\nothers=97 Threatens\nothers_2                     -0.1892092486
## 16 Cruel to\nothers=16 Cruel to\nothers_2                       -0.1040411281
## 87 Mood changes=87 Mood changes_2                               -0.0450559484
## 23 Disobedient\nat school=23 Disobedient\nat school_1           -0.0916017242
## 87 Mood changes=87 Mood changes_0                               -0.0786721226
## 20 Destroys own\nthings=20 Destroys own\nthings_0               -0.0615281278
## 89 Suspicious=89 Suspicious_0                                   -0.0569790395
## 37 Gets in\nfights=37 Gets in\nfights_0                         -0.1000725165
## 95 Temper=95 Temper_1                                           -0.1210407367
## 22 Disobedient\nat home=22 Disobedient\nat home_1               -0.1154970156
## 68 Screams a lot=68 Screams a lot_1                             -0.1398628222
##                                                                      p.value
## 87 Mood changes=87 Mood changes_1                               1.313908e-06
## 22 Disobedient\nat home=22 Disobedient\nat home_0               3.241196e-06
## 68 Screams a lot=68 Screams a lot_0                             8.966269e-06
## 89 Suspicious=89 Suspicious_1                                   4.294814e-05
## 37 Gets in\nfights=37 Gets in\nfights_1                         7.141594e-04
## 95 Temper=95 Temper_0                                           9.881427e-04
## 23 Disobedient\nat school=23 Disobedient\nat school_2           1.161329e-03
## 16 Cruel to\nothers=16 Cruel to\nothers_1                       8.406328e-03
## 19 Demands a lot\nof attention=19 Demands a lot\nof attention_1 1.049750e-02
## 94 Teases a lot=94 Teases a lot_1                               1.154388e-02
## 20 Destroys own\nthings=20 Destroys own\nthings_1               1.495504e-02
## 86 Stubborn,\nsullen=86 Stubborn,\nsullen_1                     2.326261e-02
## 94 Teases a lot=94 Teases a lot_0                               4.949443e-02
## 97 Threatens\nothers=97 Threatens\nothers_2                     3.760023e-02
## 16 Cruel to\nothers=16 Cruel to\nothers_2                       3.277699e-02
## 87 Mood changes=87 Mood changes_2                               2.210732e-02
## 23 Disobedient\nat school=23 Disobedient\nat school_1           9.703014e-03
## 87 Mood changes=87 Mood changes_0                               9.610598e-03
## 20 Destroys own\nthings=20 Destroys own\nthings_0               8.958253e-03
## 89 Suspicious=89 Suspicious_0                                   8.421011e-04
## 37 Gets in\nfights=37 Gets in\nfights_0                         1.744033e-04
## 95 Temper=95 Temper_1                                           6.100848e-06
## 22 Disobedient\nat home=22 Disobedient\nat home_1               2.082543e-06
## 68 Screams a lot=68 Screams a lot_1                             2.575441e-09
## 
## attr(,"class")
## [1] "condes" "list"
```

Write MCA results to file

```
mca_coord <- get_mca_ind(mca_pheno)$coord
write.csv(mca_coord, params$cbcl_mca_coord_outfile)
save(mca_pheno, file = params$cbcl_mca_outfile)
```

```
sessionInfo()
```

```
## R version 4.1.2 (2021-11-01)
## Platform: x86_64-pc-linux-gnu (64-bit)
## Running under: Debian GNU/Linux bookworm/sid
## 
## Matrix products: default
## BLAS:   /usr/lib/x86_64-linux-gnu/openblas-pthread/libblas.so.3
## LAPACK: /usr/lib/x86_64-linux-gnu/openblas-pthread/libopenblasp-r0.3.19.so
## 
## locale:
##  [1] LC_CTYPE=en_US.UTF-8       LC_NUMERIC=C              
##  [3] LC_TIME=en_US.UTF-8        LC_COLLATE=en_US.UTF-8    
##  [5] LC_MONETARY=en_US.UTF-8    LC_MESSAGES=en_US.UTF-8   
##  [7] LC_PAPER=en_US.UTF-8       LC_NAME=C                 
##  [9] LC_ADDRESS=C               LC_TELEPHONE=C            
## [11] LC_MEASUREMENT=en_US.UTF-8 LC_IDENTIFICATION=C       
## 
## attached base packages:
## [1] stats     graphics  grDevices utils     datasets  methods   base     
## 
## other attached packages:
##  [1] factoextra_1.0.7    FactoMineR_2.4      ggplot2_3.3.5      
##  [4] stringr_1.4.0       reshape2_1.4.4      missForest_1.4     
##  [7] itertools_0.1-3     iterators_1.0.13    foreach_1.5.2      
## [10] randomForest_4.6-14
## 
## loaded via a namespace (and not attached):
##  [1] ggrepel_0.9.1        Rcpp_1.0.8           lattice_0.20-45     
##  [4] tidyr_1.2.0          digest_0.6.29        utf8_1.2.2          
##  [7] R6_2.5.1             plyr_1.8.6           backports_1.4.1     
## [10] evaluate_0.14        highr_0.9            pillar_1.7.0        
## [13] rlang_1.0.0          car_3.0-12           jquerylib_0.1.4     
## [16] DT_0.20              rmarkdown_2.11       labeling_0.4.2      
## [19] htmlwidgets_1.5.4    munsell_0.5.0        broom_0.7.12        
## [22] compiler_4.1.2       xfun_0.29            pkgconfig_2.0.3     
## [25] htmltools_0.5.2      flashClust_1.01-2    tidyselect_1.1.1    
## [28] tibble_3.1.6         codetools_0.2-18     fansi_1.0.2         
## [31] crayon_1.4.2         dplyr_1.0.8          withr_2.4.3         
## [34] ggpubr_0.4.0         MASS_7.3-55          leaps_3.1           
## [37] grid_4.1.2           jsonlite_1.7.3       gtable_0.3.0        
## [40] lifecycle_1.0.1      magrittr_2.0.2       scales_1.1.1        
## [43] carData_3.0-5        cli_3.1.1            stringi_1.7.6       
## [46] farver_2.1.0         ggsignif_0.6.3       scatterplot3d_0.3-41
## [49] bslib_0.3.1          ellipsis_0.3.2       generics_0.1.2      
## [52] vctrs_0.3.8          tools_4.1.2          glue_1.6.1          
## [55] purrr_0.3.4          abind_1.4-5          parallel_4.1.2      
## [58] fastmap_1.1.0        yaml_2.2.2           colorspace_2.0-2    
## [61] cluster_2.1.2        rstatix_0.7.0        knitr_1.37          
## [64] sass_0.4.0
```

---

1. Radboud University Medical Center,↩︎
