## Additional File 1 for "A Multi-omics Data Analysis Workflow Packaged as a FAIR Digital Object": Additional File 1.html

Multiple Correspondence analysis of CBCL behavioral data


### Multiple Correspondence analysis of CBCL behavioral data

###### Anna Niehues^1^

This notebook performs filtering based on missingness, missing-value imputation, and multiple correspondence analysis (MCA). Visualizations of data and results are plotted.

```
print(params)
```

```
## $missing_value_cutoff
## [1] 0.5
## 
## $cbcl_infile
## [1] "cbcl_data.csv"
## 
## $cbcl_labels
## [1] "cbcl_labels.csv"
## 
## $cbcl_filtered_outfile
## [1] "cbcl_filtered.csv"
## 
## $cbcl_imputed_outfile
## [1] "cbcl_imputed.csv"
## 
## $cbcl_mca_coord_outfile
## [1] "cbcl_mca_coord.csv"
## 
## $cbcl_mca_outfile
## [1] "cbcl_mca.RData"
## 
## $imputation_method
## [1] "RF"
```

Read phenotypic data (samples in rows, survey elements in columns).

```
# read phenotypic data
pheno_in <- read.csv(params$cbcl_infile, row.names = 1)
vars_in <- colnames(pheno_in)
# labels 
pheno_labels <- read.csv(params$cbcl_labels)
knitr::kable(pheno_labels)
```

| X | variable | label |
| --- | --- | --- |
| 1 | CBCL6-18\_q3 | CBCL6\_18 - 3 Argues a lot |
| 2 | CBCL6-18\_q16 | CBCL6\_18 - 16 Cruel to others |
| 3 | CBCL6-18\_q19 | CBCL6\_18 - 19 Demands a lot of attention |
| 4 | CBCL6-18\_q20 | CBCL6\_18 - 20 Destroys own things |
| 5 | CBCL6-18\_q21 | CBCL6\_18 - 21 Destroys others things |
| 6 | CBCL6-18\_q22 | CBCL6\_18 - 22 Disobedient at home |
| 7 | CBCL6-18\_q23 | CBCL6\_18 - 23 Disobedient at school |
| 8 | CBCL6-18\_q37 | CBCL6\_18 - 37 Gets in fights |
| 9 | CBCL6-18\_q57 | CBCL6\_18 - 57 Attacks others |
| 10 | CBCL6-18\_q68 | CBCL6\_18 - 68 Screams a lot |
| 11 | CBCL6-18\_q86 | CBCL6\_18 - 86 Stubborn, sullen |
| 12 | CBCL6-18\_q87 | CBCL6\_18 - 87 Mood changes |
| 13 | CBCL6-18\_q88 | CBCL6\_18 - 88 Sulks |
| 14 | CBCL6-18\_q89 | CBCL6\_18 - 89 Suspicious |
| 15 | CBCL6-18\_q94 | CBCL6\_18 - 94 Teases a lot |
| 16 | CBCL6-18\_q95 | CBCL6\_18 - 95 Temper |
| 17 | CBCL6-18\_q97 | CBCL6\_18 - 97 Threatens others |
| 18 | CBCL6-18\_q104 | CBCL6\_18 - 104 Loud |
| 19 | CBCL6-18\_q15 | CBCL6\_18 - 15 Cruel to animals |
| 20 | CBCL6-18\_q26 | CBCL6\_18 - 26 Lacks guilt |
| 21 | CBCL6-18\_q39 | CBCL6\_18 - 39 Bad friends |
| 22 | CBCL6-18\_q43 | CBCL6\_18 - 43 Lying |
| 23 | CBCL6-18\_q81 | CBCL6\_18 - 81 Steals at home |
| 24 | CBCL6-18\_q90 | CBCL6\_18 - 90 Swearing |
| 25 | CBCL6-18\_q101 | CBCL6\_18 - 101 Truant |
| 26 | CBCL6-18\_q106 | CBCL6\_18 - 106 Vandalism |

```
cat('\n\n')
```

```
# convert 999999 to NA
pheno_in[pheno_in == "999999"] <- NA
# convert to factor
pheno_in[, vars_in] <- lapply(pheno_in[, vars_in], as.factor)
```

#### Overview of input data

```
summary(pheno_in)
```

```
##  CBCL6.18_q3 CBCL6.18_q16 CBCL6.18_q19 CBCL6.18_q20 CBCL6.18_q21 CBCL6.18_q22
##  0   :589    0   :1131    0   :839     0   :1199    0   :1207    0   :737    
##  1   :645    1   : 207    1   :392     1   : 133    1   : 130    1   :560    
##  2   :113    2   :   9    2   :114     2   :  14    2   :   4    2   : 47    
##  NA's:157    NA's: 157    NA's:159     NA's: 158    NA's: 163    NA's:160    
##  CBCL6.18_q23 CBCL6.18_q37 CBCL6.18_q57 CBCL6.18_q68 CBCL6.18_q86 CBCL6.18_q87
##  0   :1135    0   :1221    0   :1266    0   :915     0   :695     0   :981    
##  1   : 200    1   : 112    1   :  68    1   :342     1   :515     1   :284    
##  2   :  11    2   :  15    2   :   7    2   : 82     2   :128     2   : 74    
##  NA's: 158    NA's: 156    NA's: 163    NA's:165     NA's:166     NA's:165    
##  CBCL6.18_q88 CBCL6.18_q89 CBCL6.18_q94 CBCL6.18_q95 CBCL6.18_q97 CBCL6.18_q104
##  0   :901     0   :1230    0   :1218    0   :928     0   :1317    0   :1071    
##  1   :379     1   :  97    1   : 107    1   :318     1   :  10    1   : 202    
##  2   : 55     2   :  11    2   :   7    2   : 85     2   :   1    2   :  61    
##  NA's:169     NA's: 166    NA's: 172    NA's:173     NA's: 176    NA's: 170    
##  CBCL6.18_q15 CBCL6.18_q26 CBCL6.18_q39 CBCL6.18_q43 CBCL6.18_q81 CBCL6.18_q90
##  0   :1306    0   :1078    0   :1248    0   :1081    0   :1317    0   :1032   
##  1   :  36    1   : 231    1   :  92    1   : 252    1   :  23    1   : 270   
##  2   :   7    2   :  37    2   :   9    2   :  15    2   :   3    2   :  35   
##  NA's: 155    NA's: 158    NA's: 155    NA's: 156    NA's: 161    NA's: 167   
##  CBCL6.18_q101 CBCL6.18_q106
##  0   :1332     0   :1337    
##  1   :   3     1   :   1    
##  2   :   1     2   :   2    
##  NA's: 168     NA's: 164
```

```
# distribution of missing values across samples and features
# histogram of missing values per sample
hist(rowSums(is.na(pheno_in)))
```

```
# histogram of missing values per feature
hist(colSums(is.na(pheno_in)))
```

#### Filter phenotypic data

Filter features: remove non-informative (invariant) features

```
pheno_filtfeat <- pheno_in[, sapply(pheno_in, nlevels) > 1]
message("[FILTERING] Removed ", dim(pheno_in)[2]-dim(pheno_filtfeat)[2], 
        " invariant feature(s).")
```

```
## [FILTERING] Removed 0 invariant feature(s).
```

Filter samples: remove all samples that have missing values across most features.

```
pheno_filtsamp <- pheno_filtfeat[
  rowSums(is.na(pheno_filtfeat)) < params$missing_value_cutoff*dim(pheno_filtfeat)[2],]
message("[FILTERING] Removed ", dim(pheno_filtfeat)[1]-dim(pheno_filtsamp)[1], 
        " samples with more than ", params$missing_value_cutoff*100, 
        "% missing values.")
```

```
## [FILTERING] Removed 155 samples with more than 50% missing values.
```

```
# remove invariant features
pheno_filt <- pheno_filtsamp[, sapply(pheno_filtsamp, nlevels) > 1]
message("[FILTERING] Removed ", dim(pheno_filtsamp)[2]-dim(pheno_filt)[2], 
        " invariant feature(s).")
```

```
## [FILTERING] Removed 0 invariant feature(s).
```

#### Overview of filtered data

```
message("[FILTERING] Filtered data set contains ", dim(pheno_filt)[1], 
        " samples, ", dim(pheno_filt)[2], " features, and ", 
        sum(is.na(pheno_filt)), " missing values.")
```

```
## [FILTERING] Filtered data set contains 1349 samples, 26 features, and 231 missing values.
```

```
print(summary(pheno_filt))
```

```
##  CBCL6.18_q3 CBCL6.18_q16 CBCL6.18_q19 CBCL6.18_q20 CBCL6.18_q21 CBCL6.18_q22
##  0   :588    0   :1129    0   :838     0   :1197    0   :1206    0   :735    
##  1   :644    1   : 207    1   :391     1   : 133    1   : 129    1   :560    
##  2   :113    2   :   9    2   :114     2   :  14    2   :   4    2   : 47    
##  NA's:  4    NA's:   4    NA's:  6     NA's:   5    NA's:  10    NA's:  7    
##  CBCL6.18_q23 CBCL6.18_q37 CBCL6.18_q57 CBCL6.18_q68 CBCL6.18_q86 CBCL6.18_q87
##  0   :1133    0   :1219    0   :1266    0   :915     0   :695     0   :981    
##  1   : 200    1   : 112    1   :  68    1   :342     1   :515     1   :284    
##  2   :  11    2   :  15    2   :   7    2   : 82     2   :128     2   : 74    
##  NA's:   5    NA's:   3    NA's:   8    NA's: 10     NA's: 11     NA's: 10    
##  CBCL6.18_q88 CBCL6.18_q89 CBCL6.18_q94 CBCL6.18_q95 CBCL6.18_q97 CBCL6.18_q104
##  0   :901     0   :1230    0   :1218    0   :928     0   :1317    0   :1071    
##  1   :379     1   :  97    1   : 107    1   :318     1   :  10    1   : 202    
##  2   : 55     2   :  11    2   :   7    2   : 85     2   :   1    2   :  61    
##  NA's: 14     NA's:  11    NA's:  17    NA's: 18     NA's:  21    NA's:  15    
##  CBCL6.18_q15 CBCL6.18_q26 CBCL6.18_q39 CBCL6.18_q43 CBCL6.18_q81 CBCL6.18_q90
##  0   :1304    0   :1076    0   :1246    0   :1081    0   :1317    0   :1032   
##  1   :  36    1   : 231    1   :  92    1   : 250    1   :  23    1   : 270   
##  2   :   7    2   :  37    2   :   9    2   :  15    2   :   3    2   :  35   
##  NA's:   2    NA's:   5    NA's:   2    NA's:   3    NA's:   6    NA's:  12   
##  CBCL6.18_q101 CBCL6.18_q106
##  0   :1332     0   :1337    
##  1   :   3     1   :   1    
##  2   :   1     2   :   2    
##  NA's:  13     NA's:   9
```

```
# histogram of missing values per sample
hist(rowSums(is.na(pheno_filt)))
```

```
# histogram of missing values per feature
hist(colSums(is.na(pheno_filt)))
```

```
# write filtered data to output file
write.csv(pheno_filt, params$cbcl_filtered_outfile)
```

#### Impute missing values

Random forests imputation

```
#' Non-parametric missing value imputation in survey data using random forests
#' 
#' @param df Data frame containing subjects in rows and questions in columns 
impute_cbcl_rf <- function(df) {
  library(missForest)
  stopifnot(all(sapply(df, is.factor)))
  # hyperparameters: ntree, mtry, replace, sampsize, nodesize, maxnodes
  # suggestion: could also make use of additional phenotype information 
  # impute values
  df_imp <- missForest(
    df,
    ntree = 1000,
    variablewise = TRUE)
  return(df_imp)
}
```

Imputation using MCA

```
#' Imputing missing categorical data using Multiple Correspondence Analysis
#'  
#' @param df Data frame containing subjects in rows and questions in columns
#' @return list of different plots of MCA
impute_cbcl_mca <- function(df){
  library(missMDA)
  stopifnot(all(sapply(df, is.factor)))
  # estimate the number of dimensions for MCA by cross-validation
  ncp_est <- estim_ncpMCA(
    df, 
    ncp.min = 0, 
    ncp.max = 7,
    method = "Regularized", 
    method.cv = "Kfold",
    verbose = TRUE) 
  print(ncp_est)
  # impute missing values using estimated number of dimensions
  df_imputed <- imputeMCA(df, ncp = ncp_est$ncp)
  return(df_imputed)
}
```

Perform imputation based using chosen method

```
if (params$imputation_method == "RF") {
  pheno_rfimp_res <- impute_cbcl_rf(pheno_filt)
  names(pheno_rfimp_res)
  # out-of-bag (OOB) imputation error estimate
  pheno_rfimp_res$OOBerror
  # extract imputed (complete) data
  pheno_imp <- pheno_rfimp_res$ximp
} else if (params$imputation_method == "MCA") {
  pheno_mcaimp_res <- impute_cbcl_mca(pheno_filt)
  names(pheno_mcaimp_res)
  # extract imputed (complete) data
  pheno_imp <- pheno_mcaimp_res$completeObs
  summary(pheno_imp)
}
```

```
## Loading required package: randomForest
```

```
## randomForest 4.6-14
```

```
## Type rfNews() to see new features/changes/bug fixes.
```

```
## Loading required package: foreach
```

```
## Loading required package: itertools
```

```
## Loading required package: iterators
```

```
##   missForest iteration 1 in progress...done!
##   missForest iteration 2 in progress...done!
##   missForest iteration 3 in progress...done!
##   missForest iteration 4 in progress...done!
##   missForest iteration 5 in progress...done!
```

```
summary(pheno_imp)
```

```
##  CBCL6.18_q3 CBCL6.18_q16 CBCL6.18_q19 CBCL6.18_q20 CBCL6.18_q21 CBCL6.18_q22
##  0:589       0:1130       0:839        0:1201       0:1215       0:738       
##  1:646       1: 210       1:394        1: 134       1: 130       1:563       
##  2:114       2:   9       2:116        2:  14       2:   4       2: 48       
##  CBCL6.18_q23 CBCL6.18_q37 CBCL6.18_q57 CBCL6.18_q68 CBCL6.18_q86 CBCL6.18_q87
##  0:1136       0:1221       0:1274       0:920        0:701        0:989       
##  1: 202       1: 112       1:  68       1:345        1:520        1:286       
##  2:  11       2:  16       2:   7       2: 84        2:128        2: 74       
##  CBCL6.18_q88 CBCL6.18_q89 CBCL6.18_q94 CBCL6.18_q95 CBCL6.18_q97 CBCL6.18_q104
##  0:911        0:1238       0:1234       0:940        0:1338       0:1080       
##  1:383        1: 100       1: 108       1:323        1:  10       1: 207       
##  2: 55        2:  11       2:   7       2: 86        2:   1       2:  62       
##  CBCL6.18_q15 CBCL6.18_q26 CBCL6.18_q39 CBCL6.18_q43 CBCL6.18_q81 CBCL6.18_q90
##  0:1306       0:1080       0:1248       0:1083       0:1323       0:1042      
##  1:  36       1: 232       1:  92       1: 251       1:  23       1: 272      
##  2:   7       2:  37       2:   9       2:  15       2:   3       2:  35      
##  CBCL6.18_q101 CBCL6.18_q106
##  0:1345        0:1346       
##  1:   3        1:   1       
##  2:   1        2:   2
```

```
write.csv(pheno_imp, params$cbcl_imputed_outfile)
```

```
# prepare data for plot
library(reshape2)
library(stringr)
tmp_pheno_list <- list(filtered = pheno_filt, imputed = pheno_imp)
tmp_pheno_list <- lapply(names(tmp_pheno_list), function(x) {
  df <- tmp_pheno_list[[x]]
  df$data <- x
  df
})
pheno_all_wide <- do.call(rbind, tmp_pheno_list)
pheno_all_long <- melt(pheno_all_wide, id.vars = c("data"))
pheno_labels$variable_names <- make.names(pheno_labels$variable)
pheno_all_long$label <- apply(pheno_all_long, 1, function(x) {
  label <- pheno_labels[
    pheno_labels$variable_names == x[["variable"]], ][["label"]]
  str_wrap(gsub("CBCL6_18 - ", "", label), width = 12)
})

rm(tmp_pheno_list)
rm(pheno_all_wide)

# plot distribution of values, before and after imputation
library(ggplot2)
```

```
## 
## Attaching package: 'ggplot2'
```

```
## The following object is masked from 'package:randomForest':
## 
##     margin
```

```
cbf_palette <- c("#000000", "#E69F00", "#56B4E9", "#009E73", "#F0E442", "#0072B2", "#D55E00", "#CC79A7")
lapply(split(names(pheno_filt), ceiling(seq_along(names(pheno_filt)) / 4)), 
       function(x) {
         ggplot(pheno_all_long[pheno_all_long$variable %in% x,]) +
           geom_bar(aes(x = value, fill = value)) +
           scale_fill_manual(values = cbf_palette) +
           facet_grid(label ~ data, scales = "free_y")
})
```

```
## $`1`
```

```
## 
## $`2`
```

```
## 
## $`3`
```

```
## 
## $`4`
```

```
## 
## $`5`
```

```
## 
## $`6`
```

```
## 
## $`7`
```

#### Perform multiple correspondence analysis

Multiple correspondence analysis (MCA) is a dimension reduction technique that can be used for categorical data sets. The results are numerical variables (factors) representing the data.

```
#' Multiple Correspondence Analysis of survey question data with missing values
#'  
#' @param df Data frame containing subjects in rows and questions in columns
#' @return list of different plots of MCA
perform_mca <- function(df){
  library(FactoMineR)
  stopifnot(all(sapply(df, is.factor)))
  mca <- MCA(
    df, 
    ncp = 10, 
    graph = FALSE,
    method = "Indicator" # "Burt" 
  )
  return(mca)
}

names(pheno_imp) <- lapply(names(pheno_imp), function(x) {
  label <- pheno_labels[
    pheno_labels$variable_names == x, ][["label"]]
  str_wrap(gsub("CBCL6_18 - ", "", label), width = 16)
})

mca_pheno <- perform_mca(pheno_imp)
```

##### Quality of dimension reduction

Scree plot - percentage of variation (inertia) explained by dimensions.

```
library(factoextra)
```

```
## Welcome! Want to learn more? See two factoextra-related books at https://goo.gl/ve3WBa
```

```
knitr::kable(
  head(get_eigenvalue(mca_pheno), n = 10),
  caption = "Eigenvalues and explained variance for MCA dimensions 1-10.")
```

Eigenvalues and explained variance for MCA dimensions 1-10.

|  | eigenvalue | variance.percent | cumulative.variance.percent |
| --- | --- | --- | --- |
| Dim.1 | 0.3177127 | 15.885635 | 15.88563 |
| Dim.2 | 0.1708732 | 8.543658 | 24.42929 |
| Dim.3 | 0.1044737 | 5.223683 | 29.65298 |
| Dim.4 | 0.0822893 | 4.114463 | 33.76744 |
| Dim.5 | 0.0679152 | 3.395761 | 37.16320 |
| Dim.6 | 0.0586206 | 2.931030 | 40.09423 |
| Dim.7 | 0.0532005 | 2.660026 | 42.75426 |
| Dim.8 | 0.0500006 | 2.500028 | 45.25428 |
| Dim.9 | 0.0484606 | 2.423028 | 47.67731 |
| Dim.10 | 0.0452584 | 2.262918 | 49.94023 |

```
cat("\n\n")
```

```
fviz_screeplot(mca_pheno, addlabels = TRUE) +
  geom_point()
```

##### Relative contribution of individuals and variables

Biplot of individuals and behavior variable categories

```
fviz_mca_biplot(
  mca_pheno,
  repel = TRUE,
  label = "var",
  ggtheme = theme_minimal())
```

```
## Warning: ggrepel: 68 unlabeled data points (too many overlaps). Consider
## increasing max.overlaps
```

###### Correlation between **variables** and MCA dimensions

```
fviz_mca_var(
  mca_pheno, 
  choice = "mca.cor", 
  axes = c(1, 2),
  repel = TRUE, 
  ggtheme = theme_minimal())
```

```
## Warning: ggrepel: 2 unlabeled data points (too many overlaps). Consider
## increasing max.overlaps
```

###### Coordinates of **variable categories** and contribution to dimensions

```
fviz_mca_var(
  mca_pheno, 
  col.var = "contrib", # color by contribution to dimension
  repel = TRUE, 
  gradient.cols = c("#E69F00", "#56B4E9"),
  ggtheme = theme_minimal())
```

```
## Warning: ggrepel: 68 unlabeled data points (too many overlaps). Consider
## increasing max.overlaps
```

```
lapply(1:10, function(dim) {
  # contribution to MCA dimensions
  fviz_contrib(mca_pheno, choice = "var", axes = dim, top = 10)
})
```

```
## [[1]]
```

```
## 
## [[2]]
```

```
## 
## [[3]]
```

```
## 
## [[4]]
```

```
## 
## [[5]]
```

```
## 
## [[6]]
```

```
## 
## [[7]]
```

```
## 
## [[8]]
```

```
## 
## [[9]]
```

```
## 
## [[10]]
```

###### Degree of association of **variable categories** with dimensions

How well is a variable category represented by the dimensions?

```
fviz_mca_var(
  mca_pheno, 
  col.var = "cos2", # color by quality of representation
  repel = TRUE, 
  gradient.cols = c("#E69F00", "#56B4E9"),
  ggtheme = theme_minimal())
```

```
## Warning: ggrepel: 68 unlabeled data points (too many overlaps). Consider
## increasing max.overlaps
```

```
lapply(1:10, function(dim) {
  # quality of representation (cos2), squared cosine
  fviz_cos2(mca_pheno, choice = "var", axes = dim, top = 10)
})
```

```
## [[1]]
```

```
## 
## [[2]]
```

```
## 
## [[3]]
```

```
## 
## [[4]]
```

```
## 
## [[5]]
```

```
## 
## [[6]]
```

```
## 
## [[7]]
```

```
## 
## [[8]]
```

```
## 
## [[9]]
```

```
## 
## [[10]]
```

MCA dimension description

```
dimdesc_mca <- dimdesc(mca_pheno, axes = 1:10)
dimdesc_mca[paste0("Dim ", 1:10)]
```

```
## $`Dim 1`
## $quali
##                                        R2       p.value
## 95 Temper                      0.55649783 2.295292e-238
## 86 Stubborn,\nsullen           0.52276929 6.084212e-217
## 68 Screams a lot               0.51991786 3.352472e-215
## 3 Argues a lot                 0.47193363 2.340816e-187
## 87 Mood changes                0.46119831 1.785190e-181
## 104 Loud                       0.44597786 2.475998e-173
## 19 Demands a lot\nof attention 0.43070768 2.191170e-165
## 22 Disobedient\nat home        0.42080435 2.406698e-160
## 16 Cruel to\nothers            0.41149787 1.097561e-155
## 88 Sulks                       0.38096676 6.660454e-141
## 94 Teases a lot                0.37393531 1.332694e-137
## 21 Destroys\nothers things     0.36393637 5.700076e-133
## 90 Swearing                    0.35240926 1.012331e-127
## 26 Lacks guilt                 0.33690733 8.307195e-121
## 20 Destroys own\nthings        0.32630579 3.593130e-116
## 43 Lying                       0.30505838 4.277801e-107
## 37 Gets in\nfights             0.28092508  4.064426e-97
## 23 Disobedient\nat school      0.25658240  2.186252e-87
## 89 Suspicious                  0.25083980  3.879453e-85
## 57 Attacks\nothers             0.21268067  1.286958e-70
## 97 Threatens\nothers           0.16674101  4.841119e-54
## 15 Cruel to\nanimals           0.12379255  2.368485e-39
## 106 Vandalism                  0.11442645  3.035149e-36
## 39 Bad friends                 0.09219323  5.365018e-29
## 81 Steals at\nhome             0.04539332  2.641597e-14
## 101 Truant                     0.03613159  1.753651e-11
## 
## $category
##                                                                    Estimate
## 86 Stubborn,\nsullen=86 Stubborn,\nsullen_2                      0.75231375
## 95 Temper=95 Temper_2                                            0.77912180
## 68 Screams a lot=68 Screams a lot_2                              0.78664830
## 3 Argues a lot=3 Argues a lot_2                                  0.75115261
## 87 Mood changes=87 Mood changes_2                                0.77853366
## 19 Demands a lot\nof attention=19 Demands a lot\nof attention_2  0.62875944
## 104 Loud=104 Loud_2                                              0.67352484
## 23 Disobedient\nat school=23 Disobedient\nat school_1            0.07436408
## 88 Sulks=88 Sulks_2                                              0.79481515
## 37 Gets in\nfights=37 Gets in\nfights_1                          0.11189733
## 90 Swearing=90 Swearing_2                                        0.93046691
## 26 Lacks guilt=26 Lacks guilt_1                                  0.01246597
## 104 Loud=104 Loud_1                                              0.03635578
## 57 Attacks\nothers=57 Attacks\nothers_1                          0.27979161
## 22 Disobedient\nat home=22 Disobedient\nat home_2                0.79743713
## 26 Lacks guilt=26 Lacks guilt_2                                  0.66473986
## 16 Cruel to\nothers=16 Cruel to\nothers_2                        1.38910534
## 94 Teases a lot=94 Teases a lot_2                                1.46763647
## 20 Destroys own\nthings=20 Destroys own\nthings_2                0.99142941
## 21 Destroys\nothers things=21 Destroys\nothers things_2          1.97671500
## 106 Vandalism=106 Vandalism_2                                    2.38091659
## 43 Lying=43 Lying_2                                              0.95791039
## 89 Suspicious=89 Suspicious_2                                    0.97881968
## 37 Gets in\nfights=37 Gets in\nfights_2                          0.70118984
## 15 Cruel to\nanimals=15 Cruel to\nanimals_2                      1.13274558
## 97 Threatens\nothers=97 Threatens\nothers_2                      2.54223071
## 23 Disobedient\nat school=23 Disobedient\nat school_2            0.59706767
## 101 Truant=101 Truant_1                                          1.11400537
## 81 Steals at\nhome=81 Steals at\nhome_1                          0.06458332
## 39 Bad friends=39 Bad friends_2                                  0.62262181
## 57 Attacks\nothers=57 Attacks\nothers_2                          0.54707584
## 81 Steals at\nhome=81 Steals at\nhome_2                          0.65515106
## 106 Vandalism=106 Vandalism_1                                   -0.06658970
## 101 Truant=101 Truant_0                                         -1.07964417
## 81 Steals at\nhome=81 Steals at\nhome_0                         -0.71973437
## 86 Stubborn,\nsullen=86 Stubborn,\nsullen_1                     -0.13896449
## 15 Cruel to\nanimals=15 Cruel to\nanimals_1                     -0.16226827
## 3 Argues a lot=3 Argues a lot_1                                 -0.13321004
## 39 Bad friends=39 Bad friends_1                                 -0.02993251
## 39 Bad friends=39 Bad friends_0                                 -0.59268931
## 19 Demands a lot\nof attention=19 Demands a lot\nof attention_1 -0.05738701
## 15 Cruel to\nanimals=15 Cruel to\nanimals_0                     -0.97047731
## 106 Vandalism=106 Vandalism_0                                   -2.31432689
## 97 Threatens\nothers=97 Threatens\nothers_1                     -0.17669365
## 87 Mood changes=87 Mood changes_1                               -0.09967604
## 88 Sulks=88 Sulks_1                                             -0.13508097
## 68 Screams a lot=68 Screams a lot_1                             -0.09908817
## 90 Swearing=90 Swearing_1                                       -0.18439887
## 95 Temper=95 Temper_1                                           -0.07139217
## 97 Threatens\nothers=97 Threatens\nothers_0                     -2.36553706
## 89 Suspicious=89 Suspicious_1                                   -0.05270133
## 22 Disobedient\nat home=22 Disobedient\nat home_1               -0.10569899
## 43 Lying=43 Lying_1                                             -0.14552917
## 20 Destroys own\nthings=20 Destroys own\nthings_1               -0.05646145
## 57 Attacks\nothers=57 Attacks\nothers_0                         -0.82686745
## 89 Suspicious=89 Suspicious_0                                   -0.92611835
## 23 Disobedient\nat school=23 Disobedient\nat school_0           -0.67143175
## 94 Teases a lot=94 Teases a lot_1                               -0.20398826
## 21 Destroys\nothers things=21 Destroys\nothers things_1         -0.50218402
## 90 Swearing=90 Swearing_0                                       -0.74606805
## 43 Lying=43 Lying_0                                             -0.81238122
## 37 Gets in\nfights=37 Gets in\nfights_0                         -0.81308717
## 16 Cruel to\nothers=16 Cruel to\nothers_1                       -0.27277623
## 88 Sulks=88 Sulks_0                                             -0.65973418
## 20 Destroys own\nthings=20 Destroys own\nthings_0               -0.93496796
## 3 Argues a lot=3 Argues a lot_0                                 -0.61794257
## 26 Lacks guilt=26 Lacks guilt_0                                 -0.67720583
## 21 Destroys\nothers things=21 Destroys\nothers things_0         -1.47453099
## 94 Teases a lot=94 Teases a lot_0                               -1.26364821
## 19 Demands a lot\nof attention=19 Demands a lot\nof attention_0 -0.57137243
## 86 Stubborn,\nsullen=86 Stubborn,\nsullen_0                     -0.61334926
## 22 Disobedient\nat home=22 Disobedient\nat home_0               -0.69173814
## 87 Mood changes=87 Mood changes_0                               -0.67885762
## 16 Cruel to\nothers=16 Cruel to\nothers_0                       -1.11632911
## 68 Screams a lot=68 Screams a lot_0                             -0.68756013
## 104 Loud=104 Loud_0                                             -0.70988062
## 95 Temper=95 Temper_0                                           -0.70772963
##                                                                       p.value
## 86 Stubborn,\nsullen=86 Stubborn,\nsullen_2                     1.799995e-135
## 95 Temper=95 Temper_2                                           5.135356e-119
## 68 Screams a lot=68 Screams a lot_2                             9.736245e-114
## 3 Argues a lot=3 Argues a lot_2                                 1.000762e-107
## 87 Mood changes=87 Mood changes_2                               2.699477e-101
## 19 Demands a lot\nof attention=19 Demands a lot\nof attention_2  2.443133e-92
## 104 Loud=104 Loud_2                                              7.658956e-75
## 23 Disobedient\nat school=23 Disobedient\nat school_1            4.074611e-73
## 88 Sulks=88 Sulks_2                                              3.506191e-70
## 37 Gets in\nfights=37 Gets in\nfights_1                          5.280116e-66
## 90 Swearing=90 Swearing_2                                        5.472900e-65
## 26 Lacks guilt=26 Lacks guilt_1                                  1.959391e-62
## 104 Loud=104 Loud_1                                              1.478001e-61
## 57 Attacks\nothers=57 Attacks\nothers_1                          8.681261e-61
## 22 Disobedient\nat home=22 Disobedient\nat home_2                1.050129e-54
## 26 Lacks guilt=26 Lacks guilt_2                                  5.501151e-41
## 16 Cruel to\nothers=16 Cruel to\nothers_2                        1.756676e-38
## 94 Teases a lot=94 Teases a lot_2                                3.047213e-37
## 20 Destroys own\nthings=20 Destroys own\nthings_2                9.600764e-36
## 21 Destroys\nothers things=21 Destroys\nothers things_2          2.605999e-34
## 106 Vandalism=106 Vandalism_2                                    1.410914e-33
## 43 Lying=43 Lying_2                                              1.238709e-30
## 89 Suspicious=89 Suspicious_2                                    3.271862e-28
## 37 Gets in\nfights=37 Gets in\nfights_2                          5.400836e-25
## 15 Cruel to\nanimals=15 Cruel to\nanimals_2                      3.607390e-23
## 97 Threatens\nothers=97 Threatens\nothers_2                      1.474817e-18
## 23 Disobedient\nat school=23 Disobedient\nat school_2            8.730659e-12
## 101 Truant=101 Truant_1                                          1.183766e-11
## 81 Steals at\nhome=81 Steals at\nhome_1                          3.185690e-11
## 39 Bad friends=39 Bad friends_2                                  3.348643e-10
## 57 Attacks\nothers=57 Attacks\nothers_2                          5.356498e-10
## 81 Steals at\nhome=81 Steals at\nhome_2                          2.805154e-05
## 106 Vandalism=106 Vandalism_1                                    6.835768e-05
## 101 Truant=101 Truant_0                                          9.982009e-12
## 81 Steals at\nhome=81 Steals at\nhome_0                          1.216604e-14
## 86 Stubborn,\nsullen=86 Stubborn,\nsullen_1                      2.652739e-17
## 15 Cruel to\nanimals=15 Cruel to\nanimals_1                      2.375095e-17
## 3 Argues a lot=3 Argues a lot_1                                  4.585544e-18
## 39 Bad friends=39 Bad friends_1                                  2.534207e-20
## 39 Bad friends=39 Bad friends_0                                  1.558479e-27
## 19 Demands a lot\nof attention=19 Demands a lot\nof attention_1  6.962456e-29
## 15 Cruel to\nanimals=15 Cruel to\nanimals_0                      5.431001e-33
## 106 Vandalism=106 Vandalism_0                                    2.192481e-34
## 97 Threatens\nothers=97 Threatens\nothers_1                      3.424969e-36
## 87 Mood changes=87 Mood changes_1                                2.527440e-39
## 88 Sulks=88 Sulks_1                                              5.686162e-41
## 68 Screams a lot=68 Screams a lot_1                              1.497025e-42
## 90 Swearing=90 Swearing_1                                        8.534613e-43
## 95 Temper=95 Temper_1                                            1.519332e-49
## 97 Threatens\nothers=97 Threatens\nothers_0                      6.986037e-50
## 89 Suspicious=89 Suspicious_1                                    1.779528e-52
## 22 Disobedient\nat home=22 Disobedient\nat home_1                8.433702e-63
## 43 Lying=43 Lying_1                                              3.632614e-66
## 20 Destroys own\nthings=20 Destroys own\nthings_1                1.070748e-69
## 57 Attacks\nothers=57 Attacks\nothers_0                          1.329234e-71
## 89 Suspicious=89 Suspicious_0                                    4.326613e-77
## 23 Disobedient\nat school=23 Disobedient\nat school_0            3.292080e-86
## 94 Teases a lot=94 Teases a lot_1                                2.517249e-87
## 21 Destroys\nothers things=21 Destroys\nothers things_1          1.829741e-87
## 90 Swearing=90 Swearing_0                                        3.733102e-91
## 43 Lying=43 Lying_0                                              4.262231e-92
## 37 Gets in\nfights=37 Gets in\nfights_0                          5.451007e-94
## 16 Cruel to\nothers=16 Cruel to\nothers_1                        1.701640e-98
## 88 Sulks=88 Sulks_0                                             8.314739e-100
## 20 Destroys own\nthings=20 Destroys own\nthings_0               6.960676e-104
## 3 Argues a lot=3 Argues a lot_0                                 2.045335e-104
## 26 Lacks guilt=26 Lacks guilt_0                                 1.186194e-108
## 21 Destroys\nothers things=21 Destroys\nothers things_0         5.986670e-110
## 94 Teases a lot=94 Teases a lot_0                               9.295033e-120
## 19 Demands a lot\nof attention=19 Demands a lot\nof attention_0 2.789072e-120
## 86 Stubborn,\nsullen=86 Stubborn,\nsullen_0                     2.463704e-120
## 22 Disobedient\nat home=22 Disobedient\nat home_0               3.346099e-122
## 87 Mood changes=87 Mood changes_0                               1.301793e-130
## 16 Cruel to\nothers=16 Cruel to\nothers_0                       7.898867e-131
## 68 Screams a lot=68 Screams a lot_0                             8.662408e-150
## 104 Loud=104 Loud_0                                             5.461495e-152
## 95 Temper=95 Temper_0                                           8.728569e-173
## 
## attr(,"class")
## [1] "condes" "list"  
## 
## $`Dim 2`
## $quali
##                                        R2       p.value
## 106 Vandalism                  0.51240693 1.155349e-210
## 21 Destroys\nothers things     0.45702823 3.199560e-179
## 94 Teases a lot                0.40924033 1.443758e-154
## 97 Threatens\nothers           0.38208851 1.965070e-141
## 16 Cruel to\nothers            0.37850914 9.584801e-140
## 20 Destroys own\nthings        0.21148310  3.579379e-70
## 101 Truant                     0.18715916  2.714642e-61
## 15 Cruel to\nanimals           0.16689883  4.261695e-54
## 86 Stubborn,\nsullen           0.14886311  7.755455e-48
## 90 Swearing                    0.14692912  3.572596e-47
## 87 Mood changes                0.14220852  1.465223e-45
## 68 Screams a lot               0.14046593  5.741996e-45
## 22 Disobedient\nat home        0.13988977  9.013991e-45
## 95 Temper                      0.13880505  2.105198e-44
## 3 Argues a lot                 0.12430660  1.595689e-39
## 104 Loud                       0.11667259  5.494263e-37
## 88 Sulks                       0.11011511  7.973898e-35
## 26 Lacks guilt                 0.10889341  2.007511e-34
## 43 Lying                       0.10600158  1.776783e-33
## 19 Demands a lot\nof attention 0.09364773  1.823473e-29
## 89 Suspicious                  0.08769796  1.490417e-27
## 39 Bad friends                 0.07312240  6.398327e-23
## 23 Disobedient\nat school      0.02673435  1.201513e-08
## 37 Gets in\nfights             0.01634339  1.526461e-05
## 57 Attacks\nothers             0.01366826  9.494783e-05
## 
## $category
##                                                                     Estimate
## 106 Vandalism=106 Vandalism_2                                    4.467197749
## 21 Destroys\nothers things=21 Destroys\nothers things_2          3.362421269
## 94 Teases a lot=94 Teases a lot_2                                2.431617568
## 97 Threatens\nothers=97 Threatens\nothers_2                      5.873792376
## 16 Cruel to\nothers=16 Cruel to\nothers_2                        1.937002034
## 101 Truant=101 Truant_1                                          2.859831581
## 20 Destroys own\nthings=20 Destroys own\nthings_2                1.181980851
## 15 Cruel to\nanimals=15 Cruel to\nanimals_2                      1.580242072
## 22 Disobedient\nat home=22 Disobedient\nat home_0                0.114279055
## 86 Stubborn,\nsullen=86 Stubborn,\nsullen_0                      0.105355499
## 3 Argues a lot=3 Argues a lot_0                                  0.096942630
## 89 Suspicious=89 Suspicious_2                                    0.851107573
## 95 Temper=95 Temper_0                                            0.051684124
## 90 Swearing=90 Swearing_2                                        0.482087636
## 68 Screams a lot=68 Screams a lot_0                              0.037054597
## 87 Mood changes=87 Mood changes_2                                0.345442415
## 19 Demands a lot\nof attention=19 Demands a lot\nof attention_0  0.078524592
## 104 Loud=104 Loud_2                                              0.356183649
## 43 Lying=43 Lying_2                                              0.612809597
## 26 Lacks guilt=26 Lacks guilt_2                                  0.408061599
## 88 Sulks=88 Sulks_2                                              0.310780205
## 39 Bad friends=39 Bad friends_2                                  0.686541410
## 68 Screams a lot=68 Screams a lot_2                              0.244883705
## 23 Disobedient\nat school=23 Disobedient\nat school_0            0.049295417
## 95 Temper=95 Temper_2                                            0.228525457
## 37 Gets in\nfights=37 Gets in\nfights_0                          0.149619956
## 57 Attacks\nothers=57 Attacks\nothers_0                          0.118024684
## 86 Stubborn,\nsullen=86 Stubborn,\nsullen_2                      0.115379541
## 3 Argues a lot=3 Argues a lot_2                                  0.097730714
## 19 Demands a lot\nof attention=19 Demands a lot\nof attention_2  0.115717269
## 81 Steals at\nhome=81 Steals at\nhome_2                          0.349301407
## 89 Suspicious=89 Suspicious_0                                   -0.308084193
## 101 Truant=101 Truant_2                                         -1.977110971
## 37 Gets in\nfights=37 Gets in\nfights_2                         -0.141765147
## 21 Destroys\nothers things=21 Destroys\nothers things_0         -1.549740792
## 20 Destroys own\nthings=20 Destroys own\nthings_0               -0.440634846
## 37 Gets in\nfights=37 Gets in\nfights_1                         -0.007854809
## 57 Attacks\nothers=57 Attacks\nothers_1                         -0.098980918
## 106 Vandalism=106 Vandalism_1                                   -1.345784607
## 97 Threatens\nothers=97 Threatens\nothers_1                     -2.599035234
## 104 Loud=104 Loud_0                                             -0.026977343
## 39 Bad friends=39 Bad friends_0                                 -0.173490199
## 15 Cruel to\nanimals=15 Cruel to\nanimals_1                     -0.980000796
## 89 Suspicious=89 Suspicious_1                                   -0.543023380
## 23 Disobedient\nat school=23 Disobedient\nat school_1           -0.139453136
## 16 Cruel to\nothers=16 Cruel to\nothers_0                       -0.812492242
## 87 Mood changes=87 Mood changes_0                               -0.024663356
## 26 Lacks guilt=26 Lacks guilt_0                                 -0.062262442
## 94 Teases a lot=94 Teases a lot_1                               -1.352541034
## 88 Sulks=88 Sulks_0                                             -0.032650274
## 43 Lying=43 Lying_0                                             -0.175192773
## 21 Destroys\nothers things=21 Destroys\nothers things_1         -1.812680477
## 90 Swearing=90 Swearing_0                                       -0.088082658
## 39 Bad friends=39 Bad friends_1                                 -0.513051211
## 20 Destroys own\nthings=20 Destroys own\nthings_1               -0.741346005
## 43 Lying=43 Lying_1                                             -0.437616824
## 26 Lacks guilt=26 Lacks guilt_1                                 -0.345799157
## 104 Loud=104 Loud_1                                             -0.329206306
## 88 Sulks=88 Sulks_1                                             -0.278129931
## 16 Cruel to\nothers=16 Cruel to\nothers_1                       -1.124509792
## 19 Demands a lot\nof attention=19 Demands a lot\nof attention_1 -0.194241861
## 97 Threatens\nothers=97 Threatens\nothers_0                     -3.274757142
## 90 Swearing=90 Swearing_1                                       -0.394004978
## 87 Mood changes=87 Mood changes_1                               -0.320779059
## 101 Truant=101 Truant_0                                         -0.882720610
## 3 Argues a lot=3 Argues a lot_1                                 -0.194673344
## 68 Screams a lot=68 Screams a lot_1                             -0.281938302
## 95 Temper=95 Temper_1                                           -0.280209581
## 22 Disobedient\nat home=22 Disobedient\nat home_1               -0.200776421
## 86 Stubborn,\nsullen=86 Stubborn,\nsullen_1                     -0.220735041
## 106 Vandalism=106 Vandalism_0                                   -3.121413142
##                                                                       p.value
## 106 Vandalism=106 Vandalism_2                                   3.025777e-204
## 21 Destroys\nothers things=21 Destroys\nothers things_2         1.909953e-162
## 94 Teases a lot=94 Teases a lot_2                               1.336023e-140
## 97 Threatens\nothers=97 Threatens\nothers_2                     8.088885e-134
## 16 Cruel to\nothers=16 Cruel to\nothers_2                       5.153529e-108
## 101 Truant=101 Truant_1                                          9.054033e-61
## 20 Destroys own\nthings=20 Destroys own\nthings_2                1.847721e-54
## 15 Cruel to\nanimals=15 Cruel to\nanimals_2                      8.725515e-48
## 22 Disobedient\nat home=22 Disobedient\nat home_0                1.112522e-40
## 86 Stubborn,\nsullen=86 Stubborn,\nsullen_0                      3.162005e-32
## 3 Argues a lot=3 Argues a lot_0                                  6.315537e-29
## 89 Suspicious=89 Suspicious_2                                    1.248652e-21
## 95 Temper=95 Temper_0                                            1.100987e-20
## 90 Swearing=90 Swearing_2                                        1.116496e-19
## 68 Screams a lot=68 Screams a lot_0                              1.313804e-19
## 87 Mood changes=87 Mood changes_2                                3.392865e-19
## 19 Demands a lot\nof attention=19 Demands a lot\nof attention_0  1.019787e-18
## 104 Loud=104 Loud_2                                              4.493854e-16
## 43 Lying=43 Lying_2                                              3.148857e-15
## 26 Lacks guilt=26 Lacks guilt_2                                  2.392440e-14
## 88 Sulks=88 Sulks_2                                              1.601832e-13
## 39 Bad friends=39 Bad friends_2                                  1.254417e-10
## 68 Screams a lot=68 Screams a lot_2                              1.866357e-10
## 23 Disobedient\nat school=23 Disobedient\nat school_0            8.354196e-09
## 95 Temper=95 Temper_2                                            1.127135e-08
## 37 Gets in\nfights=37 Gets in\nfights_0                          5.376722e-06
## 57 Attacks\nothers=57 Attacks\nothers_0                          1.895913e-05
## 86 Stubborn,\nsullen=86 Stubborn,\nsullen_2                      1.025739e-04
## 3 Argues a lot=3 Argues a lot_2                                  1.471025e-04
## 19 Demands a lot\nof attention=19 Demands a lot\nof attention_2  1.933447e-03
## 81 Steals at\nhome=81 Steals at\nhome_2                          3.791075e-02
## 89 Suspicious=89 Suspicious_0                                    1.811236e-02
## 101 Truant=101 Truant_2                                          7.637109e-03
## 37 Gets in\nfights=37 Gets in\nfights_2                          7.436165e-03
## 21 Destroys\nothers things=21 Destroys\nothers things_0          3.923623e-03
## 20 Destroys own\nthings=20 Destroys own\nthings_0                9.604334e-04
## 37 Gets in\nfights=37 Gets in\nfights_1                          1.601775e-04
## 57 Attacks\nothers=57 Attacks\nothers_1                          2.510115e-05
## 106 Vandalism=106 Vandalism_1                                    1.892350e-05
## 97 Threatens\nothers=97 Threatens\nothers_1                      3.086455e-07
## 104 Loud=104 Loud_0                                              2.721941e-07
## 39 Bad friends=39 Bad friends_0                                  4.605832e-08
## 15 Cruel to\nanimals=15 Cruel to\nanimals_1                      1.777048e-08
## 89 Suspicious=89 Suspicious_1                                    9.691623e-09
## 23 Disobedient\nat school=23 Disobedient\nat school_1            1.624771e-09
## 16 Cruel to\nothers=16 Cruel to\nothers_0                        8.353645e-10
## 87 Mood changes=87 Mood changes_0                                3.057330e-10
## 26 Lacks guilt=26 Lacks guilt_0                                  1.294183e-10
## 94 Teases a lot=94 Teases a lot_1                                9.900856e-13
## 88 Sulks=88 Sulks_0                                              6.077909e-13
## 43 Lying=43 Lying_0                                              4.275174e-13
## 21 Destroys\nothers things=21 Destroys\nothers things_1          1.533423e-13
## 90 Swearing=90 Swearing_0                                        8.881493e-15
## 39 Bad friends=39 Bad friends_1                                  5.096449e-15
## 20 Destroys own\nthings=20 Destroys own\nthings_1                8.144639e-18
## 43 Lying=43 Lying_1                                              8.041090e-22
## 26 Lacks guilt=26 Lacks guilt_1                                  1.704758e-24
## 104 Loud=104 Loud_1                                              5.110157e-26
## 88 Sulks=88 Sulks_1                                              2.212193e-27
## 16 Cruel to\nothers=16 Cruel to\nothers_1                        4.651791e-28
## 19 Demands a lot\nof attention=19 Demands a lot\nof attention_1  1.934695e-30
## 97 Threatens\nothers=97 Threatens\nothers_0                      2.267534e-32
## 90 Swearing=90 Swearing_1                                        1.939095e-32
## 87 Mood changes=87 Mood changes_1                                3.913239e-33
## 101 Truant=101 Truant_0                                          2.428985e-36
## 3 Argues a lot=3 Argues a lot_1                                  9.160102e-41
## 68 Screams a lot=68 Screams a lot_1                              2.389719e-41
## 95 Temper=95 Temper_1                                            4.789935e-42
## 22 Disobedient\nat home=22 Disobedient\nat home_1                5.445847e-46
## 86 Stubborn,\nsullen=86 Stubborn,\nsullen_1                      4.167249e-49
## 106 Vandalism=106 Vandalism_0                                   7.379885e-159
## 
## attr(,"class")
## [1] "condes" "list"  
## 
## $`Dim 3`
## $quali
##                                         R2      p.value
## 86 Stubborn,\nsullen           0.287179893 1.136321e-99
## 95 Temper                      0.243817593 2.068621e-82
## 87 Mood changes                0.217082962 2.955941e-72
## 68 Screams a lot               0.216412624 5.258247e-72
## 97 Threatens\nothers           0.171214491 1.292904e-55
## 22 Disobedient\nat home        0.166881350 4.322295e-54
## 15 Cruel to\nanimals           0.164849794 2.226234e-53
## 106 Vandalism                  0.151844759 7.309884e-49
## 88 Sulks                       0.144301961 2.829575e-46
## 3 Argues a lot                 0.115377624 1.472585e-36
## 21 Destroys\nothers things     0.104414526 5.861900e-33
## 101 Truant                     0.093884137 1.529863e-29
## 23 Disobedient\nat school      0.089238540 4.778876e-28
## 37 Gets in\nfights             0.076648476 4.920829e-24
## 19 Demands a lot\nof attention 0.071082217 2.810008e-22
## 104 Loud                       0.070888868 3.232492e-22
## 94 Teases a lot                0.067239776 4.520870e-21
## 90 Swearing                    0.066201718 9.556888e-21
## 57 Attacks\nothers             0.046775834 9.960119e-15
## 16 Cruel to\nothers            0.033624708 1.007245e-10
## 20 Destroys own\nthings        0.028395216 3.806542e-09
## 81 Steals at\nhome             0.025878226 2.171290e-08
## 39 Bad friends                 0.024297897 6.464029e-08
## 89 Suspicious                  0.022635236 2.033032e-07
## 43 Lying                       0.011013871 5.794352e-04
## 26 Lacks guilt                 0.005132942 3.132414e-02
## 
## $category
##                                                                     Estimate
## 86 Stubborn,\nsullen=86 Stubborn,\nsullen_1                      0.275667363
## 15 Cruel to\nanimals=15 Cruel to\nanimals_2                      1.221601650
## 106 Vandalism=106 Vandalism_2                                    2.050086488
## 97 Threatens\nothers=97 Threatens\nothers_2                      3.175059465
## 22 Disobedient\nat home=22 Disobedient\nat home_1                0.269888490
## 95 Temper=95 Temper_1                                            0.302044563
## 21 Destroys\nothers things=21 Destroys\nothers things_2          1.282526871
## 68 Screams a lot=68 Screams a lot_1                              0.284790768
## 3 Argues a lot=3 Argues a lot_1                                  0.171067196
## 101 Truant=101 Truant_1                                          0.956174686
## 87 Mood changes=87 Mood changes_1                                0.291010091
## 94 Teases a lot=94 Teases a lot_2                                0.723255312
## 22 Disobedient\nat home=22 Disobedient\nat home_0                0.082936543
## 23 Disobedient\nat school=23 Disobedient\nat school_1            0.376090784
## 19 Demands a lot\nof attention=19 Demands a lot\nof attention_1  0.148246022
## 88 Sulks=88 Sulks_1                                              0.253761054
## 86 Stubborn,\nsullen=86 Stubborn,\nsullen_0                      0.029757134
## 90 Swearing=90 Swearing_1                                        0.209789598
## 37 Gets in\nfights=37 Gets in\nfights_1                          0.343851214
## 16 Cruel to\nothers=16 Cruel to\nothers_2                        0.486812577
## 20 Destroys own\nthings=20 Destroys own\nthings_2                0.349853941
## 81 Steals at\nhome=81 Steals at\nhome_1                          0.160860995
## 23 Disobedient\nat school=23 Disobedient\nat school_0            0.192255929
## 104 Loud=104 Loud_1                                              0.186689949
## 89 Suspicious=89 Suspicious_0                                    0.179230021
## 90 Swearing=90 Swearing_0                                        0.057526863
## 39 Bad friends=39 Bad friends_2                                  0.240284841
## 37 Gets in\nfights=37 Gets in\nfights_0                          0.130388023
## 68 Screams a lot=68 Screams a lot_0                              0.086401667
## 43 Lying=43 Lying_1                                              0.023403162
## 89 Suspicious=89 Suspicious_1                                    0.056509202
## 101 Truant=101 Truant_2                                          0.086983953
## 57 Attacks\nothers=57 Attacks\nothers_0                          0.335431888
## 95 Temper=95 Temper_0                                            0.097593108
## 19 Demands a lot\nof attention=19 Demands a lot\nof attention_0  0.022602614
## 26 Lacks guilt=26 Lacks guilt_1                                  0.004686854
## 97 Threatens\nothers=97 Threatens\nothers_0                     -1.250226097
## 94 Teases a lot=94 Teases a lot_0                               -0.287277840
## 20 Destroys own\nthings=20 Destroys own\nthings_1               -0.209554912
## 26 Lacks guilt=26 Lacks guilt_0                                 -0.046398477
## 43 Lying=43 Lying_0                                             -0.060976719
## 39 Bad friends=39 Bad friends_1                                 -0.047225147
## 89 Suspicious=89 Suspicious_2                                   -0.235739222
## 94 Teases a lot=94 Teases a lot_1                               -0.435977473
## 39 Bad friends=39 Bad friends_0                                 -0.193059694
## 15 Cruel to\nanimals=15 Cruel to\nanimals_0                     -0.604389357
## 81 Steals at\nhome=81 Steals at\nhome_0                         -0.227024875
## 90 Swearing=90 Swearing_2                                       -0.267316461
## 97 Threatens\nothers=97 Threatens\nothers_1                     -1.924833368
## 3 Argues a lot=3 Argues a lot_0                                 -0.005680104
## 19 Demands a lot\nof attention=19 Demands a lot\nof attention_2 -0.170848636
## 37 Gets in\nfights=37 Gets in\nfights_2                         -0.474239237
## 57 Attacks\nothers=57 Attacks\nothers_2                         -0.632424386
## 3 Argues a lot=3 Argues a lot_2                                 -0.165387091
## 23 Disobedient\nat school=23 Disobedient\nat school_2           -0.568346714
## 104 Loud=104 Loud_2                                             -0.270874870
## 22 Disobedient\nat home=22 Disobedient\nat home_2               -0.352825032
## 101 Truant=101 Truant_0                                         -1.043158638
## 106 Vandalism=106 Vandalism_0                                   -1.212915477
## 88 Sulks=88 Sulks_2                                             -0.390207090
## 68 Screams a lot=68 Screams a lot_2                             -0.371192435
## 86 Stubborn,\nsullen=86 Stubborn,\nsullen_2                     -0.305424497
## 95 Temper=95 Temper_2                                           -0.399637671
## 87 Mood changes=87 Mood changes_2                               -0.430351674
##                                                                      p.value
## 86 Stubborn,\nsullen=86 Stubborn,\nsullen_1                     1.220894e-67
## 15 Cruel to\nanimals=15 Cruel to\nanimals_2                     1.121598e-54
## 106 Vandalism=106 Vandalism_2                                   8.303829e-50
## 97 Threatens\nothers=97 Threatens\nothers_2                     8.564842e-46
## 22 Disobedient\nat home=22 Disobedient\nat home_1               9.898673e-35
## 95 Temper=95 Temper_1                                           1.515882e-34
## 21 Destroys\nothers things=21 Destroys\nothers things_2         1.245685e-33
## 68 Screams a lot=68 Screams a lot_1                             2.210859e-33
## 3 Argues a lot=3 Argues a lot_1                                 4.368478e-32
## 101 Truant=101 Truant_1                                         8.739024e-28
## 87 Mood changes=87 Mood changes_1                               2.019574e-19
## 94 Teases a lot=94 Teases a lot_2                               2.887163e-17
## 22 Disobedient\nat home=22 Disobedient\nat home_0               3.004282e-15
## 23 Disobedient\nat school=23 Disobedient\nat school_1           4.853241e-15
## 19 Demands a lot\nof attention=19 Demands a lot\nof attention_1 6.925576e-15
## 88 Sulks=88 Sulks_1                                             2.505134e-14
## 86 Stubborn,\nsullen=86 Stubborn,\nsullen_0                     5.664623e-14
## 90 Swearing=90 Swearing_1                                       6.693588e-14
## 37 Gets in\nfights=37 Gets in\nfights_1                         2.659668e-12
## 16 Cruel to\nothers=16 Cruel to\nothers_2                       1.300577e-11
## 20 Destroys own\nthings=20 Destroys own\nthings_2               8.691650e-09
## 81 Steals at\nhome=81 Steals at\nhome_1                         1.030581e-08
## 23 Disobedient\nat school=23 Disobedient\nat school_0           1.866239e-08
## 104 Loud=104 Loud_1                                             5.552532e-07
## 89 Suspicious=89 Suspicious_0                                   2.015603e-06
## 90 Swearing=90 Swearing_0                                       2.906008e-06
## 39 Bad friends=39 Bad friends_2                                 8.724179e-05
## 37 Gets in\nfights=37 Gets in\nfights_0                         2.079783e-04
## 68 Screams a lot=68 Screams a lot_0                             2.093695e-04
## 43 Lying=43 Lying_1                                             2.354014e-04
## 89 Suspicious=89 Suspicious_1                                   3.858983e-04
## 101 Truant=101 Truant_2                                         4.902333e-04
## 57 Attacks\nothers=57 Attacks\nothers_0                         1.102504e-03
## 95 Temper=95 Temper_0                                           2.934858e-03
## 19 Demands a lot\nof attention=19 Demands a lot\nof attention_0 3.435158e-03
## 26 Lacks guilt=26 Lacks guilt_1                                 3.890979e-02
## 97 Threatens\nothers=97 Threatens\nothers_0                     3.109111e-02
## 94 Teases a lot=94 Teases a lot_0                               1.313851e-02
## 20 Destroys own\nthings=20 Destroys own\nthings_1               1.087893e-02
## 26 Lacks guilt=26 Lacks guilt_0                                 1.073129e-02
## 43 Lying=43 Lying_0                                             1.142679e-04
## 39 Bad friends=39 Bad friends_1                                 4.175716e-05
## 89 Suspicious=89 Suspicious_2                                   3.233611e-05
## 94 Teases a lot=94 Teases a lot_1                               1.777899e-06
## 39 Bad friends=39 Bad friends_0                                 2.632935e-07
## 15 Cruel to\nanimals=15 Cruel to\nanimals_0                     8.967104e-09
## 81 Steals at\nhome=81 Steals at\nhome_0                         3.179871e-09
## 90 Swearing=90 Swearing_2                                       9.068613e-11
## 97 Threatens\nothers=97 Threatens\nothers_1                     2.813520e-11
## 3 Argues a lot=3 Argues a lot_0                                 7.054657e-13
## 19 Demands a lot\nof attention=19 Demands a lot\nof attention_2 5.783646e-14
## 37 Gets in\nfights=37 Gets in\nfights_2                         1.020438e-14
## 57 Attacks\nothers=57 Attacks\nothers_2                         1.564481e-15
## 3 Argues a lot=3 Argues a lot_2                                 7.711249e-16
## 23 Disobedient\nat school=23 Disobedient\nat school_2           3.690837e-16
## 104 Loud=104 Loud_2                                             3.111943e-19
## 22 Disobedient\nat home=22 Disobedient\nat home_2               1.050373e-28
## 101 Truant=101 Truant_0                                         2.067115e-29
## 106 Vandalism=106 Vandalism_0                                   8.506906e-37
## 88 Sulks=88 Sulks_2                                             1.207024e-38
## 68 Screams a lot=68 Screams a lot_2                             2.966179e-48
## 86 Stubborn,\nsullen=86 Stubborn,\nsullen_2                     1.137809e-52
## 95 Temper=95 Temper_2                                           2.126713e-57
## 87 Mood changes=87 Mood changes_2                               2.233511e-60
## 
## attr(,"class")
## [1] "condes" "list"  
## 
## $`Dim 4`
## $quali
##                                         R2       p.value
## 106 Vandalism                  0.662530038 3.178611e-318
## 81 Steals at\nhome             0.556609001 1.938944e-238
## 39 Bad friends                 0.215188918  1.502901e-71
## 43 Lying                       0.202846021  5.469063e-67
## 94 Teases a lot                0.100794889  8.850135e-32
## 26 Lacks guilt                 0.050474328  7.278077e-16
## 90 Swearing                    0.048975370  2.104057e-15
## 89 Suspicious                  0.046384194  1.313185e-14
## 21 Destroys\nothers things     0.045203879  3.019012e-14
## 20 Destroys own\nthings        0.035645411  2.462260e-11
## 97 Threatens\nothers           0.028616995  3.264427e-09
## 88 Sulks                       0.025197750  3.473922e-08
## 101 Truant                     0.022676834  1.975622e-07
## 37 Gets in\nfights             0.021234386  5.330491e-07
## 15 Cruel to\nanimals           0.020942944  6.513060e-07
## 68 Screams a lot               0.019582348  1.658384e-06
## 16 Cruel to\nothers            0.009325953  1.825656e-03
## 57 Attacks\nothers             0.006396909  1.331388e-02
## 19 Demands a lot\nof attention 0.006060676  1.671839e-02
## 104 Loud                       0.005628481  2.240059e-02
## 
## $category
##                                                                     Estimate
## 106 Vandalism=106 Vandalism_1                                    5.959244893
## 81 Steals at\nhome=81 Steals at\nhome_2                          3.016475612
## 39 Bad friends=39 Bad friends_2                                  1.088879448
## 43 Lying=43 Lying_2                                              0.818866783
## 94 Teases a lot=94 Teases a lot_2                                0.844624345
## 90 Swearing=90 Swearing_2                                        0.267732400
## 26 Lacks guilt=26 Lacks guilt_2                                  0.261304044
## 21 Destroys\nothers things=21 Destroys\nothers things_1          0.350679085
## 101 Truant=101 Truant_0                                          0.410106265
## 21 Destroys\nothers things=21 Destroys\nothers things_0          0.199317080
## 37 Gets in\nfights=37 Gets in\nfights_0                          0.018882173
## 15 Cruel to\nanimals=15 Cruel to\nanimals_0                      0.210938526
## 68 Screams a lot=68 Screams a lot_1                              0.075331855
## 16 Cruel to\nothers=16 Cruel to\nothers_0                        0.090898368
## 89 Suspicious=89 Suspicious_1                                    0.267120977
## 97 Threatens\nothers=97 Threatens\nothers_0                      0.618702541
## 19 Demands a lot\nof attention=19 Demands a lot\nof attention_0  0.033163172
## 16 Cruel to\nothers=16 Cruel to\nothers_1                        0.029269520
## 20 Destroys own\nthings=20 Destroys own\nthings_1                0.207881796
## 90 Swearing=90 Swearing_1                                       -0.146726934
## 19 Demands a lot\nof attention=19 Demands a lot\nof attention_1 -0.008226851
## 16 Cruel to\nothers=16 Cruel to\nothers_2                       -0.120167888
## 57 Attacks\nothers=57 Attacks\nothers_0                         -0.002819163
## 104 Loud=104 Loud_2                                             -0.069827306
## 57 Attacks\nothers=57 Attacks\nothers_1                         -0.101076061
## 94 Teases a lot=94 Teases a lot_1                               -0.459887041
## 26 Lacks guilt=26 Lacks guilt_1                                 -0.157615049
## 43 Lying=43 Lying_1                                             -0.442256420
## 68 Screams a lot=68 Screams a lot_2                             -0.101472478
## 106 Vandalism=106 Vandalism_2                                   -3.440150563
## 39 Bad friends=39 Bad friends_0                                 -0.545930608
## 15 Cruel to\nanimals=15 Cruel to\nanimals_2                     -0.336474757
## 37 Gets in\nfights=37 Gets in\nfights_1                         -0.127330657
## 21 Destroys\nothers things=21 Destroys\nothers things_2         -0.549996165
## 101 Truant=101 Truant_1                                         -0.486716665
## 88 Sulks=88 Sulks_2                                             -0.153892460
## 97 Threatens\nothers=97 Threatens\nothers_2                     -1.136389800
## 20 Destroys own\nthings=20 Destroys own\nthings_2               -0.348984535
## 89 Suspicious=89 Suspicious_2                                   -0.453005142
## 81 Steals at\nhome=81 Steals at\nhome_0                         -1.526391693
## 106 Vandalism=106 Vandalism_0                                   -2.519094330
##                                                                       p.value
## 106 Vandalism=106 Vandalism_1                                   4.317920e-307
## 81 Steals at\nhome=81 Steals at\nhome_2                         5.652693e-240
## 39 Bad friends=39 Bad friends_2                                  6.258597e-73
## 43 Lying=43 Lying_2                                              1.853079e-65
## 94 Teases a lot=94 Teases a lot_2                                2.548804e-31
## 90 Swearing=90 Swearing_2                                        5.000527e-16
## 26 Lacks guilt=26 Lacks guilt_2                                  2.452633e-15
## 21 Destroys\nothers things=21 Destroys\nothers things_1          5.118699e-09
## 101 Truant=101 Truant_0                                          1.254435e-07
## 21 Destroys\nothers things=21 Destroys\nothers things_0          1.728106e-06
## 37 Gets in\nfights=37 Gets in\nfights_0                          1.123385e-05
## 15 Cruel to\nanimals=15 Cruel to\nanimals_0                      2.969523e-04
## 68 Screams a lot=68 Screams a lot_1                              8.137698e-04
## 16 Cruel to\nothers=16 Cruel to\nothers_0                        1.360426e-03
## 89 Suspicious=89 Suspicious_1                                    3.550392e-03
## 97 Threatens\nothers=97 Threatens\nothers_0                      3.774499e-03
## 19 Demands a lot\nof attention=19 Demands a lot\nof attention_0  4.997852e-03
## 16 Cruel to\nothers=16 Cruel to\nothers_1                        5.356966e-03
## 20 Destroys own\nthings=20 Destroys own\nthings_1                5.518724e-03
## 90 Swearing=90 Swearing_1                                        4.885115e-02
## 19 Demands a lot\nof attention=19 Demands a lot\nof attention_1  4.566352e-02
## 16 Cruel to\nothers=16 Cruel to\nothers_2                        3.581145e-02
## 57 Attacks\nothers=57 Attacks\nothers_0                          2.025372e-02
## 104 Loud=104 Loud_2                                              6.604811e-03
## 57 Attacks\nothers=57 Attacks\nothers_1                          5.603401e-03
## 94 Teases a lot=94 Teases a lot_1                                4.319140e-03
## 26 Lacks guilt=26 Lacks guilt_1                                  1.409335e-03
## 43 Lying=43 Lying_1                                              4.247619e-05
## 68 Screams a lot=68 Screams a lot_2                              1.215667e-05
## 106 Vandalism=106 Vandalism_2                                    4.605955e-06
## 39 Bad friends=39 Bad friends_0                                  5.191060e-07
## 15 Cruel to\nanimals=15 Cruel to\nanimals_2                      4.813494e-07
## 37 Gets in\nfights=37 Gets in\nfights_1                          1.714524e-07
## 21 Destroys\nothers things=21 Destroys\nothers things_2          9.214758e-08
## 101 Truant=101 Truant_1                                          5.570701e-08
## 88 Sulks=88 Sulks_2                                              6.657623e-09
## 97 Threatens\nothers=97 Threatens\nothers_2                      7.715519e-10
## 20 Destroys own\nthings=20 Destroys own\nthings_2                8.548866e-11
## 89 Suspicious=89 Suspicious_2                                    6.523869e-14
## 81 Steals at\nhome=81 Steals at\nhome_0                          2.182949e-23
## 106 Vandalism=106 Vandalism_0                                    2.431166e-43
## 
## attr(,"class")
## [1] "condes" "list"  
## 
## $`Dim 5`
## $quali
##                                         R2      p.value
## 88 Sulks                       0.184983468 1.640588e-60
## 37 Gets in\nfights             0.172676968 3.938693e-56
## 89 Suspicious                  0.146643082 4.476825e-47
## 87 Mood changes                0.144598242 2.241317e-46
## 86 Stubborn,\nsullen           0.134894744 4.440212e-43
## 20 Destroys own\nthings        0.118042320 1.933451e-37
## 21 Destroys\nothers things     0.116881011 4.687457e-37
## 97 Threatens\nothers           0.091332540 1.015194e-28
## 23 Disobedient\nat school      0.090743539 1.570162e-28
## 104 Loud                       0.072769842 8.264569e-23
## 3 Argues a lot                 0.065960952 1.136755e-20
## 22 Disobedient\nat home        0.061302586 3.234024e-19
## 90 Swearing                    0.060877311 4.386620e-19
## 95 Temper                      0.059254372 1.402156e-18
## 57 Attacks\nothers             0.045729873 2.083551e-14
## 94 Teases a lot                0.040127137 1.071098e-12
## 16 Cruel to\nothers            0.036451784 1.402275e-11
## 68 Screams a lot               0.034456539 5.642083e-11
## 106 Vandalism                  0.024924611 4.194740e-08
## 19 Demands a lot\nof attention 0.023232189 1.347631e-07
## 81 Steals at\nhome             0.016612909 1.269377e-05
## 39 Bad friends                 0.010347472 9.117650e-04
## 26 Lacks guilt                 0.005141941 3.113403e-02
## 
## $category
##                                                                     Estimate
## 37 Gets in\nfights=37 Gets in\nfights_2                          0.608708900
## 21 Destroys\nothers things=21 Destroys\nothers things_1          0.176316987
## 20 Destroys own\nthings=20 Destroys own\nthings_1                0.235097589
## 89 Suspicious=89 Suspicious_0                                    0.364392003
## 97 Threatens\nothers=97 Threatens\nothers_1                      0.772176558
## 88 Sulks=88 Sulks_0                                              0.215219597
## 89 Suspicious=89 Suspicious_1                                    0.105924859
## 23 Disobedient\nat school=23 Disobedient\nat school_2            0.421491646
## 87 Mood changes=87 Mood changes_0                                0.163072669
## 22 Disobedient\nat home=22 Disobedient\nat home_2                0.207253977
## 90 Swearing=90 Swearing_2                                        0.252507823
## 57 Attacks\nothers=57 Attacks\nothers_1                          0.131760416
## 104 Loud=104 Loud_2                                              0.202353927
## 94 Teases a lot=94 Teases a lot_1                                0.059587478
## 95 Temper=95 Temper_0                                            0.105667348
## 68 Screams a lot=68 Screams a lot_0                              0.076915830
## 86 Stubborn,\nsullen=86 Stubborn,\nsullen_0                      0.120420476
## 3 Argues a lot=3 Argues a lot_0                                  0.008519972
## 16 Cruel to\nothers=16 Cruel to\nothers_2                        0.314353344
## 3 Argues a lot=3 Argues a lot_2                                  0.094816020
## 39 Bad friends=39 Bad friends_1                                  0.075502570
## 88 Sulks=88 Sulks_1                                              0.125168819
## 19 Demands a lot\nof attention=19 Demands a lot\nof attention_2  0.063079479
## 81 Steals at\nhome=81 Steals at\nhome_1                          0.291583753
## 19 Demands a lot\nof attention=19 Demands a lot\nof attention_0  0.004911165
## 95 Temper=95 Temper_1                                            0.037516893
## 26 Lacks guilt=26 Lacks guilt_1                                  0.037888714
## 15 Cruel to\nanimals=15 Cruel to\nanimals_1                      0.080631419
## 106 Vandalism=106 Vandalism_0                                    0.392380527
## 97 Threatens\nothers=97 Threatens\nothers_2                     -0.640928120
## 15 Cruel to\nanimals=15 Cruel to\nanimals_0                     -0.024078955
## 20 Destroys own\nthings=20 Destroys own\nthings_2               -0.174676106
## 104 Loud=104 Loud_0                                             -0.039919140
## 26 Lacks guilt=26 Lacks guilt_0                                 -0.010541589
## 22 Disobedient\nat home=22 Disobedient\nat home_0               -0.072179541
## 39 Bad friends=39 Bad friends_0                                 -0.029399603
## 81 Steals at\nhome=81 Steals at\nhome_2                         -0.410718084
## 37 Gets in\nfights=37 Gets in\nfights_1                         -0.248731713
## 68 Screams a lot=68 Screams a lot_2                             -0.061171319
## 16 Cruel to\nothers=16 Cruel to\nothers_1                       -0.117492150
## 90 Swearing=90 Swearing_1                                       -0.164232648
## 68 Screams a lot=68 Screams a lot_1                             -0.015744511
## 16 Cruel to\nothers=16 Cruel to\nothers_0                       -0.196861194
## 19 Demands a lot\nof attention=19 Demands a lot\nof attention_1 -0.067990643
## 22 Disobedient\nat home=22 Disobedient\nat home_1               -0.135074436
## 106 Vandalism=106 Vandalism_1                                   -1.065768643
## 23 Disobedient\nat school=23 Disobedient\nat school_1           -0.141777434
## 104 Loud=104 Loud_1                                             -0.162434787
## 94 Teases a lot=94 Teases a lot_0                               -0.126876362
## 57 Attacks\nothers=57 Attacks\nothers_0                         -0.120980637
## 95 Temper=95 Temper_2                                           -0.143184241
## 23 Disobedient\nat school=23 Disobedient\nat school_0           -0.279714212
## 3 Argues a lot=3 Argues a lot_1                                 -0.103335993
## 37 Gets in\nfights=37 Gets in\nfights_0                         -0.359977187
## 97 Threatens\nothers=97 Threatens\nothers_0                     -0.131248439
## 89 Suspicious=89 Suspicious_2                                   -0.470316862
## 20 Destroys own\nthings=20 Destroys own\nthings_0               -0.060421483
## 21 Destroys\nothers things=21 Destroys\nothers things_0         -0.125535529
## 86 Stubborn,\nsullen=86 Stubborn,\nsullen_2                     -0.214631079
## 87 Mood changes=87 Mood changes_2                               -0.273781103
## 88 Sulks=88 Sulks_2                                             -0.340388416
##                                                                      p.value
## 37 Gets in\nfights=37 Gets in\nfights_2                         1.405940e-52
## 21 Destroys\nothers things=21 Destroys\nothers things_1         3.356058e-38
## 20 Destroys own\nthings=20 Destroys own\nthings_1               5.183976e-38
## 89 Suspicious=89 Suspicious_0                                   3.075716e-36
## 97 Threatens\nothers=97 Threatens\nothers_1                     5.712300e-29
## 88 Sulks=88 Sulks_0                                             1.996985e-23
## 89 Suspicious=89 Suspicious_1                                   4.679212e-21
## 23 Disobedient\nat school=23 Disobedient\nat school_2           2.372609e-18
## 87 Mood changes=87 Mood changes_0                               1.149353e-16
## 22 Disobedient\nat home=22 Disobedient\nat home_2               5.675369e-16
## 90 Swearing=90 Swearing_2                                       6.642136e-16
## 57 Attacks\nothers=57 Attacks\nothers_1                         3.946813e-15
## 104 Loud=104 Loud_2                                             5.629924e-15
## 94 Teases a lot=94 Teases a lot_1                               8.630143e-13
## 95 Temper=95 Temper_0                                           4.176629e-12
## 68 Screams a lot=68 Screams a lot_0                             1.878252e-11
## 86 Stubborn,\nsullen=86 Stubborn,\nsullen_0                     6.449802e-10
## 3 Argues a lot=3 Argues a lot_0                                 7.867879e-09
## 16 Cruel to\nothers=16 Cruel to\nothers_2                       8.798005e-09
## 3 Argues a lot=3 Argues a lot_2                                 1.159649e-08
## 39 Bad friends=39 Bad friends_1                                 1.859052e-04
## 88 Sulks=88 Sulks_1                                             2.005492e-04
## 19 Demands a lot\nof attention=19 Demands a lot\nof attention_2 1.273196e-03
## 81 Steals at\nhome=81 Steals at\nhome_1                         1.518091e-03
## 19 Demands a lot\nof attention=19 Demands a lot\nof attention_0 3.211006e-03
## 95 Temper=95 Temper_1                                           4.428675e-03
## 26 Lacks guilt=26 Lacks guilt_1                                 9.155521e-03
## 15 Cruel to\nanimals=15 Cruel to\nanimals_1                     1.718960e-02
## 106 Vandalism=106 Vandalism_0                                   4.739971e-02
## 97 Threatens\nothers=97 Threatens\nothers_2                     4.766544e-02
## 15 Cruel to\nanimals=15 Cruel to\nanimals_0                     4.142973e-02
## 20 Destroys own\nthings=20 Destroys own\nthings_2               3.985444e-02
## 104 Loud=104 Loud_0                                             3.043519e-02
## 26 Lacks guilt=26 Lacks guilt_0                                 2.629111e-02
## 22 Disobedient\nat home=22 Disobedient\nat home_0               1.152547e-02
## 39 Bad friends=39 Bad friends_0                                 4.762838e-04
## 81 Steals at\nhome=81 Steals at\nhome_2                         3.968997e-04
## 37 Gets in\nfights=37 Gets in\nfights_1                         1.201865e-04
## 68 Screams a lot=68 Screams a lot_2                             1.185602e-04
## 16 Cruel to\nothers=16 Cruel to\nothers_1                       1.152336e-04
## 90 Swearing=90 Swearing_1                                       7.850274e-07
## 68 Screams a lot=68 Screams a lot_1                             5.567196e-07
## 16 Cruel to\nothers=16 Cruel to\nothers_0                       4.047242e-07
## 19 Demands a lot\nof attention=19 Demands a lot\nof attention_1 2.672569e-07
## 22 Disobedient\nat home=22 Disobedient\nat home_1               2.349161e-08
## 106 Vandalism=106 Vandalism_1                                   1.883070e-08
## 23 Disobedient\nat school=23 Disobedient\nat school_1           3.001979e-11
## 104 Loud=104 Loud_1                                             3.736218e-12
## 94 Teases a lot=94 Teases a lot_0                               1.122859e-13
## 57 Attacks\nothers=57 Attacks\nothers_0                         5.583588e-15
## 95 Temper=95 Temper_2                                           7.812962e-16
## 23 Disobedient\nat school=23 Disobedient\nat school_0           3.425413e-18
## 3 Argues a lot=3 Argues a lot_1                                 2.580379e-19
## 37 Gets in\nfights=37 Gets in\nfights_0                         5.502794e-20
## 97 Threatens\nothers=97 Threatens\nothers_0                     1.589371e-23
## 89 Suspicious=89 Suspicious_2                                   5.719780e-26
## 20 Destroys own\nthings=20 Destroys own\nthings_0               5.125435e-31
## 21 Destroys\nothers things=21 Destroys\nothers things_0         1.484306e-37
## 86 Stubborn,\nsullen=86 Stubborn,\nsullen_2                     1.398111e-43
## 87 Mood changes=87 Mood changes_2                               2.205426e-45
## 88 Sulks=88 Sulks_2                                             2.211317e-53
## 
## attr(,"class")
## [1] "condes" "list"  
## 
## $`Dim 6`
## $quali
##                                         R2      p.value
## 23 Disobedient\nat school      0.263230356 5.179053e-90
## 57 Attacks\nothers             0.135438849 2.907484e-43
## 16 Cruel to\nothers            0.130059002 1.890752e-41
## 88 Sulks                       0.096869617 1.659797e-30
## 37 Gets in\nfights             0.088330361 9.346135e-28
## 19 Demands a lot\nof attention 0.086865701 2.753109e-27
## 87 Mood changes                0.086638485 3.254939e-27
## 104 Loud                       0.084804596 1.255475e-26
## 89 Suspicious                  0.074812448 1.873452e-23
## 86 Stubborn,\nsullen           0.072504464 1.001982e-22
## 97 Threatens\nothers           0.054741280 3.512498e-17
## 39 Bad friends                 0.047130586 7.752992e-15
## 101 Truant                     0.043992740 7.085673e-14
## 95 Temper                      0.043557246 9.627073e-14
## 43 Lying                       0.040065431 1.118454e-12
## 15 Cruel to\nanimals           0.032990166 1.566713e-10
## 68 Screams a lot               0.031598812 4.123120e-10
## 22 Disobedient\nat home        0.029402595 1.893789e-09
## 81 Steals at\nhome             0.025531434 2.759006e-08
## 3 Argues a lot                 0.019569533 1.673036e-06
## 94 Teases a lot                0.011751081 3.507941e-04
## 106 Vandalism                  0.011311241 4.732694e-04
## 90 Swearing                    0.005168691 3.057570e-02
## 
## $category
##                                                                     Estimate
## 57 Attacks\nothers=57 Attacks\nothers_1                          0.408163831
## 37 Gets in\nfights=37 Gets in\nfights_1                          0.159932805
## 57 Attacks\nothers=57 Attacks\nothers_0                          0.034984937
## 23 Disobedient\nat school=23 Disobedient\nat school_1            0.507413375
## 89 Suspicious=89 Suspicious_2                                    0.466280330
## 86 Stubborn,\nsullen=86 Stubborn,\nsullen_0                      0.054443308
## 97 Threatens\nothers=97 Threatens\nothers_1                      0.438694465
## 16 Cruel to\nothers=16 Cruel to\nothers_1                        0.372431612
## 88 Sulks=88 Sulks_0                                              0.007560814
## 39 Bad friends=39 Bad friends_1                                  0.118747258
## 87 Mood changes=87 Mood changes_0                                0.029495580
## 43 Lying=43 Lying_1                                              0.008443142
## 101 Truant=101 Truant_2                                          1.222310488
## 68 Screams a lot=68 Screams a lot_0                              0.069997768
## 95 Temper=95 Temper_0                                            0.032998531
## 16 Cruel to\nothers=16 Cruel to\nothers_0                        0.223209424
## 15 Cruel to\nanimals=15 Cruel to\nanimals_1                      0.054416095
## 88 Sulks=88 Sulks_2                                              0.135240867
## 81 Steals at\nhome=81 Steals at\nhome_1                          0.042483099
## 3 Argues a lot=3 Argues a lot_0                                  0.042908328
## 23 Disobedient\nat school=23 Disobedient\nat school_0            0.344737295
## 104 Loud=104 Loud_1                                              0.145730454
## 87 Mood changes=87 Mood changes_2                                0.105035683
## 19 Demands a lot\nof attention=19 Demands a lot\nof attention_0  0.079580364
## 106 Vandalism=106 Vandalism_1                                    0.434895281
## 15 Cruel to\nanimals=15 Cruel to\nanimals_2                      0.127998366
## 43 Lying=43 Lying_2                                              0.097698865
## 19 Demands a lot\nof attention=19 Demands a lot\nof attention_1  0.091054734
## 81 Steals at\nhome=81 Steals at\nhome_2                          0.177459554
## 94 Teases a lot=94 Teases a lot_1                                0.132977690
## 22 Disobedient\nat home=22 Disobedient\nat home_0                0.081660832
## 90 Swearing=90 Swearing_1                                        0.010508342
## 20 Destroys own\nthings=20 Destroys own\nthings_2                0.104426726
## 26 Lacks guilt=26 Lacks guilt_1                                  0.015596567
## 26 Lacks guilt=26 Lacks guilt_0                                 -0.021119610
## 90 Swearing=90 Swearing_0                                       -0.029874849
## 94 Teases a lot=94 Teases a lot_2                               -0.203571391
## 106 Vandalism=106 Vandalism_0                                   -0.384840083
## 68 Screams a lot=68 Screams a lot_2                             -0.061607041
## 101 Truant=101 Truant_1                                         -0.922255678
## 68 Screams a lot=68 Screams a lot_1                             -0.008390727
## 3 Argues a lot=3 Argues a lot_1                                 -0.026726035
## 89 Suspicious=89 Suspicious_1                                   -0.293045769
## 57 Attacks\nothers=57 Attacks\nothers_2                         -0.443148768
## 81 Steals at\nhome=81 Steals at\nhome_0                         -0.219942653
## 22 Disobedient\nat home=22 Disobedient\nat home_2               -0.145180570
## 15 Cruel to\nanimals=15 Cruel to\nanimals_0                     -0.182414461
## 43 Lying=43 Lying_0                                             -0.106142007
## 95 Temper=95 Temper_1                                           -0.083526349
## 39 Bad friends=39 Bad friends_0                                 -0.089294227
## 97 Threatens\nothers=97 Threatens\nothers_0                     -0.221703084
## 86 Stubborn,\nsullen=86 Stubborn,\nsullen_1                     -0.082681136
## 104 Loud=104 Loud_2                                             -0.226550019
## 37 Gets in\nfights=37 Gets in\nfights_0                         -0.100886125
## 16 Cruel to\nothers=16 Cruel to\nothers_2                       -0.595641036
## 87 Mood changes=87 Mood changes_1                               -0.134531263
## 88 Sulks=88 Sulks_1                                             -0.142801681
## 19 Demands a lot\nof attention=19 Demands a lot\nof attention_2 -0.170635098
## 23 Disobedient\nat school=23 Disobedient\nat school_2           -0.852150670
##                                                                      p.value
## 57 Attacks\nothers=57 Attacks\nothers_1                         9.260561e-38
## 37 Gets in\nfights=37 Gets in\nfights_1                         8.402359e-29
## 57 Attacks\nothers=57 Attacks\nothers_0                         2.322430e-25
## 23 Disobedient\nat school=23 Disobedient\nat school_1           9.809154e-22
## 89 Suspicious=89 Suspicious_2                                   2.940306e-19
## 86 Stubborn,\nsullen=86 Stubborn,\nsullen_0                     8.650780e-19
## 97 Threatens\nothers=97 Threatens\nothers_1                     3.134048e-18
## 16 Cruel to\nothers=16 Cruel to\nothers_1                       4.153438e-18
## 88 Sulks=88 Sulks_0                                             1.057736e-16
## 39 Bad friends=39 Bad friends_1                                 9.982221e-16
## 87 Mood changes=87 Mood changes_0                               7.174748e-15
## 43 Lying=43 Lying_1                                             2.967631e-11
## 101 Truant=101 Truant_2                                         2.417941e-10
## 68 Screams a lot=68 Screams a lot_0                             2.707393e-10
## 95 Temper=95 Temper_0                                           5.731466e-10
## 16 Cruel to\nothers=16 Cruel to\nothers_0                       7.294971e-10
## 15 Cruel to\nanimals=15 Cruel to\nanimals_1                     7.468445e-09
## 88 Sulks=88 Sulks_2                                             2.142831e-07
## 81 Steals at\nhome=81 Steals at\nhome_1                         2.515544e-07
## 3 Argues a lot=3 Argues a lot_0                                 2.740328e-07
## 23 Disobedient\nat school=23 Disobedient\nat school_0           2.821536e-07
## 104 Loud=104 Loud_1                                             7.684025e-06
## 87 Mood changes=87 Mood changes_2                               1.013053e-04
## 19 Demands a lot\nof attention=19 Demands a lot\nof attention_0 4.012817e-04
## 106 Vandalism=106 Vandalism_1                                   7.080125e-04
## 15 Cruel to\nanimals=15 Cruel to\nanimals_2                     9.076154e-04
## 43 Lying=43 Lying_2                                             3.714839e-03
## 19 Demands a lot\nof attention=19 Demands a lot\nof attention_1 3.855411e-03
## 81 Steals at\nhome=81 Steals at\nhome_2                         4.966021e-03
## 94 Teases a lot=94 Teases a lot_1                               8.466527e-03
## 22 Disobedient\nat home=22 Disobedient\nat home_0               9.085795e-03
## 90 Swearing=90 Swearing_1                                       1.823838e-02
## 20 Destroys own\nthings=20 Destroys own\nthings_2               2.026995e-02
## 26 Lacks guilt=26 Lacks guilt_1                                 4.026636e-02
## 26 Lacks guilt=26 Lacks guilt_0                                 3.225094e-02
## 90 Swearing=90 Swearing_0                                       8.447564e-03
## 94 Teases a lot=94 Teases a lot_2                               2.324392e-03
## 106 Vandalism=106 Vandalism_0                                   3.816544e-04
## 68 Screams a lot=68 Screams a lot_2                             5.141944e-05
## 101 Truant=101 Truant_1                                         7.947877e-06
## 68 Screams a lot=68 Screams a lot_1                             7.761251e-06
## 3 Argues a lot=3 Argues a lot_1                                 5.013702e-06
## 89 Suspicious=89 Suspicious_1                                   5.617279e-07
## 57 Attacks\nothers=57 Attacks\nothers_2                         5.264622e-08
## 81 Steals at\nhome=81 Steals at\nhome_0                         5.568097e-09
## 22 Disobedient\nat home=22 Disobedient\nat home_2               5.937805e-10
## 15 Cruel to\nanimals=15 Cruel to\nanimals_0                     2.402193e-11
## 43 Lying=43 Lying_0                                             3.235112e-13
## 95 Temper=95 Temper_1                                           1.201609e-14
## 39 Bad friends=39 Bad friends_0                                 3.817280e-15
## 97 Threatens\nothers=97 Threatens\nothers_0                     1.099986e-16
## 86 Stubborn,\nsullen=86 Stubborn,\nsullen_1                     1.536493e-23
## 104 Loud=104 Loud_2                                             8.258545e-25
## 37 Gets in\nfights=37 Gets in\nfights_0                         3.499955e-26
## 16 Cruel to\nothers=16 Cruel to\nothers_2                       2.697536e-26
## 87 Mood changes=87 Mood changes_1                               9.060175e-27
## 88 Sulks=88 Sulks_1                                             3.211998e-28
## 19 Demands a lot\nof attention=19 Demands a lot\nof attention_2 2.682672e-28
## 23 Disobedient\nat school=23 Disobedient\nat school_2           1.854352e-69
## 
## attr(,"class")
## [1] "condes" "list"  
## 
## $`Dim 7`
## $quali
##                                         R2       p.value
## 97 Threatens\nothers           0.294840629 7.894787e-103
## 43 Lying                       0.190188388  2.200047e-62
## 101 Truant                     0.189502578  3.889128e-62
## 89 Suspicious                  0.136212886  1.591194e-43
## 20 Destroys own\nthings        0.078266573  1.511445e-24
## 94 Teases a lot                0.072740227  8.444136e-23
## 81 Steals at\nhome             0.071351168  2.312442e-22
## 21 Destroys\nothers things     0.053403782  9.096760e-17
## 39 Bad friends                 0.051603704  3.267146e-16
## 57 Attacks\nothers             0.041839980  3.219621e-13
## 16 Cruel to\nothers            0.033959657  7.976497e-11
## 90 Swearing                    0.026545837  1.368787e-08
## 88 Sulks                       0.021787245  3.644394e-07
## 19 Demands a lot\nof attention 0.019195088  2.163274e-06
## 37 Gets in\nfights             0.017238957  8.269105e-06
## 68 Screams a lot               0.016976612  9.896266e-06
## 15 Cruel to\nanimals           0.014978601  3.880904e-05
## 87 Mood changes                0.013159624  1.343292e-04
## 22 Disobedient\nat home        0.009026206  2.237898e-03
## 23 Disobedient\nat school      0.008404013  3.414239e-03
## 95 Temper                      0.006063215  1.668969e-02
## 26 Lacks guilt                 0.005198658  2.996209e-02
## 
## $category
##                                                                     Estimate
## 101 Truant=101 Truant_1                                          1.434107723
## 43 Lying=43 Lying_2                                              0.641727509
## 89 Suspicious=89 Suspicious_2                                    0.629538569
## 97 Threatens\nothers=97 Threatens\nothers_2                      2.415451198
## 94 Teases a lot=94 Teases a lot_1                                0.235424660
## 94 Teases a lot=94 Teases a lot_0                                0.019576007
## 81 Steals at\nhome=81 Steals at\nhome_0                          0.039542017
## 57 Attacks\nothers=57 Attacks\nothers_2                          0.441576742
## 16 Cruel to\nothers=16 Cruel to\nothers_1                        0.023922013
## 21 Destroys\nothers things=21 Destroys\nothers things_1          0.342421942
## 88 Sulks=88 Sulks_2                                              0.077128951
## 87 Mood changes=87 Mood changes_0                                0.046469471
## 22 Disobedient\nat home=22 Disobedient\nat home_2                0.072018973
## 37 Gets in\nfights=37 Gets in\nfights_2                          0.142681498
## 39 Bad friends=39 Bad friends_1                                  0.246521369
## 21 Destroys\nothers things=21 Destroys\nothers things_0          0.257889192
## 81 Steals at\nhome=81 Steals at\nhome_2                          0.379691894
## 95 Temper=95 Temper_0                                            0.030117604
## 26 Lacks guilt=26 Lacks guilt_1                                  0.037168762
## 20 Destroys own\nthings=20 Destroys own\nthings_1                0.236249830
## 19 Demands a lot\nof attention=19 Demands a lot\nof attention_1  0.048469136
## 23 Disobedient\nat school=23 Disobedient\nat school_2            0.102957275
## 90 Swearing=90 Swearing_1                                        0.092019268
## 86 Stubborn,\nsullen=86 Stubborn,\nsullen_0                      0.017656852
## 22 Disobedient\nat home=22 Disobedient\nat home_1               -0.045723451
## 26 Lacks guilt=26 Lacks guilt_0                                 -0.004146536
## 3 Argues a lot=3 Argues a lot_1                                 -0.021202926
## 95 Temper=95 Temper_2                                           -0.031675959
## 23 Disobedient\nat school=23 Disobedient\nat school_1           -0.030843412
## 43 Lying=43 Lying_1                                             -0.333180502
## 37 Gets in\nfights=37 Gets in\nfights_0                         -0.029129824
## 23 Disobedient\nat school=23 Disobedient\nat school_0           -0.072113863
## 88 Sulks=88 Sulks_0                                             -0.008099026
## 94 Teases a lot=94 Teases a lot_2                               -0.255000667
## 87 Mood changes=87 Mood changes_2                               -0.064484974
## 37 Gets in\nfights=37 Gets in\nfights_1                         -0.113551674
## 89 Suspicious=89 Suspicious_0                                   -0.317298872
## 15 Cruel to\nanimals=15 Cruel to\nanimals_2                     -0.267443419
## 88 Sulks=88 Sulks_1                                             -0.069029925
## 68 Screams a lot=68 Screams a lot_2                             -0.083556681
## 19 Demands a lot\nof attention=19 Demands a lot\nof attention_2 -0.074979502
## 90 Swearing=90 Swearing_2                                       -0.155844308
## 16 Cruel to\nothers=16 Cruel to\nothers_0                       -0.089286907
## 21 Destroys\nothers things=21 Destroys\nothers things_2         -0.600311134
## 39 Bad friends=39 Bad friends_2                                 -0.425038148
## 81 Steals at\nhome=81 Steals at\nhome_1                         -0.419233911
## 20 Destroys own\nthings=20 Destroys own\nthings_2               -0.427082508
## 97 Threatens\nothers=97 Threatens\nothers_0                     -0.666798730
## 101 Truant=101 Truant_0                                         -0.697284335
## 97 Threatens\nothers=97 Threatens\nothers_1                     -1.748652467
##                                                                      p.value
## 101 Truant=101 Truant_1                                         1.774639e-63
## 43 Lying=43 Lying_2                                             4.134850e-63
## 89 Suspicious=89 Suspicious_2                                   8.896123e-45
## 97 Threatens\nothers=97 Threatens\nothers_2                     1.096862e-43
## 94 Teases a lot=94 Teases a lot_1                               1.356631e-21
## 94 Teases a lot=94 Teases a lot_0                               5.719065e-17
## 81 Steals at\nhome=81 Steals at\nhome_0                         4.794455e-16
## 57 Attacks\nothers=57 Attacks\nothers_2                         1.231242e-13
## 16 Cruel to\nothers=16 Cruel to\nothers_1                       7.515358e-11
## 21 Destroys\nothers things=21 Destroys\nothers things_1         3.894557e-05
## 88 Sulks=88 Sulks_2                                             1.133118e-03
## 87 Mood changes=87 Mood changes_0                               1.366100e-03
## 22 Disobedient\nat home=22 Disobedient\nat home_2               1.627224e-03
## 37 Gets in\nfights=37 Gets in\nfights_2                         2.023764e-03
## 39 Bad friends=39 Bad friends_1                                 3.670700e-03
## 21 Destroys\nothers things=21 Destroys\nothers things_0         7.209492e-03
## 81 Steals at\nhome=81 Steals at\nhome_2                         9.023813e-03
## 95 Temper=95 Temper_0                                           9.251137e-03
## 26 Lacks guilt=26 Lacks guilt_1                                 1.105910e-02
## 20 Destroys own\nthings=20 Destroys own\nthings_1               1.231764e-02
## 19 Demands a lot\nof attention=19 Demands a lot\nof attention_1 1.302299e-02
## 23 Disobedient\nat school=23 Disobedient\nat school_2           1.559428e-02
## 90 Swearing=90 Swearing_1                                       2.398047e-02
## 86 Stubborn,\nsullen=86 Stubborn,\nsullen_0                     4.827340e-02
## 22 Disobedient\nat home=22 Disobedient\nat home_1               4.586498e-02
## 26 Lacks guilt=26 Lacks guilt_0                                 4.400445e-02
## 3 Argues a lot=3 Argues a lot_1                                 4.234613e-02
## 95 Temper=95 Temper_2                                           3.403657e-02
## 23 Disobedient\nat school=23 Disobedient\nat school_1           2.447597e-02
## 43 Lying=43 Lying_1                                             1.974411e-02
## 37 Gets in\nfights=37 Gets in\nfights_0                         1.446936e-02
## 23 Disobedient\nat school=23 Disobedient\nat school_0           5.125010e-03
## 88 Sulks=88 Sulks_0                                             1.484040e-03
## 94 Teases a lot=94 Teases a lot_2                               8.258386e-04
## 87 Mood changes=87 Mood changes_2                               1.457329e-04
## 37 Gets in\nfights=37 Gets in\nfights_1                         1.369424e-04
## 89 Suspicious=89 Suspicious_0                                   1.587476e-05
## 15 Cruel to\nanimals=15 Cruel to\nanimals_2                     1.270511e-05
## 88 Sulks=88 Sulks_1                                             2.152480e-06
## 68 Screams a lot=68 Screams a lot_2                             2.071378e-06
## 19 Demands a lot\nof attention=19 Demands a lot\nof attention_2 1.169446e-06
## 90 Swearing=90 Swearing_2                                       9.571489e-09
## 16 Cruel to\nothers=16 Cruel to\nothers_0                       1.049744e-11
## 21 Destroys\nothers things=21 Destroys\nothers things_2         3.612881e-14
## 39 Bad friends=39 Bad friends_2                                 1.583308e-15
## 81 Steals at\nhome=81 Steals at\nhome_1                         6.098980e-22
## 20 Destroys own\nthings=20 Destroys own\nthings_2               1.430403e-24
## 97 Threatens\nothers=97 Threatens\nothers_0                     1.071522e-24
## 101 Truant=101 Truant_0                                         3.641722e-46
## 97 Threatens\nothers=97 Threatens\nothers_1                     6.886148e-54
## 
## attr(,"class")
## [1] "condes" "list"  
## 
## $`Dim 8`
## $quali
##                                         R2      p.value
## 20 Destroys own\nthings        0.229377070 6.992254e-77
## 37 Gets in\nfights             0.177408424 8.298612e-58
## 21 Destroys\nothers things     0.150806447 1.665333e-48
## 81 Steals at\nhome             0.106826569 9.545319e-34
## 57 Attacks\nothers             0.106656610 1.084931e-33
## 22 Disobedient\nat home        0.096848888 1.685635e-30
## 39 Bad friends                 0.094274054 1.145123e-29
## 16 Cruel to\nothers            0.058728049 2.043026e-18
## 15 Cruel to\nanimals           0.046337296 1.357373e-14
## 104 Loud                       0.033859094 8.555324e-11
## 94 Teases a lot                0.024456427 5.794425e-08
## 43 Lying                       0.024383580 6.093054e-08
## 3 Argues a lot                 0.024353054 6.222718e-08
## 87 Mood changes                0.022601452 2.080881e-07
## 101 Truant                     0.019771179 1.456753e-06
## 19 Demands a lot\nof attention 0.014495481 5.398228e-05
## 95 Temper                      0.013777769 8.811147e-05
## 88 Sulks                       0.013276441 1.240423e-04
## 89 Suspicious                  0.012979107 1.519255e-04
## 23 Disobedient\nat school      0.008390660 3.445322e-03
## 90 Swearing                    0.007933406 4.698679e-03
## 26 Lacks guilt                 0.005930569 1.825794e-02
## 
## $category
##                                                                     Estimate
## 37 Gets in\nfights=37 Gets in\nfights_2                          0.585981853
## 21 Destroys\nothers things=21 Destroys\nothers things_0          0.016537843
## 81 Steals at\nhome=81 Steals at\nhome_0                          0.305693522
## 57 Attacks\nothers=57 Attacks\nothers_2                          0.586909213
## 22 Disobedient\nat home=22 Disobedient\nat home_2                0.236686950
## 39 Bad friends=39 Bad friends_2                                  0.514694874
## 15 Cruel to\nanimals=15 Cruel to\nanimals_2                      0.390402345
## 20 Destroys own\nthings=20 Destroys own\nthings_2                0.365784290
## 16 Cruel to\nothers=16 Cruel to\nothers_1                        0.220900156
## 94 Teases a lot=94 Teases a lot_1                                0.135317051
## 104 Loud=104 Loud_1                                              0.096078632
## 3 Argues a lot=3 Argues a lot_2                                  0.076521997
## 16 Cruel to\nothers=16 Cruel to\nothers_0                        0.114041823
## 94 Teases a lot=94 Teases a lot_0                                0.015065932
## 43 Lying=43 Lying_2                                              0.167989459
## 87 Mood changes=87 Mood changes_1                                0.069137107
## 88 Sulks=88 Sulks_1                                              0.052212549
## 19 Demands a lot\nof attention=19 Demands a lot\nof attention_1  0.046061591
## 95 Temper=95 Temper_1                                            0.050801150
## 101 Truant=101 Truant_2                                          0.623373249
## 23 Disobedient\nat school=23 Disobedient\nat school_0            0.072696007
## 90 Swearing=90 Swearing_1                                        0.002102103
## 88 Sulks=88 Sulks_0                                              0.006037865
## 21 Destroys\nothers things=21 Destroys\nothers things_2          0.257043438
## 104 Loud=104 Loud_0                                              0.012606244
## 26 Lacks guilt=26 Lacks guilt_1                                  0.043927102
## 106 Vandalism=106 Vandalism_2                                    0.252135859
## 23 Disobedient\nat school=23 Disobedient\nat school_1            0.036293170
## 94 Teases a lot=94 Teases a lot_2                               -0.150382983
## 101 Truant=101 Truant_0                                         -0.038377209
## 15 Cruel to\nanimals=15 Cruel to\nanimals_1                     -0.145849354
## 88 Sulks=88 Sulks_2                                             -0.058250414
## 23 Disobedient\nat school=23 Disobedient\nat school_2           -0.108989177
## 81 Steals at\nhome=81 Steals at\nhome_2                         -0.062094376
## 39 Bad friends=39 Bad friends_0                                 -0.183392970
## 90 Swearing=90 Swearing_0                                       -0.039607541
## 43 Lying=43 Lying_0                                             -0.051756021
## 95 Temper=95 Temper_2                                           -0.060010611
## 19 Demands a lot\nof attention=19 Demands a lot\nof attention_2 -0.054865935
## 3 Argues a lot=3 Argues a lot_1                                 -0.052553500
## 89 Suspicious=89 Suspicious_2                                   -0.181974117
## 37 Gets in\nfights=37 Gets in\nfights_1                         -0.333206055
## 101 Truant=101 Truant_1                                         -0.584996040
## 87 Mood changes=87 Mood changes_2                               -0.085982472
## 43 Lying=43 Lying_1                                             -0.116233438
## 57 Attacks\nothers=57 Attacks\nothers_1                         -0.229208163
## 104 Loud=104 Loud_2                                             -0.108684877
## 15 Cruel to\nanimals=15 Cruel to\nanimals_0                     -0.244552991
## 22 Disobedient\nat home=22 Disobedient\nat home_1               -0.143497138
## 16 Cruel to\nothers=16 Cruel to\nothers_2                       -0.334941979
## 39 Bad friends=39 Bad friends_1                                 -0.331301904
## 57 Attacks\nothers=57 Attacks\nothers_0                         -0.357701050
## 81 Steals at\nhome=81 Steals at\nhome_1                         -0.243599146
## 20 Destroys own\nthings=20 Destroys own\nthings_0               -0.018313618
## 21 Destroys\nothers things=21 Destroys\nothers things_1         -0.273581281
## 20 Destroys own\nthings=20 Destroys own\nthings_1               -0.347470673
##                                                                      p.value
## 37 Gets in\nfights=37 Gets in\nfights_2                         1.196534e-55
## 21 Destroys\nothers things=21 Destroys\nothers things_0         3.045335e-44
## 81 Steals at\nhome=81 Steals at\nhome_0                         1.593032e-34
## 57 Attacks\nothers=57 Attacks\nothers_2                         9.867746e-30
## 22 Disobedient\nat home=22 Disobedient\nat home_2               8.498805e-28
## 39 Bad friends=39 Bad friends_2                                 6.289628e-22
## 15 Cruel to\nanimals=15 Cruel to\nanimals_2                     4.925646e-14
## 20 Destroys own\nthings=20 Destroys own\nthings_2               2.551137e-12
## 16 Cruel to\nothers=16 Cruel to\nothers_1                       3.541658e-11
## 94 Teases a lot=94 Teases a lot_1                               5.747568e-08
## 104 Loud=104 Loud_1                                             8.561468e-08
## 3 Argues a lot=3 Argues a lot_2                                 1.178424e-07
## 16 Cruel to\nothers=16 Cruel to\nothers_0                       3.240707e-07
## 94 Teases a lot=94 Teases a lot_0                               2.209007e-06
## 43 Lying=43 Lying_2                                             6.281246e-05
## 87 Mood changes=87 Mood changes_1                               6.350512e-05
## 88 Sulks=88 Sulks_1                                             2.183248e-04
## 19 Demands a lot\nof attention=19 Demands a lot\nof attention_1 7.674503e-04
## 95 Temper=95 Temper_1                                           8.817701e-04
## 101 Truant=101 Truant_2                                         3.018450e-03
## 23 Disobedient\nat school=23 Disobedient\nat school_0           8.522443e-03
## 90 Swearing=90 Swearing_1                                       9.754644e-03
## 88 Sulks=88 Sulks_0                                             1.295002e-02
## 21 Destroys\nothers things=21 Destroys\nothers things_2         1.645267e-02
## 104 Loud=104 Loud_0                                             1.725782e-02
## 26 Lacks guilt=26 Lacks guilt_1                                 2.178364e-02
## 106 Vandalism=106 Vandalism_2                                   3.716269e-02
## 23 Disobedient\nat school=23 Disobedient\nat school_1           4.224653e-02
## 94 Teases a lot=94 Teases a lot_2                               3.878603e-02
## 101 Truant=101 Truant_0                                         2.897973e-02
## 15 Cruel to\nanimals=15 Cruel to\nanimals_1                     1.160495e-02
## 88 Sulks=88 Sulks_2                                             1.131306e-02
## 23 Disobedient\nat school=23 Disobedient\nat school_2           9.228254e-03
## 81 Steals at\nhome=81 Steals at\nhome_2                         5.529444e-03
## 39 Bad friends=39 Bad friends_0                                 1.701525e-03
## 90 Swearing=90 Swearing_0                                       1.614574e-03
## 43 Lying=43 Lying_0                                             1.528179e-03
## 95 Temper=95 Temper_2                                           1.337949e-03
## 19 Demands a lot\nof attention=19 Demands a lot\nof attention_2 4.925485e-04
## 3 Argues a lot=3 Argues a lot_1                                 2.255473e-04
## 89 Suspicious=89 Suspicious_2                                   3.481384e-05
## 37 Gets in\nfights=37 Gets in\nfights_1                         3.379824e-05
## 101 Truant=101 Truant_1                                         2.201737e-05
## 87 Mood changes=87 Mood changes_2                               1.745262e-05
## 43 Lying=43 Lying_1                                             1.529930e-05
## 57 Attacks\nothers=57 Attacks\nothers_1                         8.820882e-06
## 104 Loud=104 Loud_2                                             3.354460e-06
## 15 Cruel to\nanimals=15 Cruel to\nanimals_0                     7.009791e-08
## 22 Disobedient\nat home=22 Disobedient\nat home_1               9.690705e-09
## 16 Cruel to\nothers=16 Cruel to\nothers_2                       3.676675e-10
## 39 Bad friends=39 Bad friends_1                                 1.850735e-10
## 57 Attacks\nothers=57 Attacks\nothers_0                         7.103389e-15
## 81 Steals at\nhome=81 Steals at\nhome_1                         5.584916e-33
## 20 Destroys own\nthings=20 Destroys own\nthings_0               5.843970e-44
## 21 Destroys\nothers things=21 Destroys\nothers things_1         1.295327e-48
## 20 Destroys own\nthings=20 Destroys own\nthings_1               5.536308e-67
## 
## attr(,"class")
## [1] "condes" "list"  
## 
## $`Dim 9`
## $quali
##                                         R2       p.value
## 57 Attacks\nothers             0.291306521 2.283116e-101
## 19 Demands a lot\nof attention 0.141923377  1.832506e-45
## 90 Swearing                    0.139800401  9.666868e-45
## 39 Bad friends                 0.101183019  6.618565e-32
## 81 Steals at\nhome             0.079722900  5.214686e-25
## 97 Threatens\nothers           0.058992326  1.691220e-18
## 20 Destroys own\nthings        0.047250609  7.122813e-15
## 101 Truant                     0.042797392  1.642883e-13
## 43 Lying                       0.041942136  2.996685e-13
## 21 Destroys\nothers things     0.041188279  5.087847e-13
## 22 Disobedient\nat home        0.038409807  3.566869e-12
## 26 Lacks guilt                 0.028314798  4.024587e-09
## 15 Cruel to\nanimals           0.027907835  5.334689e-09
## 87 Mood changes                0.027782133  5.819712e-09
## 89 Suspicious                  0.026453043  1.459474e-08
## 86 Stubborn,\nsullen           0.024096672  7.426355e-08
## 88 Sulks                       0.019361953  1.929219e-06
## 37 Gets in\nfights             0.018681129  3.077735e-06
## 68 Screams a lot               0.017688351  6.078203e-06
## 3 Argues a lot                 0.014364510  5.903263e-05
## 23 Disobedient\nat school      0.010375752  8.943980e-04
## 104 Loud                       0.005397195  2.619596e-02
## 16 Cruel to\nothers            0.005162305  3.070806e-02
## 
## $category
##                                                                      Estimate
## 57 Attacks\nothers=57 Attacks\nothers_2                          1.0936372223
## 19 Demands a lot\nof attention=19 Demands a lot\nof attention_1  0.1420371188
## 39 Bad friends=39 Bad friends_2                                  0.5194156418
## 81 Steals at\nhome=81 Steals at\nhome_1                          0.4832328959
## 97 Threatens\nothers=97 Threatens\nothers_0                      0.4652812997
## 101 Truant=101 Truant_0                                          0.5757937366
## 43 Lying=43 Lying_1                                              0.0828700310
## 22 Disobedient\nat home=22 Disobedient\nat home_2                0.1540892546
## 81 Steals at\nhome=81 Steals at\nhome_0                          0.0561129845
## 21 Destroys\nothers things=21 Destroys\nothers things_1          0.0113347945
## 26 Lacks guilt=26 Lacks guilt_2                                  0.1525359298
## 87 Mood changes=87 Mood changes_0                                0.0511835302
## 89 Suspicious=89 Suspicious_0                                    0.1136858272
## 20 Destroys own\nthings=20 Destroys own\nthings_2                0.1679659866
## 90 Swearing=90 Swearing_1                                        0.1994265335
## 88 Sulks=88 Sulks_0                                              0.0110664512
## 86 Stubborn,\nsullen=86 Stubborn,\nsullen_0                      0.0144829976
## 68 Screams a lot=68 Screams a lot_2                              0.0707960621
## 19 Demands a lot\nof attention=19 Demands a lot\nof attention_0  0.0262388805
## 3 Argues a lot=3 Argues a lot_1                                  0.0404086609
## 15 Cruel to\nanimals=15 Cruel to\nanimals_2                      0.2175810252
## 16 Cruel to\nothers=16 Cruel to\nothers_1                        0.0209002991
## 86 Stubborn,\nsullen=86 Stubborn,\nsullen_2                      0.0366318258
## 104 Loud=104 Loud_2                                              0.0482487993
## 21 Destroys\nothers things=21 Destroys\nothers things_2          0.1220656537
## 95 Temper=95 Temper_0                                            0.0167444603
## 94 Teases a lot=94 Teases a lot_2                                0.1163879188
## 95 Temper=95 Temper_1                                           -0.0140549333
## 22 Disobedient\nat home=22 Disobedient\nat home_0               -0.0814804495
## 16 Cruel to\nothers=16 Cruel to\nothers_0                       -0.0225776117
## 89 Suspicious=89 Suspicious_2                                   -0.1130098229
## 23 Disobedient\nat school=23 Disobedient\nat school_1           -0.0003472391
## 68 Screams a lot=68 Screams a lot_1                             -0.0557607020
## 39 Bad friends=39 Bad friends_0                                 -0.1817188825
## 23 Disobedient\nat school=23 Disobedient\nat school_0           -0.0572415482
## 97 Threatens\nothers=97 Threatens\nothers_2                     -0.3650282751
## 15 Cruel to\nanimals=15 Cruel to\nanimals_0                     -0.0068467912
## 3 Argues a lot=3 Argues a lot_2                                 -0.0562580851
## 101 Truant=101 Truant_2                                         -0.3428415662
## 81 Steals at\nhome=81 Steals at\nhome_2                         -0.5393458804
## 89 Suspicious=89 Suspicious_1                                   -0.0006760044
## 88 Sulks=88 Sulks_1                                             -0.0536735256
## 37 Gets in\nfights=37 Gets in\nfights_2                         -0.1868115243
## 15 Cruel to\nanimals=15 Cruel to\nanimals_1                     -0.2107342340
## 87 Mood changes=87 Mood changes_1                               -0.0352341855
## 86 Stubborn,\nsullen=86 Stubborn,\nsullen_1                     -0.0511148234
## 20 Destroys own\nthings=20 Destroys own\nthings_1               -0.0223463986
## 101 Truant=101 Truant_1                                         -0.2329521704
## 39 Bad friends=39 Bad friends_1                                 -0.3376967593
## 57 Attacks\nothers=57 Attacks\nothers_1                         -0.6373537513
## 43 Lying=43 Lying_0                                             -0.0326423413
## 20 Destroys own\nthings=20 Destroys own\nthings_0               -0.1456195880
## 21 Destroys\nothers things=21 Destroys\nothers things_0         -0.1334004482
## 97 Threatens\nothers=97 Threatens\nothers_1                     -0.1002530245
## 19 Demands a lot\nof attention=19 Demands a lot\nof attention_2 -0.1682759993
## 90 Swearing=90 Swearing_2                                       -0.3431848956
##                                                                      p.value
## 57 Attacks\nothers=57 Attacks\nothers_2                         9.540099e-90
## 19 Demands a lot\nof attention=19 Demands a lot\nof attention_1 3.538726e-27
## 39 Bad friends=39 Bad friends_2                                 8.080074e-23
## 81 Steals at\nhome=81 Steals at\nhome_1                         5.563362e-21
## 97 Threatens\nothers=97 Threatens\nothers_0                     2.938942e-19
## 101 Truant=101 Truant_0                                         1.835117e-14
## 43 Lying=43 Lying_1                                             3.219870e-14
## 22 Disobedient\nat home=22 Disobedient\nat home_2               5.012576e-13
## 81 Steals at\nhome=81 Steals at\nhome_0                         8.563744e-13
## 21 Destroys\nothers things=21 Destroys\nothers things_1         8.973690e-13
## 26 Lacks guilt=26 Lacks guilt_2                                 6.011879e-10
## 87 Mood changes=87 Mood changes_0                               9.259074e-10
## 89 Suspicious=89 Suspicious_0                                   7.291166e-09
## 20 Destroys own\nthings=20 Destroys own\nthings_2               3.170232e-07
## 90 Swearing=90 Swearing_1                                       1.572144e-06
## 88 Sulks=88 Sulks_0                                             3.747899e-05
## 86 Stubborn,\nsullen=86 Stubborn,\nsullen_0                     5.537838e-05
## 68 Screams a lot=68 Screams a lot_2                             9.013116e-05
## 19 Demands a lot\nof attention=19 Demands a lot\nof attention_0 2.483637e-04
## 3 Argues a lot=3 Argues a lot_1                                 2.495588e-03
## 15 Cruel to\nanimals=15 Cruel to\nanimals_2                     5.829814e-03
## 16 Cruel to\nothers=16 Cruel to\nothers_1                       8.813182e-03
## 86 Stubborn,\nsullen=86 Stubborn,\nsullen_2                     1.431010e-02
## 104 Loud=104 Loud_2                                             2.595200e-02
## 21 Destroys\nothers things=21 Destroys\nothers things_2         2.848538e-02
## 95 Temper=95 Temper_0                                           2.936407e-02
## 94 Teases a lot=94 Teases a lot_2                               4.168510e-02
## 95 Temper=95 Temper_1                                           3.783430e-02
## 22 Disobedient\nat home=22 Disobedient\nat home_0               2.670069e-02
## 16 Cruel to\nothers=16 Cruel to\nothers_0                       8.605001e-03
## 89 Suspicious=89 Suspicious_2                                   1.050222e-03
## 23 Disobedient\nat school=23 Disobedient\nat school_1           8.833611e-04
## 68 Screams a lot=68 Screams a lot_1                             4.797693e-04
## 39 Bad friends=39 Bad friends_0                                 4.651819e-04
## 23 Disobedient\nat school=23 Disobedient\nat school_0           2.632415e-04
## 97 Threatens\nothers=97 Threatens\nothers_2                     1.714967e-04
## 15 Cruel to\nanimals=15 Cruel to\nanimals_0                     8.152576e-05
## 3 Argues a lot=3 Argues a lot_2                                 7.807364e-05
## 101 Truant=101 Truant_2                                         3.007724e-05
## 81 Steals at\nhome=81 Steals at\nhome_2                         2.007049e-06
## 89 Suspicious=89 Suspicious_1                                   8.261359e-07
## 88 Sulks=88 Sulks_1                                             5.053645e-07
## 37 Gets in\nfights=37 Gets in\nfights_2                         4.926369e-07
## 15 Cruel to\nanimals=15 Cruel to\nanimals_1                     3.008550e-08
## 87 Mood changes=87 Mood changes_1                               2.121035e-08
## 86 Stubborn,\nsullen=86 Stubborn,\nsullen_1                     1.777749e-08
## 20 Destroys own\nthings=20 Destroys own\nthings_1               1.901068e-09
## 101 Truant=101 Truant_1                                         1.625302e-10
## 39 Bad friends=39 Bad friends_1                                 8.869659e-12
## 57 Attacks\nothers=57 Attacks\nothers_1                         3.061012e-12
## 43 Lying=43 Lying_0                                             4.730312e-13
## 20 Destroys own\nthings=20 Destroys own\nthings_0               1.041775e-13
## 21 Destroys\nothers things=21 Destroys\nothers things_0         8.824444e-14
## 97 Threatens\nothers=97 Threatens\nothers_1                     2.888958e-16
## 19 Demands a lot\nof attention=19 Demands a lot\nof attention_2 1.833509e-28
## 90 Swearing=90 Swearing_2                                       1.540435e-42
## 
## attr(,"class")
## [1] "condes" "list"  
## 
## $`Dim 10`
## $quali
##                                         R2      p.value
## 23 Disobedient\nat school      0.194285885 7.240928e-64
## 101 Truant                     0.143704033 4.527762e-46
## 20 Destroys own\nthings        0.116286488 7.372868e-37
## 21 Destroys\nothers things     0.094674919 8.500872e-30
## 89 Suspicious                  0.086984783 2.521765e-27
## 43 Lying                       0.067849892 2.910583e-21
## 26 Lacks guilt                 0.065984022 1.118015e-20
## 88 Sulks                       0.055085779 2.748370e-17
## 39 Bad friends                 0.047842984 4.686718e-15
## 37 Gets in\nfights             0.043424701 1.056808e-13
## 15 Cruel to\nanimals           0.038741034 2.828731e-12
## 16 Cruel to\nothers            0.032188809 2.735901e-10
## 87 Mood changes                0.031013615 6.191491e-10
## 95 Temper                      0.025984363 2.017764e-08
## 22 Disobedient\nat home        0.025254723 3.339928e-08
## 104 Loud                       0.023199241 1.378574e-07
## 97 Threatens\nothers           0.019491455 1.765146e-06
## 94 Teases a lot                0.019079707 2.341494e-06
## 106 Vandalism                  0.011760662 3.485127e-04
## 81 Steals at\nhome             0.007191972 7.768436e-03
## 3 Argues a lot                 0.006629133 1.137588e-02
## 68 Screams a lot               0.005843307 1.936902e-02
## 86 Stubborn,\nsullen           0.005407086 2.602123e-02
## 19 Demands a lot\nof attention 0.004871806 3.737557e-02
## 
## $category
##                                                                    Estimate
## 23 Disobedient\nat school=23 Disobedient\nat school_2            0.67379438
## 101 Truant=101 Truant_2                                          2.05303478
## 20 Destroys own\nthings=20 Destroys own\nthings_0                0.17077239
## 21 Destroys\nothers things=21 Destroys\nothers things_0          0.12748458
## 43 Lying=43 Lying_1                                              0.13949827
## 26 Lacks guilt=26 Lacks guilt_1                                  0.12274736
## 43 Lying=43 Lying_0                                              0.00493366
## 39 Bad friends=39 Bad friends_1                                  0.04251719
## 89 Suspicious=89 Suspicious_2                                    0.36176368
## 15 Cruel to\nanimals=15 Cruel to\nanimals_0                      0.16770382
## 16 Cruel to\nothers=16 Cruel to\nothers_2                        0.31349497
## 87 Mood changes=87 Mood changes_0                                0.04917263
## 88 Sulks=88 Sulks_2                                              0.13078286
## 37 Gets in\nfights=37 Gets in\nfights_1                          0.16973477
## 104 Loud=104 Loud_2                                              0.10473439
## 22 Disobedient\nat home=22 Disobedient\nat home_2                0.10437368
## 94 Teases a lot=94 Teases a lot_2                                0.25576069
## 97 Threatens\nothers=97 Threatens\nothers_0                      0.37427992
## 106 Vandalism=106 Vandalism_1                                    0.59400236
## 37 Gets in\nfights=37 Gets in\nfights_0                          0.05936676
## 81 Steals at\nhome=81 Steals at\nhome_2                          0.23100018
## 39 Bad friends=39 Bad friends_2                                  0.08728653
## 68 Screams a lot=68 Screams a lot_1                              0.02949504
## 19 Demands a lot\nof attention=19 Demands a lot\nof attention_2  0.03548679
## 97 Threatens\nothers=97 Threatens\nothers_1                      0.20543074
## 3 Argues a lot=3 Argues a lot_2                                  0.03145893
## 95 Temper=95 Temper_0                                            0.04485506
## 22 Disobedient\nat home=22 Disobedient\nat home_1               -0.03508491
## 104 Loud=104 Loud_1                                             -0.06509436
## 23 Disobedient\nat school=23 Disobedient\nat school_1           -0.31395665
## 94 Teases a lot=94 Teases a lot_1                               -0.15292568
## 3 Argues a lot=3 Argues a lot_0                                 -0.02773966
## 86 Stubborn,\nsullen=86 Stubborn,\nsullen_2                     -0.03369465
## 101 Truant=101 Truant_0                                         -0.74993936
## 43 Lying=43 Lying_2                                             -0.14443193
## 15 Cruel to\nanimals=15 Cruel to\nanimals_2                     -0.10413277
## 26 Lacks guilt=26 Lacks guilt_2                                 -0.11017424
## 22 Disobedient\nat home=22 Disobedient\nat home_0               -0.06928877
## 88 Sulks=88 Sulks_0                                             -0.02628169
## 97 Threatens\nothers=97 Threatens\nothers_2                     -0.57971066
## 101 Truant=101 Truant_1                                         -1.30309542
## 20 Destroys own\nthings=20 Destroys own\nthings_2               -0.11598523
## 89 Suspicious=89 Suspicious_0                                   -0.09187777
## 37 Gets in\nfights=37 Gets in\nfights_2                         -0.22910152
## 95 Temper=95 Temper_2                                           -0.09416271
## 87 Mood changes=87 Mood changes_1                               -0.04079974
## 23 Disobedient\nat school=23 Disobedient\nat school_0           -0.35983774
## 15 Cruel to\nanimals=15 Cruel to\nanimals_1                     -0.06357105
## 88 Sulks=88 Sulks_1                                             -0.10450117
## 26 Lacks guilt=26 Lacks guilt_0                                 -0.01257313
## 39 Bad friends=39 Bad friends_0                                 -0.12980372
## 89 Suspicious=89 Suspicious_1                                   -0.26988591
## 21 Destroys\nothers things=21 Destroys\nothers things_1         -0.09284994
## 20 Destroys own\nthings=20 Destroys own\nthings_1               -0.05478716
##                                                                      p.value
## 23 Disobedient\nat school=23 Disobedient\nat school_2           4.467196e-63
## 101 Truant=101 Truant_2                                         3.036511e-42
## 20 Destroys own\nthings=20 Destroys own\nthings_0               7.970375e-38
## 21 Destroys\nothers things=21 Destroys\nothers things_0         6.667640e-31
## 43 Lying=43 Lying_1                                             1.188525e-20
## 26 Lacks guilt=26 Lacks guilt_1                                 5.226231e-20
## 43 Lying=43 Lying_0                                             1.740064e-16
## 39 Bad friends=39 Bad friends_1                                 6.241548e-14
## 89 Suspicious=89 Suspicious_2                                   2.567661e-13
## 15 Cruel to\nanimals=15 Cruel to\nanimals_0                     3.367887e-13
## 16 Cruel to\nothers=16 Cruel to\nothers_2                       3.588655e-11
## 87 Mood changes=87 Mood changes_0                               1.510718e-10
## 88 Sulks=88 Sulks_2                                             5.974860e-10
## 37 Gets in\nfights=37 Gets in\nfights_1                         4.752758e-08
## 104 Loud=104 Loud_2                                             6.991402e-08
## 22 Disobedient\nat home=22 Disobedient\nat home_2               3.387601e-07
## 94 Teases a lot=94 Teases a lot_2                               6.454482e-06
## 97 Threatens\nothers=97 Threatens\nothers_0                     1.869493e-04
## 106 Vandalism=106 Vandalism_1                                   3.792689e-04
## 37 Gets in\nfights=37 Gets in\nfights_0                         2.182719e-03
## 81 Steals at\nhome=81 Steals at\nhome_2                         2.878924e-03
## 39 Bad friends=39 Bad friends_2                                 3.894027e-03
## 68 Screams a lot=68 Screams a lot_1                             7.599814e-03
## 19 Demands a lot\nof attention=19 Demands a lot\nof attention_2 1.144177e-02
## 97 Threatens\nothers=97 Threatens\nothers_1                     1.275468e-02
## 3 Argues a lot=3 Argues a lot_2                                 2.512867e-02
## 95 Temper=95 Temper_0                                           4.132442e-02
## 22 Disobedient\nat home=22 Disobedient\nat home_1               4.456779e-02
## 104 Loud=104 Loud_1                                             3.832652e-02
## 23 Disobedient\nat school=23 Disobedient\nat school_1           2.671062e-02
## 94 Teases a lot=94 Teases a lot_1                               1.460073e-02
## 3 Argues a lot=3 Argues a lot_0                                 1.209979e-02
## 86 Stubborn,\nsullen=86 Stubborn,\nsullen_2                     1.200844e-02
## 101 Truant=101 Truant_0                                         7.259956e-03
## 43 Lying=43 Lying_2                                             1.548733e-03
## 15 Cruel to\nanimals=15 Cruel to\nanimals_2                     9.718027e-04
## 26 Lacks guilt=26 Lacks guilt_2                                 6.010094e-04
## 22 Disobedient\nat home=22 Disobedient\nat home_0               1.009493e-04
## 88 Sulks=88 Sulks_0                                             8.057012e-05
## 97 Threatens\nothers=97 Threatens\nothers_2                     7.144098e-06
## 101 Truant=101 Truant_1                                         5.939868e-06
## 20 Destroys own\nthings=20 Destroys own\nthings_2               3.565021e-06
## 89 Suspicious=89 Suspicious_0                                   3.773843e-08
## 37 Gets in\nfights=37 Gets in\nfights_2                         2.243706e-08
## 95 Temper=95 Temper_2                                           2.765079e-09
## 87 Mood changes=87 Mood changes_1                               1.043440e-09
## 23 Disobedient\nat school=23 Disobedient\nat school_0           8.423664e-10
## 15 Cruel to\nanimals=15 Cruel to\nanimals_1                     1.209783e-10
## 88 Sulks=88 Sulks_1                                             8.041547e-12
## 26 Lacks guilt=26 Lacks guilt_0                                 6.653597e-13
## 39 Bad friends=39 Bad friends_0                                 5.434447e-16
## 89 Suspicious=89 Suspicious_1                                   8.045518e-17
## 21 Destroys\nothers things=21 Destroys\nothers things_1         2.050187e-30
## 20 Destroys own\nthings=20 Destroys own\nthings_1               6.098026e-32
## 
## attr(,"class")
## [1] "condes" "list"
```

Write MCA results to file

```
mca_coord <- get_mca_ind(mca_pheno)$coord
write.csv(mca_coord, params$cbcl_mca_coord_outfile)
save(mca_pheno, file = params$cbcl_mca_outfile)
```

```
sessionInfo()
```

```
## R version 4.1.2 (2021-11-01)
## Platform: x86_64-pc-linux-gnu (64-bit)
## Running under: Debian GNU/Linux bookworm/sid
## 
## Matrix products: default
## BLAS:   /usr/lib/x86_64-linux-gnu/openblas-pthread/libblas.so.3
## LAPACK: /usr/lib/x86_64-linux-gnu/openblas-pthread/libopenblasp-r0.3.19.so
## 
## locale:
##  [1] LC_CTYPE=en_US.UTF-8       LC_NUMERIC=C              
##  [3] LC_TIME=en_US.UTF-8        LC_COLLATE=en_US.UTF-8    
##  [5] LC_MONETARY=en_US.UTF-8    LC_MESSAGES=en_US.UTF-8   
##  [7] LC_PAPER=en_US.UTF-8       LC_NAME=C                 
##  [9] LC_ADDRESS=C               LC_TELEPHONE=C            
## [11] LC_MEASUREMENT=en_US.UTF-8 LC_IDENTIFICATION=C       
## 
## attached base packages:
## [1] stats     graphics  grDevices utils     datasets  methods   base     
## 
## other attached packages:
##  [1] factoextra_1.0.7    FactoMineR_2.4      ggplot2_3.3.5      
##  [4] stringr_1.4.0       reshape2_1.4.4      missForest_1.4     
##  [7] itertools_0.1-3     iterators_1.0.13    foreach_1.5.2      
## [10] randomForest_4.6-14
## 
## loaded via a namespace (and not attached):
##  [1] ggrepel_0.9.1        Rcpp_1.0.8           lattice_0.20-45     
##  [4] tidyr_1.2.0          digest_0.6.29        utf8_1.2.2          
##  [7] R6_2.5.1             plyr_1.8.6           backports_1.4.1     
## [10] evaluate_0.14        highr_0.9            pillar_1.7.0        
## [13] rlang_1.0.0          car_3.0-12           jquerylib_0.1.4     
## [16] DT_0.20              rmarkdown_2.11       labeling_0.4.2      
## [19] htmlwidgets_1.5.4    munsell_0.5.0        broom_0.7.12        
## [22] compiler_4.1.2       xfun_0.29            pkgconfig_2.0.3     
## [25] htmltools_0.5.2      flashClust_1.01-2    tidyselect_1.1.1    
## [28] tibble_3.1.6         codetools_0.2-18     fansi_1.0.2         
## [31] crayon_1.4.2         dplyr_1.0.8          withr_2.4.3         
## [34] ggpubr_0.4.0         MASS_7.3-55          leaps_3.1           
## [37] grid_4.1.2           jsonlite_1.7.3       gtable_0.3.0        
## [40] lifecycle_1.0.1      magrittr_2.0.2       scales_1.1.1        
## [43] carData_3.0-5        cli_3.1.1            stringi_1.7.6       
## [46] farver_2.1.0         ggsignif_0.6.3       scatterplot3d_0.3-41
## [49] bslib_0.3.1          ellipsis_0.3.2       generics_0.1.2      
## [52] vctrs_0.3.8          tools_4.1.2          glue_1.6.1          
## [55] purrr_0.3.4          abind_1.4-5          parallel_4.1.2      
## [58] fastmap_1.1.0        yaml_2.2.2           colorspace_2.0-2    
## [61] cluster_2.1.2        rstatix_0.7.0        knitr_1.37          
## [64] sass_0.4.0
```

---

1. Radboud University Medical Center,↩︎
